# A 4D transcriptomic map for the evolution of multiple sclerosis-like lesions in the marmoset brain

**DOI:** 10.1101/2023.09.25.559371

**Authors:** Jing-Ping Lin, Alexis Brake, Maxime Donadieu, Amanda Lee, Riki Kawaguchi, Pascal Sati, Daniel H. Geschwind, Steven Jacobson, Dorothy P. Schafer, Daniel S. Reich

## Abstract

Single-time-point histopathological studies on postmortem multiple sclerosis (MS) tissue fail to capture lesion evolution dynamics, posing challenges for therapy development targeting development and repair of focal inflammatory demyelination. To close this gap, we studied experimental autoimmune encephalitis (EAE) in the common marmoset, the most faithful animal model of these processes. Using MRI-informed RNA profiling, we analyzed ∼600,000 single-nucleus and ∼55,000 spatial transcriptomes, comparing them against EAE inoculation status, longitudinal radiological signals, and histopathological features. We categorized 5 groups of microenvironments pertinent to neural function, immune and glial responses, tissue destruction and repair, and regulatory network at brain borders. Exploring perilesional microenvironment diversity, we uncovered central roles of EAE-associated astrocytes, oligodendrocyte precursor cells, and ependyma in lesion formation and resolution. We pinpointed imaging and molecular features capturing the pathological trajectory of WM, offering potential for assessing treatment outcomes using marmoset as a platform.

**One sentence summary:** A cross-modality study to identify the spatiotemporal-based diversity of primate brain cells during white matter inflammatory demyelination to inform lesion detection, stratification, and management in multiple sclerosis.

## Main

Multiple sclerosis (MS) is a complex disease characterized by focal inflammation and loss of myelin in the central nervous system (CNS). While the underlying cause of MS is unclear, the interplay of inappropriate immune response and eventual failure to adequately repair myelin are important mechanisms (1). Despite considerable success in controlling peripherally derived inflammation with MS disease-modifying therapies (2), much more must be understood about the cellular dynamics of lesion progression, especially in acute and subacute phases, to develop treatments that facilitate timely remyelination.

Although experimental autoimmune encephalomyelitis (EAE) in the mouse has provided important insights into CNS myelin-directed inflammation, most of the current pathophysiological understanding of MS comes from studying postmortem human tissue or, in rare fulminant presentations, brain biopsies. However, a single time point, especially at the end of life, cannot capture the signaling profiles of lesion growth and resolution. To close this gap, we employed a clinically relevant model to study the initiation of and reaction to MS-like lesions. Relative to rodents, common marmosets (*Callithrix jacchus*) have high genetic, physiological, and immunological similarities to humans (3). Marmoset EAE recapitulates aspects of MS lesion evolution substantially better than mouse EAE (4), allowing the development of clinically transferable methods to monitor and predict lesion outcomes for treatment assessment.

Structural magnetic resonance imaging (MRI) is noninvasive and can sensitively monitor the spatiotemporal changes within MS lesions (5). However, it is not sufficiently specific to discern the cellular and molecular diversity that accounts for lesion heterogeneity. To bridge this gap, we performed a cross-modality study, joining longitudinal MRI, histopathology, spatial transcriptome mapping, and single-nucleus RNA profiling to dissect global and local signaling in lesion evolution. Leveraging the strength of each approach, we here summarize the sequence of radiological and biological events, nominating candidates for lesion stratification, molecular MRI, and treatment evaluation using the marmoset EAE platform.

### ▪ Model: Marmoset EAE recapitulates the formation of white matter lesions in MS

The hallmark of MS is multifocal, inflammatory loss of myelin in white matter (WM), gray matter (GM), spinal cord, and optic nerve (6). MRI biomarkers, such as gadolinium (Gd) enhancement (7), leptomeningeal enhancement (8), the central vein sign (9,10), and paramagnetic rim lesions (11) have been used to characterize the pathological course and predict clinical outcomes of MS (12). Mouse EAE, the most extensively investigated preclinical model of MS, typically does not form brain lesions (13); marmoset EAE, however, develops lesions in all aforementioned CNS areas (14–18). Moreover, relative to GM volume, WM in marmosets expanded evolutionarily more than 5 fold compared to the volumetric ratio in mice (19), allowing WM lesions in marmosets to be followed radiologically and histopathologically with detailed spatial resolution **(Fig1A).**

**Fig1.**
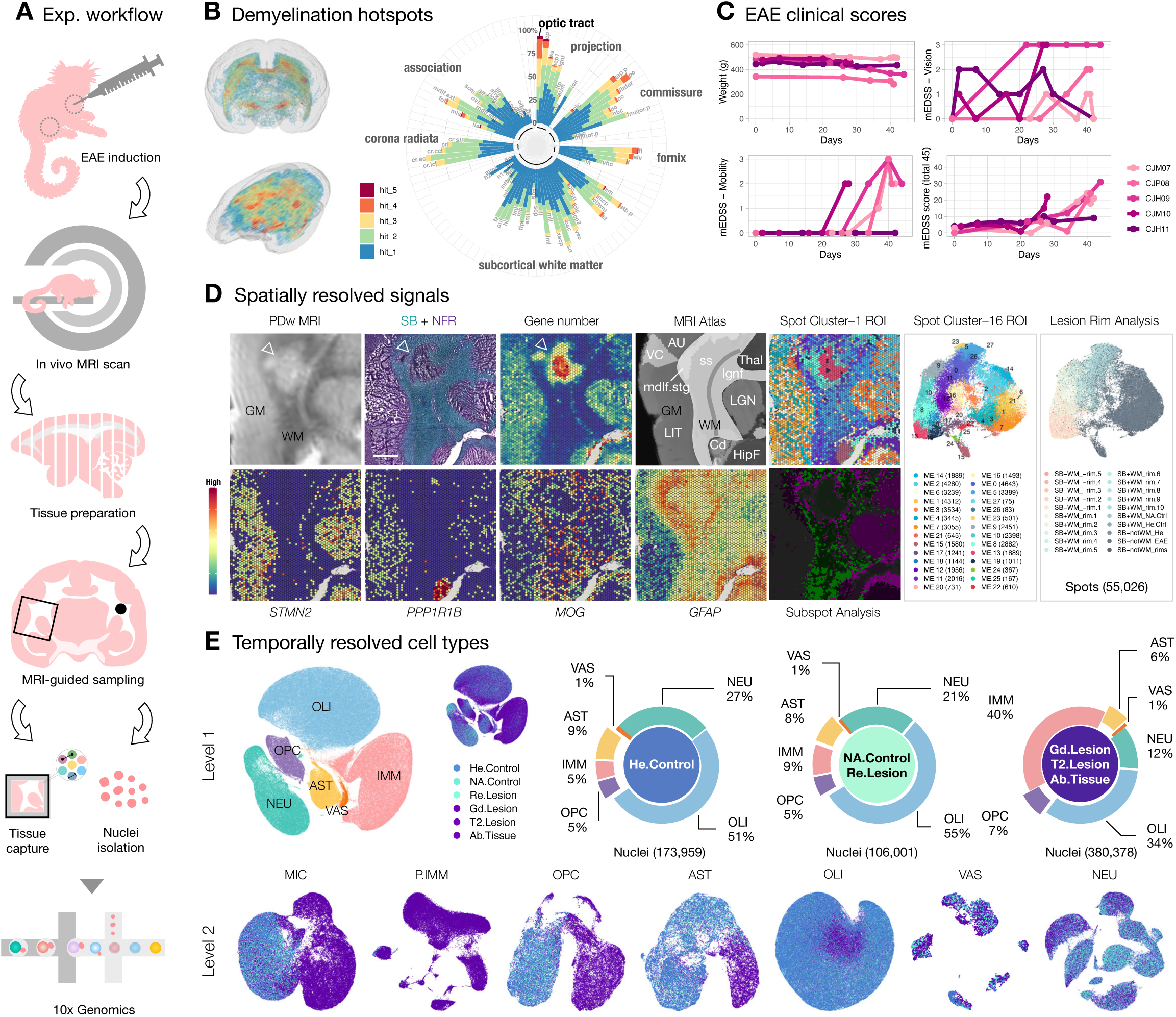
Marmoset experimental autoimmune encephalomyelitis (EAE) recapitulates the development and repair of multiple sclerosis-type white matter (WM) lesions and enables detailed mapping of spatiotemporal organization at the individual lesion level. (A) Experimental workflow for inducing EAE and preparing tissue samples for single-nucleus and spatial transcriptome analysis using the 10x Genomics platform. (B) Visual representation and quantification of lesion load in different WM tracts across 5 EAE animals. Higher lesion loads were observed in projection and commissural WM fibers, with the optic tract (opt) being particularly susceptible to demyelination. Refer to source data for the full list of abbreviations for WM tracts. (C) Line plots depict the changes in body weights and EAE clinical scores (range: 0–45) of the 5 EAE animals over time, measured using the expanded disability status scale developed specifically for marmosets (mEDSS). Subcategories of the mEDSS scores, such as vision and mobility, are summarized separately. (D) Overview of phenotypic characterization of a typical WM lesion (indicated by arrowheads) is presented, including proton density-weighted (PDw) magnetic resonance imaging (MRI), histological staining with Sudan black (SB) and nuclear fast red (NFR), spatial transcriptome profiling (gene number and selected markers), supervised anatomical indexing using an MRI atlas as reference, and unsupervised microenvironment (ME) classification with bioinformatic tools (spot and subspot level analysis). UMAP scatter plots summarizing a total of 55,026 spatial transcriptome spots were analyzed across 16 brain regions of interest (ROI) and colored based on transcriptome profile similarity (ME0–27) and spatial organization relative to demyelinated areas (lesion rim analysis). Refer to source data for the full list of abbreviations for brain regions. Scale bar = 1mm. (E) UMAP scatter plots illustrate level 1 (L1) and level 2 (L2) analyses of transcriptomes with single-nucleus resolution, color-coded by cell class identity or disease condition. Donut charts provide the relative proportions of cell classes in each disease group, including healthy (He) control, normal-appearing (NA) control, resolved (Re) lesion, gadolinium (Gd) positive lesion, T**_2_**-hyperintense (T2) MRI detected lesion, and abnormal (Ab) appearing tissue. In the L1 analysis, canonical cell-type markers were used to annotate central and peripheral immune cells (IMM), oligodendrocyte progenitor cells (OPC), oligodendrocytes (OLI), astrocytes (AST), vasculature and meningeal cells (VAS), and neurons (NEU). In the L2 analysis, the IMM cell class was further divided into microglia (MIC) and peripheral immune cells (P.IMM). Notably, as lesions developed, substantial cellular diversity was observed, particularly among glial and immune cells.

The use of antigens in suspension to induce autoimmune diseases dates back ∼75 years (20), and how different immunogens skew immune responses in the modeling of autoimmune diseases has been widely discussed (21,13). EAE induction in mice typically requires the mixing of myelin components in mycobacteria-containing mineral oil (complete Freund’s adjuvant, CFA) followed by a pertussis toxin booster for optimal reproducibility (21); however, the requirement for CFA serving as the “danger” signal to stimulate EAE appears to depend on species (21) and immune status (22,4). Marmosets have a human-like immune system, trained from early life onward through natural exposure to environmental pathogens. This fundamentally differs from laboratory rodent models, which are often bred and housed under specific pathogen-free (SPF) conditions. Marmosets can be sensitized by human myelin peptides in an adjuvant lacking microbial components (incomplete Freund’s adjuvant, IFA) to develop a disease with high neurological, radiological, and cellular/humoral immune similarities to that induced with CFA (23).

Inspired by these findings (23), we revamped our prior inoculation protocol by swapping CFA with IFA and human WM homogenate with myelin oligodendrocyte glycoprotein (hMOG) peptides emulsified in an enclosed connective device (**Methods**), which significantly reduced batch effects attributable to different WM donors and improved efficacy in inducing MS-like lesions across brain regions. In our hand, marmosets require only a single intradermal injection of hMOG/IFA to achieve the full spectrum of EAE. Across 5 adult marmosets immunized with hMOG/IFA emulsion, WM areas, including the optic tract (opt), visual projections, and commissural fibers that connect the brain hemispheres appear most vulnerable to inflammatory demyelination (**Fig1B**). The anatomical structures of these demyelination hotspots are largely periventricular, which phenocopies the prevalence of periventricular lesions found in MS (24,25). Lesions of marmoset EAE show Gd enhancement when the blood-brain barrier (BBB) is open (26,27), the central vein sign in developing lesions (28), and iron accumulation at the lesion (29) recapitulating these aspects of MS.

In addition to radiological and histopathological signatures, the first clinical sign of marmoset EAE, manifest within 1 week post injection (wpi), was a form of visual impairment and or muscle weakness, followed by mobility decline within 4 wpi; the total disability score peaked around 6 wpi (**Fig1C**). The expanded disability status scale (mEDSS) utilized here was specifically developed for marmosets (30) and captures alertness, spontaneous mobility, tremor, muscle tone, grip strength, sensory response, eye movement, pupillary reflex, vocalization, bladder function, and tail strength, allowing quantification of neurological impairment as the disease progress. The radiological and clinical presentation of visual abnormalities observed in marmoset EAE recapitulates that of many cases of MS, in which changes in visual acuity and optic neuritis are often found before other impairments (31–34), further corroborating that marmoset EAE is a relevant model in mimicking important aspects of MS.

### ▪ Study design: cross-modality imaging of MS-like WM inflammatory demyelination

The view of using a mechanism-driven framework to rate MS as a spectrum (1) over the traditional distinct clinical descriptors (relapsing-remitting, secondary progressive, and primary progressive, (35,36)) has guided the focus of our studying tissue damage at the individual lesion level. As overlapping pathological and compensatory pathways contribute to heterogeneity in lesion and clinical presentations (1), we categorized tissue by radiological features instead of by the onset or severity of neurological symptoms for each animal (**FigS1-2**). We then employed a cross-modality approach to map the cellular and molecular dynamics over time and space to appreciate the significance of focal and global signaling as lesions evolve (**Fig1D-E, FigS3**).

From 11 marmosets (**TableS1**), we derived the current transcriptomic map with spatial and single-nucleus resolution. To identify spatially enriched signals pertinent to lesion formation, we identified abnormal areas on proton density weighted (PDw) MRI (16) and confirmed demyelination by Sudan black (SB) lipid staining with nuclear fast red (NFR) contrast (**Methods**). We then profiled transcriptome at the region of interest (ROI) with 10x Visium (**TableS2**), anatomically annotated the ROI by MRI atlas indexing, and bioinformatically processed the data to categorize subregions (**Fig1D, FigS4-5**). The detection of *STMN2* (cortical and subcortical GM), *PPP1R1B* (caudate), *MOG* (WM), and *GFAP* (glial reactivity) transcripts robustly highlight well-characterized anatomical/pathological features of the tissue (37–40). Across 16 ROI, SB^-^-GM had more gene transcripts compared to SB^+^-WM, as expected, since cortical GM generally has higher cell density than WM. Compared to SB^-^-GM, SB^-^-WM had an even higher transcriptional complexity (**FigS3**), suggesting a hypercellular response to demyelinated WM, consistent with prior histopathological studies (41).

To understand the dynamics of these cells as lesions evolve, we utilized serial MRI to guide tissue sampling and estimate the age of the lesion retrospectively (**Methods**). We integrated and analyzed a total of 43 snRNA-seq libraries (**FigS1, TableS3**), with WM from healthy control (n = 13) and WM with T_2_-hyperintense demyelinated MRI feature from EAE animals (n = 14) being the most extensively sampled groups. We used additional categories, including normal-appearing (NA) WM from EAE animals (n = 2), Gd-enhancing demyelinating WM lesions (n = 2), and resolved WM lesions that no longer T_2_-hyperintense on the terminal MRI (n = 3) to group the rest of WM samples. In parallel, we included leukocortical T_2_-hyperintense lesions (n = 2) along with matching healthy (n = 2) and NA (n = 1) controls, and nearby abnormal-looking lateral geniculate nucleus (LGN) tissue on MRI (n = 2) along with matching healthy controls (n = 2), to explore tissue-specific or shared responses. We implemented a hierarchical workflow comparing across cell classes (Level 1, L1) and subclusters within a class (Level 2, L2) to better realize the importance of each signaling change (**Methods**, (42)). Overall, we found a remarkable expansion in the number and diversity of glial and immune cells as lesions develop (**Fig1E**).

In the following sections, we first present our findings in relation to the spatial organization of different approach-detectable changes from the most advanced to the earliest stages across modalities (**Fig2, FigS5-8**). We then describe the transition in cellular composition and signaling network from healthy to diseased state (**Fig3-4, FigS9-14**). We identify imaging features that capture the turning point when brain tissue yields to pathological attack (**Fig5**), which we envision would be clinically adaptable for lesion monitoring. By linking longitudinal MRI detectable changes that inform the disease history of the sampled areas, histology detectable changes that label molecular-based alterations, and RNA profiling detectable changes that manifests early distortion collectively, our analysis focuses on: (a) identifying glial-vascular-immune interactions; (b) comparing regional signaling networks within and across microenvironments; and (c) finding molecular and imaging features to advance identification and classification of MS lesions.

**Fig2.**
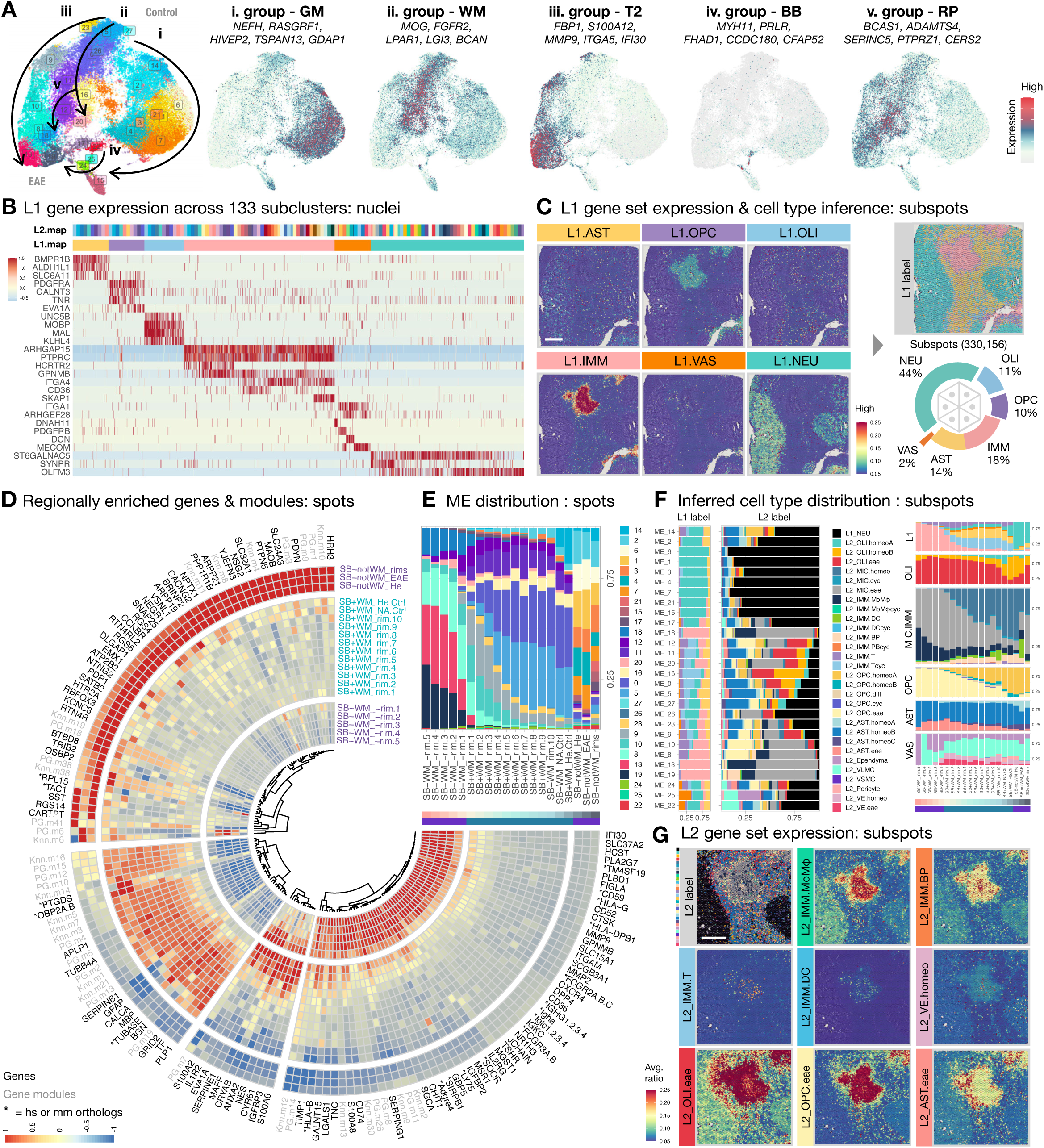
Spatially resolved pathways and cellular composition highlight the dynamics of in-situ and ex-situ tissue responses to pathological insults. (A) UMAP scatter plots are color-coded by microenvironment (ME) clustering and gene group expression, including GM (ME14, 2, 6, 1, 3, 4, 7, 21, 15, 17), WM (ME26, 27, 5, 0, 16, 20, 11), T2 lesion (T2, ME23, 9, 10, 8, 13, 19), brain borders (BB, ME22, 25, 24), and repair (RP, ME12, 18). The direction of arrows indicates increased prevalence in EAE. (B) Heatmap summarizes the z-scored expression of genes that clearly segregate major cell classes (L1.map; color code in **C**). Up to 200 nuclei were sampled from each group across 133 subclusters (L2.map; color code in **Fig3B**). (C) Spatial heatmaps of a region of interest (ROI) with representative white matter lesions showing the averaged expression of gene sets listed in (A) for each cell class across BayesSpace-enhanced subspots. By comparing the expression score across tested gene sets, L1 cell types were inferred for each subspot based on profile similarity (See **FigS4** and **Methods** for detail). The spatial distribution of assigned L1 labels was reconstituted and overlaid onto the ROI and largely agrees with the anatomical structures of the brain and expression pattern of the genes. A total of 330,156 subspots were quantified across 16 ROI. The relative proportion of cell classes are summarized in the donut chart. Scale bar = 1mm. (D) Circularized heatmap depicts the enrichment of genes and modules as a function of distance from the demyelinated (Sudan black negative, SB-) lesion core across 10x Visium spots pooled from 12 ROI with optimal contrast between SB and NFR staining (**FigS4** and **Methods**). The "Color Deconvolution" for Samples 1–4 was unsuccessful due to suboptimal contrast between SB and NFR staining, resulting in their exclusion from the lesion subregion assignment in the rim analysis; however, they are included for ME clustering analysis. Gene names starting with “*” indicate human (hs) or mouse (mm) orthologs of marmoset gene identification numbers (See **Table S9** for the full list). (E) Stacked column graph summarizes the relative proportion and distribution of classified ME as a function of distance from the SB-deprived lesion core across 10x Visium spots pooled from 12 ROI. (F) Stacked bar graphs summarize the relative proportion of L1 and L2 labels assigned to BayesSpace enhanced subspots across classified ME from 16 ROI (**left**). Stacked bar graph summarizes the relative proportion and distribution of L1 and L2 labels as a function of distance from the demyelinated lesion core across BayesSpace-enhanced subspots pooled from 12 ROI (**right**). The expression of gene sets used to infer L2 labels across subclusters are in **FigS8A.** hierarchical workflow was applied for L2 cell-type inference, which involved comparing the gene sets among subclusters within the same L1 cell class to assign an L2 cell type with the highest score. (G) Spatial distribution of assigned L2 labels is overlaid onto the ROI of a representative WM lesion. Spatial heatmaps of the ROI show the averaged expression of gene sets: L2_IMM.MoMφ for monocytes and macrophages (TMEM150C, CD36), L2_IMM.BP for B cells and plasmablasts (OSBPL10, JCHAIN), L2_IMM.T for T cells (KLRK1, NCR3), L2_IMM.DC for dendritic cells (CIITA, CPVL), L2_VE.homeo for vascular endothelial cell (SMAD6, VEGFC), L2_OLI.eae for oligodendrocyte subtype (VAT1L, SERPINB1, IGFBP3), L2_OPC.eae for OPC subtypes (EVA1A, A2M, GLIS3), L2_AST.eae for astrocyte subtypes (TPM2, TNC, SLC39A14). Scale bar = 1mm.

**Fig3.**
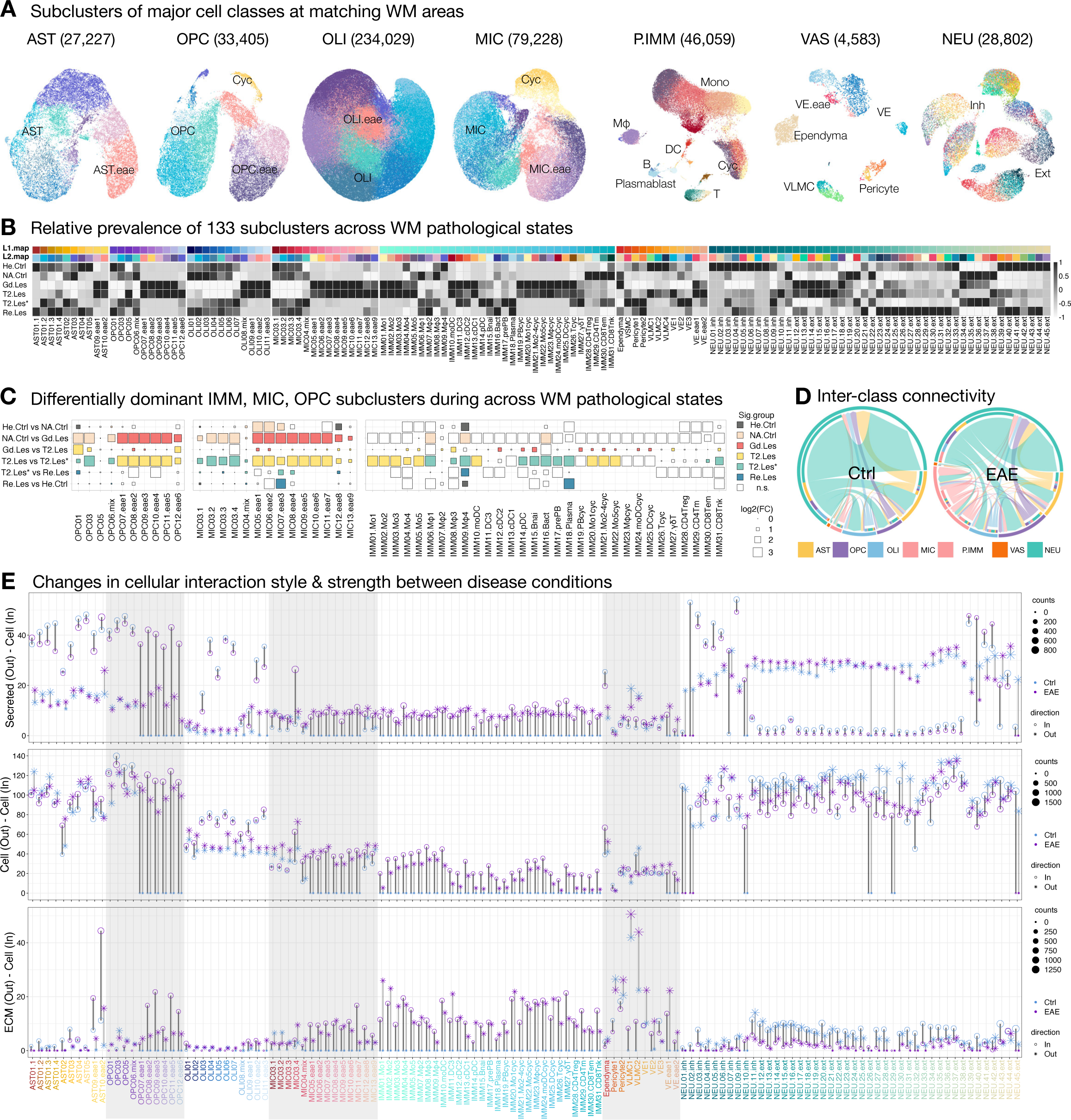
Temporally resolved cellular composition and connectivity mapping illustrate the succession of glial and immune cells with pathologically altered interactivity. (A) UMAP scatter plots display the L2 subclustering of white matter (WM) cell classes. The number of nuclei analyzed in each L2 UMAP plot is listed in parentheses. The relative distribution of homeostatic, cycling, and EAE-enriched glia are labeled. Abbreviations: Mono (monocytes), Mφ (macrophages), DC (dendritic cells), B (B cells), T (T cells), Cyc (cycling cells), VE (vascular endothelial cells), VLMC (vascular leptomeningeal cells), Inh (inhibitory neurons), Ext (excitatory neurons), T2.Les (<45 days old), T2.Les* (∼1000 days old), and Re.Les (prior T**_2_**-hyperintense signal that had resolved at the time of tissue collection) were grouped and analyzed. (B) Heatmap shows the z-scored number of nuclei for each subcluster across different WM pathological states. Two levels of color index are used for each subcluster to aid label tracking. L1.map coloring is consistent with labels of Cleveland dot plot in (**E**) and scatter plots in **FigS10**. L2.map coloring is consistent with UMAP plots in (**A**) and **FigS7, 8, 9, 11**. (C) Dot plots depict the change in nuclei proportion during the transition across WM pathological states. Squares show the relative enrichment of subclusters within each major cell class in each pair of pathological states. Significantly (false discovery rate, FDR < 0.05 & absolute fold change, abs(Log2FC > 0.25) enriched subclusters are colored accordingly. (D) Chord plots show the cumulative changes in interaction probability inferred by CellChat among major cell classes between control and EAE WM. The outer ring of the color bar represents the relative proportion of significant interactions employed by each cell class for each condition. The inner ring of the discontinuous color bars represents the relative proportion of signals sent to each cell class, and large arrows indicate signals received from each cell class. (E) Cleveland dot plots summarize the changes in outgoing (asterisk) and incoming (open circle) communications inferred by CellChat among subclusters of cells residing in WM of control (blue) and EAE (purple) animals. Three categories of interactions are quantified: secretes autocrine/paracrine signaling interactions (secreted–cell), cell-cell contact interactions (cell–cell), and extracellular matrix (ECM)-receptor interactions (ECM–cell). The level of signaling change for a matched subcluster pair between conditions is summarized as the bar length (light gray for outgoing and dark gray for incoming signals), and the alternating gray shaded columns distinguish major cell classes.

**Fig4.**
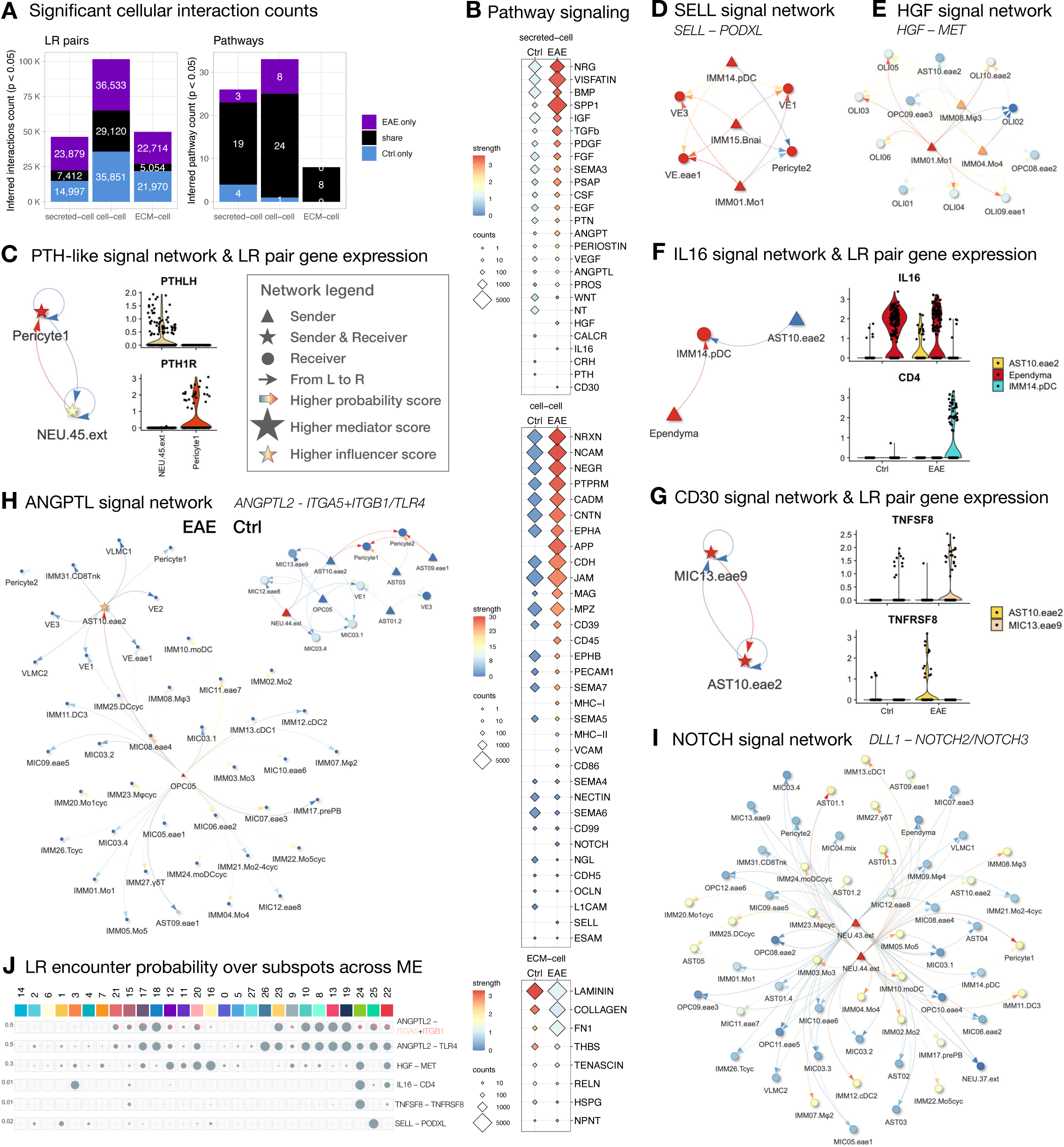
Comparative network analysis and spatial ligand-receptor mapping discover global changes in pathway signaling, uncover context-dependent interactions, and identify cellular links with microenvironmental significance. (A) Stacked bar graphs summarize the profile of ligand-receptor (LR) pairs and signaling pathways that are shared by or unique to WM of control and EAE animals. (B) Dot plots summarize the differences in pathway profile or strength among subclusters residing in the WM of control and EAE animals. (C) Inferred parathyroid hormone (PTH) signaling between pericytes and neurons, confirmed by the detection of PTHLH (Parathyroid Hormone Like Hormone) in neurons and PTH1R (Parathyroid Hormone 1 Receptor) in pericytes. The signaling role of each network is calculated by CellChat and summarized in visNetwork. Legend applies to panels D–I. (D) Inferred LR pairs (SELL–PODXL) of the SELL (Selectin L) pathway between vasculature and immune cells in EAE animals. (E) Inferred LR pairs (HGF–MET) of the HGF (Hepatocyte Growth Factor) pathway between immune cells, oligodendrocytes, OPC, and astrocyte subtypes in EAE animals. (F) Inferred LR pairs (IL16–CD4) of IL16 (Interleukin 16) pathway between ependyma, plasmacytoid dendritic cells (pDC), and EAE enriched astrocyte subtype in EAE animals. (G) Inferred LR pairs (TNFSF8–TNFRSF8) of the CD30 (Tumor Necrosis Factor Receptor Superfamily Member 8) pathway between EAE enriched microglia and astrocyte subtypes in EAE animals. (H) Inferred LR pairs (ANGPTL2–ITGA5+IGAB1, ANGPTL2–TLR4) of the ANGPTL (Angiopoietin-like) pathway between multiple subclusters in WM of control and EAE animals. (I) Inferred LR pairs (DLL1–NOTCH2, DLL1–NOTCH3) of the NOTCH pathway between multiple subclusters in WM of EAE animals. (J) Pie charts summarize the proportion of subspots with detection of both ligands and receptors for each inferred pathway in panels C–I across classified ME. The ratio of subspots with targeted ligand overlapping completely with receptor and cofactors (if applicable) are colored in gray; if only one of the receptor components is involved, it colored accordingly.

**Fig5.**
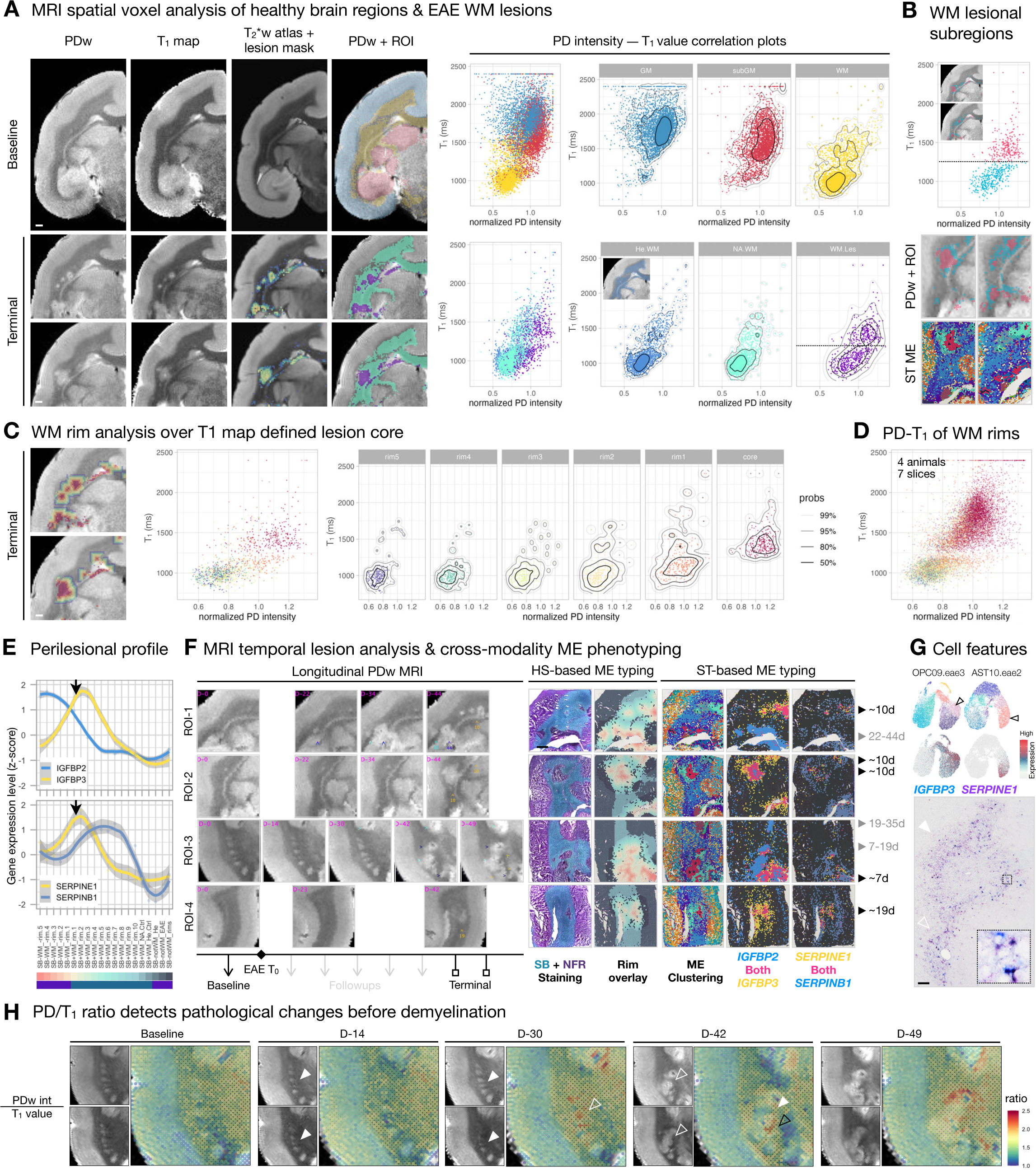
MRI features distinguish lesion subregions, mark the trajectory of white matter (WM) pathology, and phenotype lesion microenvironments (ME) with temporal significance. (A) **Left**: To identify MRI features that could inform lesion dynamics, proton density-weighted (PDw) images and T**_1_** maps acquired in the same imaging session were registered to a T**_2_**\*w MRI atlas at baseline and terminal time points. The lesion masks were created by subtracting the normalized baseline intensity value from the terminal PDw image and then overlaid onto the registered T**_2_**\*w MRI atlas (**FigS4** and **Methods**). The regions of interest (ROI) consist of atlas-annotated anatomical structures and lesion subregions, which were used to group and color-code each voxel and then overlaid onto PDw images for visualization. Scale bar = 1mm. **Right**: Scatter plots with density contours (legend in (C)) show the correlation of PDw intensity and T**_1_** value (PD-T**_1_**) for each voxel across ROI, as indicated on overlaid PDw images. As expected, cortical gray matter (GM) and subcortical gray matter (subGM) have higher T**_1_** value (longer longitudinal relaxation time) than WM. The normal appearing WM (NA.WM) and WM lesion (WM.Les) areas from the terminal PDw image (**bottom**) are compared against the equivalent areas (He.WM) at baseline (demonstrated in the inset). The PD-T**_1_** distribution of NA.WM largely agrees with He.WM, but there is a horizontal shift in PDw intensity and a vertical split in T**_1_** values (T**_1_** = 1250, annotated by a horizontal dashed line) into two populations for WM.Les. (B) **Top**: A cutoff T**_1_** value of 1250 ms (horizontal dashed line) was applied to WM.Les voxels, which are color-coded accordingly on the scatter plot and the overlaid PDw image (inset). **Bottom**: The subregional structure of PD-T**_1_** values resembles that identified by spatial transcriptome ME clustering. Voxels with high T**_1_** values typically reside at the lesion core, whereas voxels with low T**_1_** values primarily populate the lesion edge. (C) 5 concentric rims, outward from the PD-T**_1_** defined lesion core, color WM subregions on the PDw image and scatter plots. The PD-T**_1_** distribution of the rim5 area (750 μm away from the lesion core) is similar to that of He.WM, while PDw values gradually increase as voxels approach the lesion core. Scale bar = 1mm. (D) Scatter plot summarizes the PD-T**_1_** distribution of WM rims across 7 EAE brain slices from 4 animals, uncovering a similar WM pathological trajectory to that shown in (**C**). (E) Line plots summarize the relative abundance of *IGFBP2*, *IGFBP3*, *SERPINE1*, and *SERPINB1* expression as a function of distance from the demyelinated (Sudan black (SB) negative) lesion core. Black arrows pointed to the intersection of SB^+^ and SB^-^ areas. (F) Relative expression profile of IGFBP and SERPIN families differentiate lesions by age. **Left**: Snapshots of PDw images across time in 4 representative ROI from 3 animals. Days (D) post EAE induction (T0) are labeled in magenta for each ROI, and lesion age is estimated retrospectively from the serial MRI. The appearance of each lesion is annotated by an arrowhead, and different arrowhead colors are used to track different lesions. **Right**: The MRI-matching ROI were further imaged through the scope of histological staining (HS) and spatial transcriptome (ST) to subdivide brain regions into ME. The relative abundance is binarized by filtering the gene expression of the IGFBP (z-score >1) and SERPIN (z-score > 0.5) family, such that spots below the cutoff are colored dark gray. Scale bar = 1 mm. (G) **Top**: UMAP plots of OPC and AST colored by L2 subcluster and gene expression. Lesion edge-enriched genes, such as *IGFBP3* and *SERPINE1*, are highly expressed by subtypes of OPC and AST, respectively. **Bottom**: Immunohistochemical staining of IGFBP3 (blue) and SERPINE1 (purple) in a midcoronal section of the marmoset brain with enlarged area in 50 x 50 μm^2^ box. High IGFBP3 and SERPINE1 labeling are in close proximity to a dilated blood vessel (open arrowhead) and are distant from a flattened blood vessel (solid arrowhead). Scale bar = 100 μm. (H) PDw MRI and T**_1_** map images from baseline (before EAE induction) and 4 follow-up time points after EAE induction. Normalized PDw intensities and T**_1_** values were extracted, and the PD/T**_1_** ratio was calculated and overlaid onto the T**_1_** map as heatmaps. Open arrowheads indicate MRI-identifiable tissue changes, solid arrowheads indicate normal-appearing brain area. White arrowheads point to a similar brain area across time and imaging contrasts, and the black arrowhead point to a different brain area with high PD/T**_1_** ratio.

### ▪ Approach: cross-modality analysis resolves the spatial distribution of major cell populations and identifies 5 microenvironment groups that mark the development and resolution of WM pathology

Given the differential sensitivity of each analysis modality (**FigS3**), we first utilized histopathology (SB lipid staining) to define the lesion as a SB^-^-WM area. We then annotated intralesional, perilesional, and extralesional (IL, PL, EL) WM subregions as a function of distance from lesion core to analyze spatial transcriptomes (**FigS4B**). By integrating 10x Visium spots across 16 ROI with Seurat (**Methods**), we clustered a total of 28 microenvironments (ME) into 5 groups by transcriptomic profile similarity and identified key genes differentially expressed in each group (**Fig2A**). ME group i (ME14, 2, 6, 1, 3, 4, 7, 21, 15, 17, expressing *NEFH*, *RASGRF1*, *HIVEP2*, *TSPAN13*, *GDAP1*) and group ii (ME26, 27, 5, 0, 16, 20, 11, expressing *MOG*, *FGFR2*, *LPAR1*, *LGI3*, *BCAN*) are enriched in SB^-^-GM and SB^+^-WM, respectively, which agrees with known anatomy of the brain (**FigS5A-B**). We found ME group iii (ME23, 9, 10, 8, 13, 19, expressing *FBP1*, *S100A12*, *MMP9*, *ITGA5*, *IFI30*) to be enriched in MRI-defined T_2_ lesions (**FigS5B**) and ME group iv (ME22, 25, 24, expressing *MYH11*, *PRLR*, *FHAD1*, *CCDC180*, *CFAP52*) to delineate brain borders (BB), such as meninges, blood vessels, and ventricles (**FigS5A**). Interestingly, we found ME group v (ME12, 18, expressing *BCAS1*, *ADAMTS4*, *SERINC5*, *PTPRZ1*, *CERS2*) to be enriched at the border of WM lesions (**FigS5D**). While ME group v partially shared gene expression with groups ii and iii (**Fig2A**), cells in ME group v are particularly enriched with genes that are important for OPC differentiation, early myelinating oligodendrocytes, and remyelination (43–47), suggesting the presence of reparative activities at the lesion edge.

The neighboring spots of 10x Visium are 100 μm apart, often wider than the distance between cell pairs in the marmoset brain; therefore, we employed BayesSpace tool to enhance spatial resolution into subspots (∼20 μm apart) in order to better identify the source of regionally restricted signals. Leveraging L1 and L2 markers identified by snRNA-seq analysis (**FigS8-9**), we inferred cell type for each BayesSpace enhanced subspot by the relative enrichment of the denoted gene sets using a hierarchical workflow (**FigS4C, Methods**). We then cross-indexed the level of regional differentially expressed genes (rDEG, **Fig2D**), the prevalence of ME groups (**Fig2E, FigS5**), the gene modules that varied along UMAP trajectory (**FigS6B**), the expression of ME enriched genes (**FigS7A**), and the inferred L1 cell classes (**Fig2B-C**) and L2 subclusters (**Fig2F-G**) to WM subregions in relation to the spatial organization of the lesion core. With this cross-modality analysis, we aimed to dissect signaling dynamics by their environmental and cellular compositional significance.

As expected, we found L1.IMM population (labeled by *ARHGAP15*, *PTPRC*, *HCRTR2*, *GPNMB*, *ITGA4*, *CD36*, *SKAP1*) dominates the lesion core (**Fig2C**). Within L1.IMM population, >50% of the subspots were mapped to EAE-enriched microglia (L2_MIC.eae, expressing *MSR1*, *MLANA*, *FLT1*, *C3*), followed by monocytes and macrophages (L2_IMM.MoMφ, expressing *TMEM150C*, *CD36*), B cells and plasmablasts (L2_IMM.BP, expressing *OSBPL10*, *JCHAIN*), and dendritic cells (L2_IMM.DC, expressing *CIITA*, *CPVL*) at the lesion core (**Fig2F-G**). To a lesser extent (∼10%), L1.OPC population (labeled by *PDGFRA*, *GALNT3*, *TNR*, *EVA1A*) constitutes EAE-enriched OPC (L2_OPC.eae, expressing *EVA1A*, *A2M*, *GLIS3*) and cycling OPC (L2_OPC.cyc, expressing *CENPP*, *TOP2A*) was mapped to the lesion core (**Fig2F**). Together, these findings suggest that IMM and OPC cell classes are the prevailing players at the lesion core.

The cell type inference workflow employed here chooses not to display the probability of all possible cell types as a relative percentage, in which only one cell type with the highest score of the denoted gene set was assigned to a subspot that contains mixed signals from more than one cell type (**FigS4C**). Although it enables visual representation of the principal cell types associated with different anatomical and pathological structures (**Fig2C**, **FigS8**), it does not fully represent the complexity of cellular composition, especially when cell density is high (<20 μm apart), such as the hypercellular acute/subacute lesion core. In particular, this workflow might lead to under-emphasis of populations expressing unique but lower-level markers, such as the vascular cell class (VAS), but the relative enrichment and spatial organization of such cells can still be retrieved by gene set expression plots.

For example, the distribution of vascular cells (L1.VAS, expressing *ITGA1*, *ARHGEF28*, *DNAH11*, *PDGFRB*, *DCN*, *MECOM*, **Fig2C**) and vascular endothelial cells (L2_VE.homeo, expressing *SMAD6*, *VEGFC*, **Fig2G**) were resolved, agreeing with the vessel features identified by MRI and histological staining (**FigS2-3**), and with prior knowledge that MS-like WM lesions expand around a central vein (9,10,28). Such resolution makes it possible to localize certain cell types, in particular T cells, dendritic cells, and a subset of B cells and plasmablasts, to the perivascular area. Indeed, ME22 (enriched with VAS markers, *SLC6A13*, *MYH11*, *DCN*, *IGF2*, *SLC13A4*, **FigS5C**) was detected at the lesion core (**Fig2E**, SB-WM_-rim.5), regardless of its overall low prevalence (610 out of 55,026 spots, **Fig1D**). Additionally, gene modules involved in the regulation of blood vessel morphogenesis (Knn.m2), blood vessel endothelial cell migration (Knn.m9), and angiogenesis (PG.m8) are highly enriched in the lesion core (**Fig2D**, **FigS6B, TableS4**). Moreover, rDEG of the lesion core (**Fig2D**), *IFI30* (Interferon-gamma-inducible protein 30) and *DPP4* (dipeptidyl peptidase-4) are involved in sprouting angiogenesis (48) and maintaining the level of pro-angiogenic factors (49), and we found these to be expressed by vascular leptomeningeal cells (VLMC) and immune cells (**FigS7B**). Together, we found that the lesion core harbors unique ME involving VAS cells and signaling important for vessel health. Our model, workflow, and data quality are thus sufficient to identify factors known to be important for MS, such as angiogenesis (50), stressing the pertinency of our work to nominate new candidates for MS research.

### ▪ Intralesional WM: an epicenter of innate and adaptive immune activities, comprised of microenvironments involved in angiogenesis, lipid metabolism, cell proliferation, and ferroptosis

In addition to involvement in angiogenesis, *IFI30* marks the high infiltration of immune cells (51) and is highly enriched in ME group iii (**Fig2A**). Genes that are essential for the weakening of VE junctions (*TM4SF19*, (52)) and lymphocyte trans-endothelial migration (*CD52*, (53)) are highly enriched in the intralesional WM (**Fig2D**). Moreover, intralesional WM-enriched ME group iii (ME19, 13, 8) and ME18 (**Fig2E**) are marked by genes that are primarily expressed by immune cells (*MMP9*, *FBP1*, *S100A12*, *GPNMB*, *CXCR4*, **FigS7A-B**). Interestingly, *CXCR4* (a hub gene in MS-related pathways (54)) is pathogenically regulated by Epstein-Barr virus (EBV) infection of B cells (55), which is thought to be in the causal chain of MS (56). Genes involved in lipid storage/catabolism and macrophage differentiation/activation (*CD36*, *SLC37A2*, *MSR1*, *NR1H3*, *PLA2G7*) are differentially enriched in intralesional WM and are primarily expressed by myeloid cells (microglia, monocytes, and macrophages) and γδT cells (**FigS7B**). Genes that are shared across ME19, 13, 8 (*PTTG1*, *PCNA*, **FigS7A**) are expressed by all cycling immune cells (IMM19-26, **FigS7B**), with concomitant enrichment of gene modules involved in DNA repair and apoptosis (PG.m14, PG.m8, **TableS4**). Together, this suggests active myelin destruction and a pro-inflammatory state in the intralesional WM, consistent with known pathology of active MS lesions.

Compared to perilesional WM, ME19 and 13 are significantly elevated in intralesional WM (**FigS5C**). ME13-enriched genes (*SLC15A1*, *TSHR*, *MLANA*, *CYP27A1*, *FIGLA*) are mainly expressed by myeloid cells, which are essential players in chemokine/cytokine production and responses (PG.m26, PG.m8, Knn.m2, **TableS4**). Interestingly, *FIGLA* (a sex-specific transcription factor that suppresses sperm-associated genes (57)) is not detected in microglia of control but is particularly elevated in myeloid cells of EAE (**FigS7B**); whether it associates with a sex bias in MS prevalence is unknown. Genes that distinguish ME19 (*MZB1*, *POU2AF1*, *GBP5*, *LTB*, *CD2*) from other ME are mainly expressed by B and T cell lineages (**FigS7B**), agreeing with the regional enrichment of gene modules involved in B and T cell activation and antigen presentation (Knn.m2, Knn.m13, **Fig2D, FigS6B, TableS4**). In parallel, rDEG encode various immunoglobulins (*JCHAIN*, *IGLC*, *IGLA*, *IGKC*, *IGHGs*, *FCGRs*) and major histocompatibility complex (MHC) class I (*HLA-B*, *HLA-G*) and II (*HLA-DPB1*, *CD74*) are elevated in intralesional WM (**Fig2D**). Genes (*SDF2L1*, *EDEM1*) that are involved in misfolded protein binding (Knn.m30, **TableS4**) are highly expressed by the B cell and plasmablast (BP) lineage, except naïve B cells (IMM15.Bnai, **FigS7B**). BP population (L2_IMM.BP) is particularly enriched in ME19 compared to other ME (**Fig2F**) and is in proximity with blood vessels and lesion border (**Fig2G**), suggesting the source and location of humoral immune response to myelin destruction. All told, a full spectrum of adaptive and innate immunity is manifest during the development of WM lesions in marmoset EAE.

Interestingly, in addition to the immune cell involvement that is often the hallmark for intralesional WM-enriched ME (ME19, 13, 8, 18), *IQCK* (IQ motif containing K), a novel risk factor for Alzheimer’s disease (AD) (58–61), is uniquely enriched in astrocytes (AST) and ependyma of the area (**FigS7B**). More broadly, we found that ME with heavy glial/vascular contributions increased considerably toward the lesion edge when ME19 and 13 decreased drastically from their peak at the lesion core (**Fig2E**). For example, genes that suppress ferroptosis (*AIFM2*, *MGST1*, (62,63)) are enriched in ME19, 13, 8, 18 (**FigS7**), whereas genes that induce ferroptosis (*SLC7A11*, *TMEM164*, (64,65)) are expressed by AST, VLMC, and ependyma of ME group iv, which is significantly enriched in perilesional WM compared to intralesional WM (**FigS5C**). Together, this suggests a transition from immune (intralesional WM) to glial/vascular (perilesional WM)-dominant ME with a mixture of destructive and protective signals as lesions evolve, and we further deconvolute this complexity in the following sections.

### ▪ Perilesional WM: a junction of immune-vascular-glial cell interactions, comprised of microenvironments involved in lesion expansion, settlement, and remyelination

Compared to intralesional and extralesional WM, ME group ii (ME20), iii (ME10, 8), iv (ME24), and v (ME12, 18) are significantly enriched in perilesional WM (**FigS5C**), which underscores the level of signal diversity at the lesion border. Genes that distinguish ME8, 10 (*SERPINE1*, *HEYL*, *HBEGF*, *EVA1A*, *CRABP2*) from other ME are highly expressed by EAE-enriched OPC and AST subclusters (**FigS7B**), which primarily populate the inner (L2_OPC.eae) and outer (L2_AST.eae, expressing *TPM2*, *TNC*, *SLC39A14*) rings of perilesional WM in relation to the lesion core (**Fig2C, F, G**). Particularly, *EVA1A* (an autophagy regulator that typically benefits human health (66,67)) uniquely distinguishes all OPC.eae subclusters (OPC07-12) from homeostatic populations. Genes enriched in ME20, 12 (*LPAR1*, *ANLN*, *TMEM144*, *FAM222A*, *SYNJ2*) are primarily expressed by all oligodendrocytes (OLI), and genes enriched in ME12, 24 (*MSMO1*, *MVD*, *MYOC*, *CYP2J2*, *SLC2A1*, *BGN*) are expressed by AST, OLI.eae, VAS. In addition to L2_OPC.eae and L2_AST.eae, the proportion of L2_OLI.eae (*VAT1L*, *SERPINB1*, *IGFBP3*) and L2_VE.eae (*PDLIM1*, *ADAMTS1*, *TNFRSF6B*) increases across perilesional WM-enriched ME (**Fig2F**). Compared to their homeostatic counterparts (**FigS7B**), all EAE-enriched glial and vascular subclusters express more *CRYAB* (heat shock protein) that elevates at the SB^+^ perilesional WM and extends into extralesional WM (**Fig2D**). Together, these findings suggest that glial and vascular cells respond to stress signals absence of apparent in situ myelin loss, which might be beneficial in maintaining the physiological functions of cells.

At the intersection of SB^+^- and SB^-^-WM, overlapping signals involved in hemostasis, inflammation, proliferation, and tissue remodeling phases are at play. We found an elevation of the *MAFF* transcription factor at the perilesional WM (**Fig2D**), which indicates an increased blood vessel permeability through inhibiting inter-endothelial proteins (e.g. ZO-1, occludin, claudin-5 (68)). Genes involved in complement and coagulation cascades (KEGG:04610, **TableS4**) are elevated across PL WM-enriched ME, which prompted us to perform supervised analyses focusing on these systems to understand intercellular communication between glial and vascular cells. The central elements (*C3*, *C5*) of the complement system are primarily expressed by MIC, and the classical pathway components (*C1QA*, *C1QB*, *C1R*, *C1S*) are expressed by IMM and VLMC (**FigS7B**). In contrast, the lectin pathway components (*COLEC11*, *FCN3*, *MASP1*) are enriched in VE, pericytes, and AST, and *C7* in the terminal pathway is expressed by VLMC (**FigS7B**). Interestingly, we found that *CFB* in the alternative pathway is expressed by VE, ependyma, and AST.eae, and the levels of pathway inhibitors (*CFI*, *CFH*) are elevated in VAS, AST, and OPC.eae (**FigS7B**). We found *CFB*^+^ ependyma to be unique to the EAE condition (**FigS7C**), suggesting an active complement response to EAE at the CSF-brain barrier. Moreover, we found factors that promote (*PLAU*, *ANXA2*, *S100A6*) and inhibit (*SERPINE1*) fibrinolysis to resolve coagulation by controlling plasmin production (69) are uniquely expressed by AST.eae (**FigS7B**). Factors that mediate leukocyte trafficking (*CYR61* (70)), anti-inflammatory activities (IL1R2, a non-signaling “decoy” receptor (71)), and wound healing (*IGFBP3* (72)) are significantly enriched at the PL WM. Consistent with rDEG results, the gene module involved in cell junction assembly, fibronectin binding, keratinocyte proliferation, angiogenesis, and response to insulin (PG.m7, **TableS4**) are highly enriched at the lesion border. Together, these results suggest that ependyma, along with AST and VAS, contribute to complement-mediated tissue damage by initiating and regulating antibody-independent (lectin and alternative pathways) cascades. Additionally, AST.eae appears to hold a central role in coordinating multiple signals pertinent to different phases of wound repair.

In parallel, gene modules involved in oligodendrocyte differentiation and regulation of myelination (PG.m7, **TableS4**) are enriched at PL WM, and DEGs that distinguish ME18, 12, 11, 20, 16 (*CERS2*, *SERINC5*, *ADAMTS4*, *CAGE1*, *REEP3*, *BCAS1*) from other ME are primarily expressed by oligodendrocyte lineage (**FigS7B**), which prompted us to look for ME that are particularly relevant to remyelination. We computed and selected DEG that are enriched in differentiating OPC (OPC05, enriched with *TNFRSF21*, *BCAS1*, *SERINC5*, *RHOQ*, *ENPP6*) compared to other OPC/OLI populations and calculated a differentiating OPC gene module (dOPC.m) score across candidate ME. We found that ME18 has the highest dOPC.m score (z-score = 2.54, **TableS5**), primarily populating the SB^-^-WM area, and that the density of spots increased with the age of lesions (**FigS7D**), suggesting the presence of remyelination activities at the PL WM that starts as early as 10 days post-EAE induction. Overall, we see a great ME diversity at the PL WM, with overall a transition from inflammation-related ME to ME featuring a complex glial-vascular interaction.

### ▪ Extralesional WM: an area responding to diffuse activation, comprised of sensitized microenvironments prone to develop new lesions

Compared to PL and NA WM, ME group ii (ME26, 5, 0, 16, 11), iii (ME23, 9), and iv (ME25) are enriched in EL WM. Particularly, the proportion of ME23, 9 appears to be elevated in SB^+^-WM of EAE compared to control and SB^-^-WM of EAE (**FigS5B**). No unique genes clearly distinguish ME23, 9 from the rest of ME; instead, we found a graded expression profile shared by ME groups ii-iv to different degrees, suggesting a transition between homeostatic and pathologic states. Genes that are elevated across many EL WM-enriched ME (*GFAP*, *APLP1*, *CALCA*) are involved in reactive glial responses, plaque neurotoxicity of AD, and vessel dilatation (37,73–76). Gene modules involved in endothelial cell differentiation and blood circulation are enriched in ME25 (Knn.m21, PG.m19, **TableS4**). Whereas genes that are expressed by OLI and important for WM health are enriched in ME26, 5, 0, 16, 11 (*PLP1*, *TF*, *MBP*, **FigS7A-B**), as expected, gene modules regulating cytokine response, autophagy, and double-strand break repair (Knn.m5, PG.m12, **TableS4**) are enriched in ME16 in EL WM (**Fig2D**, **FigS7A**). Interestingly, we found that *BGN* (a critical ECM regulator that boosts inflammatory signaling through Toll-like receptors (77)), expressed by pericytes and VSMC, is significantly enriched in the EL WM (**Fig2D**, **FigS7A**). Together, these results suggest the presence of global glial and vascular responses to EAE induction in EL WM, where dilated vessels and stressed glia could indicate the impending development of new focal lesions.

### ▪ Normal-appearing tissue: a domain containing latent components of EAE with altered metabolic processes

To further understand the extent of EAE-related changes, we compared WM and GM without a clear histology detectable change to their healthy counterparts. While most genes are shared between EL, NA, and healthy WM (**Fig2D**), *PTGDS* (an anti-inflammatory enhancer that suppresses Aβ accumulation (78,79)) is particularly enriched in the WM of EAE animals. The gene module involved in chemotaxis and cellular lipid metabolic process is elevated in NA WM compared to healthy WM (Knn.m14, **Fig2D**). As expected, non-WM areas (cortical and subcortical GM), are enriched with genes and modules important for the function of neurons in both control and EAE. However, genes that increase pyruvate and lactate in serum and CSF are elevated in EAE GM and IL WM compared to control (HP:0002490, HP:0002151, HP:0003542, Knn.m6, PG.m6, **Fig2D**, **TableS4**). Compared to healthy GM, gene modules that regulate synapse assembly, synaptic vesicle exocytosis and priming are reduced in EAE GM (PG.m18, Knn.m19, **Fig2D**, **TableS4**). Moreover, genes encoding proteins that are elevated at the BBB of AD brain (*PRL15*, (80)), function to desensitize ferroptosis (*TRIB2*, (81)), and interact with vimentin to influence cholesterol transport (*OSBP2*, (82)) are decreased in the EAE GM compared to control (**Fig2D**). Together, these results suggest the presence of pathological changes at sites not detectable by conventional histology, which reiterates the importance of considering the additive effects of global parenchymal alterations to understanding the pathogenesis of inflammatory demyelination.

### ▪ Transition to diseased microenvironment: OPC and microglia are among the first responders in EAE, followed by enrichment of monocyte derivatives, and replaced by lingering lymphocytes as lesions evolve

To further understand the dynamics of intercellular interactions as tissue transitions from physiological to pathological states, we characterized cellular composition and mapped cellular connectivity as lesions evolve using snRNA-seq. A total of 595,472 nuclei were recovered in L2 analysis, and 133 subclusters were annotated across conditions, with 36 subclusters unique to the EAE condition (**FigS9-10**). For glial cells, we numbered subclusters by their L1 cell class identity followed by a crude division of their prevalence in EAE samples; subclusters enriched in EAE samples are denoted with “eae.” Compared to control, the proportion of MIC and OPC in EAE expanded about 5 and 2 times, respectively (**FigS9B**). Unlike MIC and OPC, no cycling AST cluster was observed in EAE (**FigS9C-D**). For immune cells primarily derived from the periphery (P.IMM), we used a convention of numerical order (IMM01-31) followed by a crude division of leukocyte lineage, for they were found almost exclusively in EAE samples. Among P.IMM, most are monocytes (56.4%), followed by cycling leukocytes (15.8%), macrophages (9%), dendritic cells (8%), T cells (6.6%), and B cells and plasmablasts (4.2%) (**TableS6**). For neurons, we used a numerical order followed by a crude category of neurotransmitter. We labeled NEU subclusters enriched with *GAD1*/*GAD2* expression as inhibitory (inh, 18.2%) and others as excitatory (ext, 81.8%) (**TableS6**).

We explored the tissue-specific and shared responses to EAE by comparing parietal WM (pWM), parietal cortex (pCTX), and lateral geniculate nucleus (LGN) to their healthy or NA controls, with a total of 189,091 nuclei analyzed (**TableS7**). We found a significant expansion of MIC and P.IMM partitions in all tissue of EAE animals (**FigS11A**); however, the compositions of MIC, OPC, and AST partitions were unique to each tissue type (**FigS11B**). Specifically, the OPC and AST compositions of EAE were more similar in the pCTX and LGN compared to that of pWM. On the other hand, the MIC composition of EAE was more similar in the pWM and pCTX regions compared to that in the LGN region. Interestingly, we observed considerable similarity in the enrichment of transcription factors in EAE across different tissue types for each glial cell class (**FigS11C**). The shared transcription factors across tissue types are involved in myeloid/foam cell differentiation and ISGF3 complex (Type-I interferon signaling) in MIC, repression of transcription activity in OLI, promoter binding in OPC, and mineralocorticoid receptor (hormone response) binding to transcribe coregulators in AST (**FigS11D**). These findings provide an initial framework to understand the divergence and convergence in cellular and transcription factor changes across tissue and cell types in response to EAE.

Given that we found no unique subcluster in response to EAE across different coarse tissue types, we focused on the better-sampled WM areas to map their cellular dynamics as lesions develop, analyzing a total of 453,333 nuclei from matched brain areas (**Fig3A**, **TableS2**). WM samples are grouped by inoculation status and radiological findings, which combined can inform the temporal trajectory of tissue damage under pathological insults. WM from healthy control animals (He.Ctrl), and from EAE animals without radiological signs of demyelination (NA.Ctrl), with Gd-enhancing lesion indicating an open BBB (Gd.Les), with T_2_-hyperintense signal for <45 days (T2.Les) or >1000 days (T2.Les*), and with prior T**_2_**-hyperintense signal that had resolved at the time of tissue collection (Re.Les), were grouped and analyzed.

All 133 subclusters are collectively present in the WM samples; however, we found that acute lesion stages (Gd and T2 lesions) tend to have different cellular profiles than other WM groups (**Fig3B**). We quantify this observation by a proportional test for subclusters within IMM, MIC, and OPC classes across stages (**Fig3C**). Compared to healthy control and Gd lesion, NA control is enriched with OPC06.mix and MIC04.mix subclusters, suggesting that these cells are early responders to demyelination-independent stimuli and are transitioning from homeostatic to pathologic states. Compared to NA control and T2 lesion, Gd lesion is enriched with naïve B cell (IMM15.Bnai, *SELL*^+^), plasmacytoid DC (IMM14.pDC, *SELL*^+^), conventional DC (IMM12.cDC2, *CCR7*^+^), and cycling glial and immunes cells (OPC08.eae2, MIC05-06, IMM19-25, *CENPP*^+^). Given that *SELL* (L-selectin) promotes the initial tethering and rolling of leukocytes to the endothelium (83), IMM14-15 likely represents a population that has not yet entered brain parenchyma for further specialized subtype differentiation. The expression of *CCR7* (a chemokine receptor required for DC maturation and lymphocyte migration (84)) and highly proliferative glial and immune cells indicate an active inflammatory propagation stage when the BBB is open.

As lesions develop, monocytes (IMM01-05), monocyte-derived DC (IMM10.moDC), and proliferating monocytes (IMM20-22) continue to be the dominant leukocytes within T2 lesions. *CD44*^+^ OPC (OPC10-12 (85)), *TSHR*^+^ microglia (MIC08-13 (86)), and *ITGAX*^+^ gamma-delta T cells (IMM27.γδT (87)) become more prevalent. Compared to younger T2 lesions, the composition of glial cells in older T2 lesions (L2.Les*) returns to a homeostatic-like profile (OPC01-06, MIC03-04 dominant) and is similar to that of resolved lesions and healthy control WM (**Fig3C**). However, we found that macrophages (IMM06-09), pDC, B lineage cells (IMM14-18), CD8 effector memory T cells (IMM30.CD8Tem), and *KLRK1*^+^/*KLRD1*^+^ natural killer T cells (IMM31.CD8Tnk) lingered in older T2 lesions. As lesions resolve, there is an enrichment of plasma cells (IMM18.Plasma) and *LYVE1*^+^ perivascular macrophages (IMM09.Mφ4), though the proportion of Mφ4 never recovers to that of control (**Fig3C**). Similarly, we found a persistent enrichment of a microglial subcluster (MIC07.eae3, expressing higher *IGFBP3* and *TUBB2B* than the homeostatic subclusters) in EAE WM, including older T2 lesions and resolved lesions, indicating the presence of a long-lasting microglial state associated with EAE.

### ▪ Intercellular connectivity in diseased microenvironments: a global shift in the connectivity landscape across cell types, particularly for ECM-mediated signaling

To further understand the significance of the highly diverse cellular composition in EAE, we compared the intercellular connectivity across conditions by querying the ligand-receptor (LR) relationships among subclusters with balanced nuclei numbers (**Methods**). Given that resident and peripheral immune cells are the most expanded cell types in response to EAE, we found increased interactivity between immune and all other cell classes in EAE compared to controls, as expected (**Fig3D**). Also, predicted interactions between OPC and other cell classes were greatly increased, whereas interactions between AST and NEU were decreased. We quantified this observation across conditions and summarized intercellular connectivity by signaling direction and type. Cells expressing ligands in established LR pairs are denoted as the senders of outgoing signals (Out), and cells expressing receptors as receivers of incoming signals (In). We grouped types of ligands by their mode of action, separately quantifying secreted autocrine and paracrine signaling (secreted–cell), cell contact-mediated signaling (cell–cell), and ECM-mediated signaling (ECM–cell) (**Fig3E**, **FigS12**).

In healthy WM, we found AST, OPC, OLI, and NEU.inh to be the primary receivers of the secretory signals and VLMC and pericytes to be the major senders of the secretory and ECM signals (**FigS12**). In EAE WM, we found an overall decrease in the receipt of secretory signals for homeostatic-enriched glial subclusters (AST01-05, OPC01-06, OLI03-07), and an increase in ECM interactions for EAE-enriched glial subclusters (AST09-10, OPC07-12, OLI08-11). Moreover, immune cells (IMM01-31) generally strongly interact with ECM, whereas NEU reduce their cell-cell and ECM-cell contact strength (**Fig3E**).

Interestingly, we observed drastic changes in the communication profile across conditions for AST10.eae2, AST02, MIC03.4, ependyma, pericytes, VLMC, and VE, prompting us to analyze their regulatory roles in signaling networks (**Methods**). While most signaling pathways are shared between the control and EAE, the profile of significant LR pairs is vastly different, indicating a global change in communication partners across conditions, especially for ECM-cell interactions (**Fig4A**). Interestingly, across shared signaling pathways, we found increased strength of the secretory signals and cell-cell contacts but diminished strength of ECM-cell signaling in EAE compared to control, suggesting increased short- and long-range cellular communication and reduced structural integrity in EAE WM (**Fig4B**). In healthy WM, we found parathyroid hormone-like hormone (PTHLH) signaling between pericytes and a subcluster of excitatory neurons (NEU.45.ext, *SLC17A6^+^,* **Fig4C**), suggesting subcortical neurovascular crosstalk to regulate calcium levels and blood flow in homeostasis (88,89). Given that the tissue contribution of NEU.45.ext is biased toward the parietal corpus callosum (pCC) sampling site (**SourceData_FigS9**) with only 1 matched T2.les sample (**FigS1D**), whether this neurovascular crosstalk is attenuated in EAE requires further study. Interestingly, our data suggest that *SELL*^+^ monocytes (IMM01.Mo1) may contact vascular cells for regulating entrance to the parenchyma (**Fig4D, 4J**), and that they communicate with oligodendrocyte lineage to inhibit differentiation and stimulate OPC proliferation via a secretory HGF signal (**Fig4E** (90,91)). Together, these results suggest that crosstalk between immune and glial cells might impact myelin plasticity in pathologic conditions.

In EAE WM, we found intimate interactions between vascular/glial-immune (MHC-I, MHC-II), glial-immune (CD45, CD86), immune-vascular (VCAM), and all-all cells (APP), except the involvement of AST cell class (**FigS13A-F**). Nevertheless, AST, along with other cell classes, considerably altered their communication partners to signal cell growth (EGF, PDGF, VEGF), adhesion and migration (APP1, SEMA7A, Tenascin), and neural development (SEMA7A, NGL, SEMA5A) in EAE compared to control environments (**FigS13G-N**). Additionally, AST10.eae2 uniquely interacts with IMM14.pDC and MIC13.eae9 through secretory IL16 and CD30 signals, respectively, suggesting a role in initiating and regulating immune responses (**Fig4F-G**). Moreover, AST10.eae2 appears to propagate ANGPTL (angiopoietin-like) proinflammatory signal received from differentiating OPC (OPC05) to VAS cells and natural killer T cells (IMM31.CD8Tnk) in EAE, which in healthy WM is primarily influenced by NEU.44.ext without crosstalk between OPC05 and AST (**Fig4H**). In both homeostatic and pathological conditions, we found OPC05-derived ANGPTL2 signal to VE1 and immune cells (IMM and/or MIC), suggesting the requirement of tight regulation involving the immune and vascular systems during myelination and consistent with the unfluctuating OPC05 proportion across WM groups (**Fig3C**). Other than being dismissed from the secretory ANGPTL2 signaling network in EAE, NEU.44.ext, together with NEU.43.ext, communicate with other IMM and AST cells through cell contact-based NOTCH signals (**Fig4I**). Given that NEU43-44 uniquely expresses *VANGL1*, a planar cell polarity gene preferentially expressed in ventricular zones (92), the elevation of *DLL1* (Notch ligand delta-like 1) of NEU43-44 suggests the activation of periventricular neural stem cells (93,94) to control immune cell fate (95).

As independent support for the predicted ligand-receptor relationships, we mapped their spatial colocalization probability at the enhanced subspot level across ME clusters (**Methods**). Consistent with the findings described in Fig2, we found immune-vascular interactions to be enriched in ME group iv and interactions pertinent to glial and immune functions enriched in ME groups ii and iii. Specifically, we found frequent *SELL*-*PODXL* contacts in ME25 (EL WM enriched ME, **FigS5C**), which might mark the early stages of vascular invasion of leukocytes to form new lesions. ME24 (delineating ventricles, **FigS5A**), on the other hand, appears to be the hotspot of secretory signals that attract pDC to ependyma via chemokine (*IL16*), activate astrocytes via cytokine (*TNFSF8*), and alter oligodendrocyte lineage functions via growth factor (*HGF*). While *ANGPTL2* signaling did not restrict to a unique ME, the overall encounter probability increased in IL and PL WM-enriched ME compared to that of NA and Healthy WM or GM (**FigS5D**). Together, these data paint a detailed intercellular interaction map of the evolution of inflammatory, demyelinating WM lesions in primate with unprecedented spatial resolution.

### ▪ Identifying pathological turning points via MRI features: high PD/T_1_ ratio signifies the formation of future lesions

To increase the clinical applicability of our findings, we further explored whether any of the subregional features cataloged here can be identified by MRI, a noninvasive approach that is the standard practice for monitoring MS. First, we quantitatively accessed MRI voxels using the distribution of PDw intensity (roughly proportional to the concentration of hydrogen atoms) and T**_1_** values (longitudinal relaxation time in ms when excited protons return to equilibrium) (**Methods**). We first benchmarked that PD-T**_1_** distribution can differentiate anatomical brain regions; as expected, we saw a clear segregation of WM compared to cortical GM or subcortical GM (subGM) by a crude T**_1_** value cutoff (1250 ms) (**Fig5A**). We then compared PD-T**_1_** distribution across different WM groups and found a gradual change toward higher PD intensity from healthy (He.WM), normal-appearing (NA.WM), to WM lesions (WM.Les) (**Fig5A**). Furthermore, a considerable proportion of voxels within WM lesion presented higher T**_1_** values (>1250 ms).

To further understand the spatial significance of such division, we overlaid two populations back to the terminal MRI image by their coordinates and found that their subregional structures resembled those of the lesion organization identified by spatial transcriptome analysis of the same tissue (**Fig5B**). To further understand the potential of PD-T**_1_** distribution in capturing the WM transition from normal to lesional subregions, we generated 5 concentric rims outward from the T**_1_**-defined lesion core (T**_1_**> 1250 ms). The PD-T**_1_** distribution of the rim5 area (750 μm away from the lesion core) resemble healthy WM, and PD values gradually increase as voxels get approach the lesion core (**Fig5C**). Interestingly, PD values increase earlier along this trajectory than T**_1_** values, and this pattern is common across lesions and animals (**Fig5D**).

Given that MS-like lesions tend to develop centrifugally from their central vein, the lesion core marks the oldest and the lesion edge the most recently damaged areas. Therefore, we investigated whether the changes in signal profile enriched in each subregion can be used to label lesion age — e.g., whether the putatively older lesion (core) corresponds to the lesion core transcriptomic profile (*IGFBP2*^high^/*IGFBP3*^low^) and the newly formed lesion (outer rim) corresponds to the lesion edge transcriptomic profile (*IGFBP2*^high^/*IGFBP3*^high^ or *SERPINE1*^high^/*SERPINB1*^low^) — as validated by findings on longitudinal MRI (**Fig5E**). Indeed, we found that lesions less than 7 days old had a signaling profile resembling that of the lesion edge and that older lesions have a signaling profile more like the lesion core (**Fig5F**). Given that the *IGFBP3* and *SERPINE1* are particularly elevated in OPC09.eae3 and AST10.eae2 at the edge (**Fig5G**) where lesion expansion or containment could occur, targeting these cell types alone or considering their intercellular network collectively (**FigS14**) might be of therapeutic and or diagnostic interests.

Finally, we found that the PD/T**_1_** ratio is a sensitive imaging tool to detect inflammatory events prior to demyelination, with a minimum requirement of manual adjustment in image processing (**Methods**). PD/T**_1_** ratio clearly distinguishes pre-demyelinating subregions (rim1-like area, with high PD but low T**_1_** values, **Fig5C**) from areas with high PD and high T**_1_** values (GM, subGM, and demyelinated WM, **Fig5A-D**). PD/T**_1_** ratio successfully highlights WM regions where future lesions occur (D-42 and D-49) at a time point (D-30) when the pathological changes are not clear on PDw MRI or T**_1_** map alone (**Fig5H**), and/or when the pattern of changes is difficult to distinguish from normal anatomical structures (the putamen in the example shown).

## Discussion

In this study, we performed a cross-modality analysis, joining longitudinal MRI, histopathological features, and single-nucleus/spatial transcriptomic profiling to elucidate the dynamics of MS-like WM lesions in marmoset EAE. Radiologically, we found multifocal lesions across marmoset brain regions, particularly in periventricular WM tracts, recapitulating hallmarks of MS. Histopathologically, we found SB lipid staining unambiguously delineates WM lesions, resembling the morphology of myelin substance imaged by structural MRI. Transcriptomically, we found pathological changes before manifesting myelin destruction, a substantial expansion in the number and diversity of immune/glial cells over time, and distinctive cellular interconnectivity among SB-defined lesional sub-compartments. We found a transcriptomic profile switch within 10 days after lesion formation, concomitant with an elevation of reparative and remyelinating activities at the lesion edge. We identified PD/T**_1_** ratio as a sensitive, noninvasive imaging readout to predict the expansion of demyelinating lesions, which might be applied clinically to track lesion dynamics longitudinally. Considering the three domains of MS management efforts—detect lesions, stop lesion expansion, and repair established lesions—we provide an unprecedentedly detailed molecular map to inform the cellular source of the overlapping pathological and compensational pathways in time and space.

Marmosets are naturally infected with an EBV-related gamma herpesvirus (96) and are exposed to environmental pathogens throughout life, in a manner similar to the way these factors impact the development and aging of the human immune system. These developmental features might predispose marmosets to a hyperimmune response to CNS-derived epitopes inoculated and presented later in life (22). Marmosets can be sensitized by intradermal injection of hMOG/IFA at the dorsal area of the axillary or inguinal lymph nodes, an afferent compartment where T cells are activated before entering the parenchyma (22). Upon entering the targeted compartment of the CNS, T cells interact with glia, recruit monocytes and macrophages, and release cytokines that lead to myelin damage. As phagocytes clear myelin debris into the draining compartment (such as cervical lymph nodes), the new release of myelin epitopes further activates new T-cell specificities, and epitope spreading leads to additional myelin destruction (97). We discuss our findings regarding compartmentalization of autoimmune responses in the following sections.

In NA WM before detectable myelin destruction, T cells, albeit low in nuclei counts, are enriched as a proportion of peripheral immune cells (**Fig3B**). At the same time, widespread glial/vascular responses to demyelination-independent stimuli are apparent. The elevation of structural remodeling genes (*GFAP*, *CALCA*, *BGN*, **Fig2D**) and an increased proportion of transitioning (OPC06 and MIC04, **Fig3C**) and stressed (L2_OLI.eae, **FigS8B**) glial cells underscore the presence of latent components in disease development. Not surprisingly given the high dimensionality of this modality, transcriptome profiling is the most sensitive in our dataset, and pathological activity can be marked by an increased transcriptome complexity in NA WM (**FigS3**). This finding guided our subsequent efforts to develop analysis methodologies for noninvasive measurement (MRI) to account for this latent element in disease monitoring (**Fig5**).

Routes for immune cells to the CNS include the crossing of blood-brain (vessel), blood-CSF (e.g., choroid plexus stroma, meningeal subpial space, post-capillary perivascular space), and CSF-brain (ventricle) barriers (98). T cells enter after recognizing local antigen presentation cells (APC) at brain borders. In line with blood-brain or blood-CSF crossing, we found that monocytes, B cells, and DC are predicted to be in contact with VE and pericytes via *SELL*-*PODXL* signaling (**Fig4D**) in proximity to the central vein of WM lesions (**Fig2G**, **FigS8B**); and with VE and VLMC via VCAM1-integrins signaling in association with other immune cells for extravasation to at sites of inflammation (**FigS13E**). In line with CSF-brain crossing, we found that ependyma increased paracrine signaling and cell-cell interactivities (**FigS12B**) in a manner that is predicted to attract pDC via IL16 chemokine signaling (**Fig4F**). Given that the *IL16* expression level of ependyma is comparable across conditions (**FigS7C**), we interpret this result to suggest that the chemotaxis of pDC occurs following entry of pDC into the CSF space from vessels, facilitated by elevation of VCAM1 (**FigS7C**).

In line with the described reactivation of infiltrated immune cells in the CNS (22,97), we found that natural killer T cells and pDC are predicated to respectively recognize MHC-I and MHC-II expressed by vascular, immune, and glial APC (**FigS13A-B**). Subsequent cellular and humoral responses, including immune/glial cell proliferation, myeloid recruitment, and antibody-(in)dependent complement cascades, encompass the ME of lesion core (**Fig2E-G**, **3C**, **S7**). Additionally, the involvement of perivascular macrophages in Treg homeostasis (**FigS13D**), myeloid-derived HGF in promoting OPC proliferation (**Fig4E**), contact-dependent immune modulation mediated by *VANGL1^+^* periventricular neurons (**Fig4I**, (92,99)), and ependyma-derived CFB (part of the alternative complement pathway, **FigS7B-C**), are events only seen in EAE. Interestingly, we found that *CDKN2A* (encodes p16^INK4a^ and p14^ARF^), a cell senescence marker that inhibits cell division and neural stem cell potential (100), is distinctively expressed by *VANGL1^+^* neurons (NEU43-45, **FigS7B**) and is elevated in the ependyma of EAE (**FigS7C**), suggesting microenvironmental aging in the periventricular zone.

In line with the loss of lipid by SB labeling, we found that genes involved in lipid storage and catabolism (*CD36*, *SLC37A2*, *MSR1*, *NR1H3*, *PLA2G7,* **FigS7**), macrophage-derived foam-cell differentiation (**FigS11D**), and ferroptosis regulatory activities (*AIFM2*, *MGST1, SLC7A11*, *TMEM164,* **FigS7**, (62–65)) are elevated at sites of myelin destruction. Cytokine-mediated oligodendroglial cell death, endoplasmic reticulum stress-induced myelin detachment, and engulfing of myelin debris by phagocytes are all part of inflammatory demyelination and recapitulated in our transcriptomic data. While we did not assess whether or how epitope spreading might impact the lesion dynamics in the current study, we observed multiple waves of demyelination on longitudinal MRI (**Fig5F**), resulting in a discordant formation of lesions over time. Whether this stage-wised myelin destruction is progressively mediated by different T-cell specificities requires further investigation; however, we can leverage this feature and compare lesions of different ages within the same brain by MRI. We discuss the molecular diversity and potential significance of these processes for MS pathogenesis and management in the following sections.

By examining the ME profile of the perilesional WM—where destructive, protective, and reparative signals overlap—we identified a transition from ME8, 10, 12 (comprising inflamed but NA tissue and newly established young lesions) to ME19, 13, 18 (comprising older lesions that developed more than 10 days prior to transcriptomic analysis). As expected, we found that heavy involvement of immune cells is the hallmark of fully developed lesions (ME19, 13); however, EAE-associated astrocytes, OPC, and vascular cells dominate tissue’s transition phases (**Fig2**, **FigS5**). Interestingly, NA ME12 is enriched with astrocyte- and ependyma-derived *NADK2* and *WLS*, which function as metabolic regulators upon increased energy demands (101) and regulate the secretion of Wnt (102), which itself can impact radial glial cell fate (103). These findings suggesting that even early lesions activate of a protective response.

In lesional ME (ME19, 13, 8), genes associated with susceptibility to MS (*HLA-DPB1*, (104)), risk of developing progressive MS (*NR1H3*, (105)), and circulating markers that discriminate chronic active versus inactive MS lesions (*CHIT1*, (106)), are regionally elevated. Moreover, ependyma- and astrocyte-derived *IQCK* (**FigS7A-B**), an AD risk gene associated with Aβ and Tau load in astrocytes (58,61), is uniquely elevated in the lesional ME. While the function of IQCK is unclear, the circular form of *IQCK* transcripts (*circRNA*) is overexpressed in multiple system atrophy, a neurodegenerative disease (107). Given that *circRNA* is often enriched in the secreted exosome of body fluids (108), future studies linking the level of *circIQCK* in MS to astroglial or ependymal activities with liquid biopsy might be of diagnostic interest.

In the young lesional ME (ME8, 10), EAE-associated AST and OPC are the major players (L2_AST.eae, L2_OPC.eae, **Fig2F**). Here, we found elevated senescence-associated secretory phenotype (SASP) and autophagy activities. SASP collectively corresponds to the presence of soluble and insoluble components (growth factors, inflammatory cytokines, proteases, and ECM proteins) secreted by senescent cells (109), which can be positively or negatively regulated by autophagy (110–112). Inducers and members of SASP (*CYR61*, *TNC*, *HBEGF*, *IGFBP3*, *SERPINE1*) expressed by EAE-associated *CHI3L1*^+^ astrocytes and *EVA1A*^+^ OPC were enriched in developing lesions (**Fig2D**, **FigS7A-B**) and are directly or indirectly involved in autophagy regulation (113–118). These findings may be of particular significance for MS-associated pathology, and we discuss it in further detail in the following sections.

Aside from AST and OPC, *CYR61* is uniquely elevated in the ependyma of EAE (**FigS7C**), collectively contributing to leukocyte trafficking and senescence. While discordant results on the inhibitory, permissive, or contradictory roles of tenascins in remyelination and neuroinflammation have been discussed (119), we observed loss of *TNR* expression by homeostatic OPC and gain of *TNC* expression by *CHI3L1*^+^ astrocytes and *EVA1A*^+^ OPC (**FigS7B**). Tenascin-C derived peptide (TNIIIA2) induces p16^INK4a^ and subsequent HB-EGF release, which transforms tissue properties to favor hyper-proliferation and invasive migration (120); additionally, elevation of *HBEGF* in EAE AST and OPC suggests their involvement in neuroprotection (121).

The level of *IGFBP3*, regulating IGF-1 bioactivity in circulation and inducing senescence (122), is enriched in the EAE-associated oligodendrocyte lineage (OPC07-12, OLI09-11). Moreover, *SERPINE1*, counteracting the tPA-mediated inhibition of IGFBP3 (123), is unique to a subcluster of *CHI3L1*^+^ astrocytes (AST10.eae2) of young lesions (**FigS7B**). Surprisingly, we found reduction of *IGFBP3* accompanied by elevation of *IGFBP2* (another IGF-1 regulator) as lesions aged (**Fig5E-F**). While the succession of IGF-1/IGFBP levels in body fluids of MS and their clinical relevance has been reported (124–128), the glial source of such senescence markers associated with lesion activities was not previously recognized, and our results suggest that AST10.eae2 is the upstream regulator of the IGFBP-mediated SASP cascade. Indeed, we found that AST10.eae2, the most dominant astrocyte subtype in Gd-enhancing lesions (**FigS10B**), undergoes the most drastic changes in ECM-cell signaling relative to all other astrocytes subtypes across conditions (**Fig3E**). AST10.eae2 is a distinct subtype from the previously described AIMS (astrocytes inflamed in MS in chronic active lesions, (129)) as it does not express classical complement components (*C1S*, *C1R*); instead, it expresses components and regulators of lectin (*MASP1*) and the alternative (*CFI*) complement pathways, which are not triggered by antibody recognition (**FigS7B**).

Interestingly, other than the SASP members enriched in young lesional ME, genes (*CHI3L1*, *EVA1A*) that distinguish EAE-associated astrocytes and OPC from their homeostatic counterparts can induce autophagy (130,131), a process that collaborates with apoptosis pathways to control oligodendrocyte number (132). Similarly, *ANGPTL2*, a secretory pathway between differentiating OPC-immune/vascular cells (**Fig4H**), is another SASP molecule (133) that regulates autophagy (134). Autophagy is required for removal of cytoplasm to promote oligodendrocyte development and myelin compaction (135,136) and generates a permissive environment for remyelination (137). In agreement with the putative beneficial role of autophagy in PL WM, we identified a remyelinating ME18 that increases in proportion as lesion age (**FigS7D**), highly expresses differentiating OPC genes (*TNFRSF21*, *BCAS1*, *SERINC5*, *RHOQ*, *ENPP6*), and resides in proximity to areas with elevated expression of autophagy genes. Together, these findings pinpoint the spatiotemporal features of a regulatory network with implications and potential targets for the therapeutic promotion of remyelination.

### Pitfall and limitations

While the animal model and approaches employed here improve our understanding of lesion dynamics in some respects, the sample sizes of the dataset need to be expanded to query sex-, age-, and region-specific responses to EAE. Additionally, while the SB/NFR staining is sensitive enough to identify WM lesions, it has limited ability to identify the location of GM lesions, which makes a targeted analysis of cortical and subcortical lesions, which are also formed in marmoset EAE (18), challenging in the current analysis. Another limitation of our analysis is that, despite the reported clinical, radiological, and immunological similarity of CFA- and IFA-induced marmoset EAE (23), the WM groups comprised of older (>1000 days) or resolved lesions in the current dataset were exclusively derived from CFA-induced marmosets. Thus, additional experiments on EAE samples induced by our newer hMOG/IFA protocol are required to corroborate the current findings in aged and resolved groups. Finally, given that the temporal resolution of our longitudinal MRI scans was limited to 7 days to allow the animals to recover sufficiently from anesthesia, a different imaging approach would be required to date early lesions more precisely in order to capture their rapidly changing dynamics.

In conclusion, our comprehensive clinical, radiological, and single-cell and spatial transcriptional characterization of lesion development and repair in marmoset EAE identifies region- and stage-specific microenvironments and summarizes the sequence of events in the evolution of MS-like lesions. We found a distinct type of astrocyte and OPC reactivity that may comprise the earliest macroglial response to inflammatory demyelination. We leveraged our multimodal data to develop an image-based approach to detect impending lesions. Our findings implicate a wealth of molecules with diagnostic and therapeutic potential, particularly in the space of neuroglial protection and repair, and point to ways of developing circulating biomarkers, molecular MRI, and preclinical trial designs that could have implications for therapy development in MS.

**Table 1.**
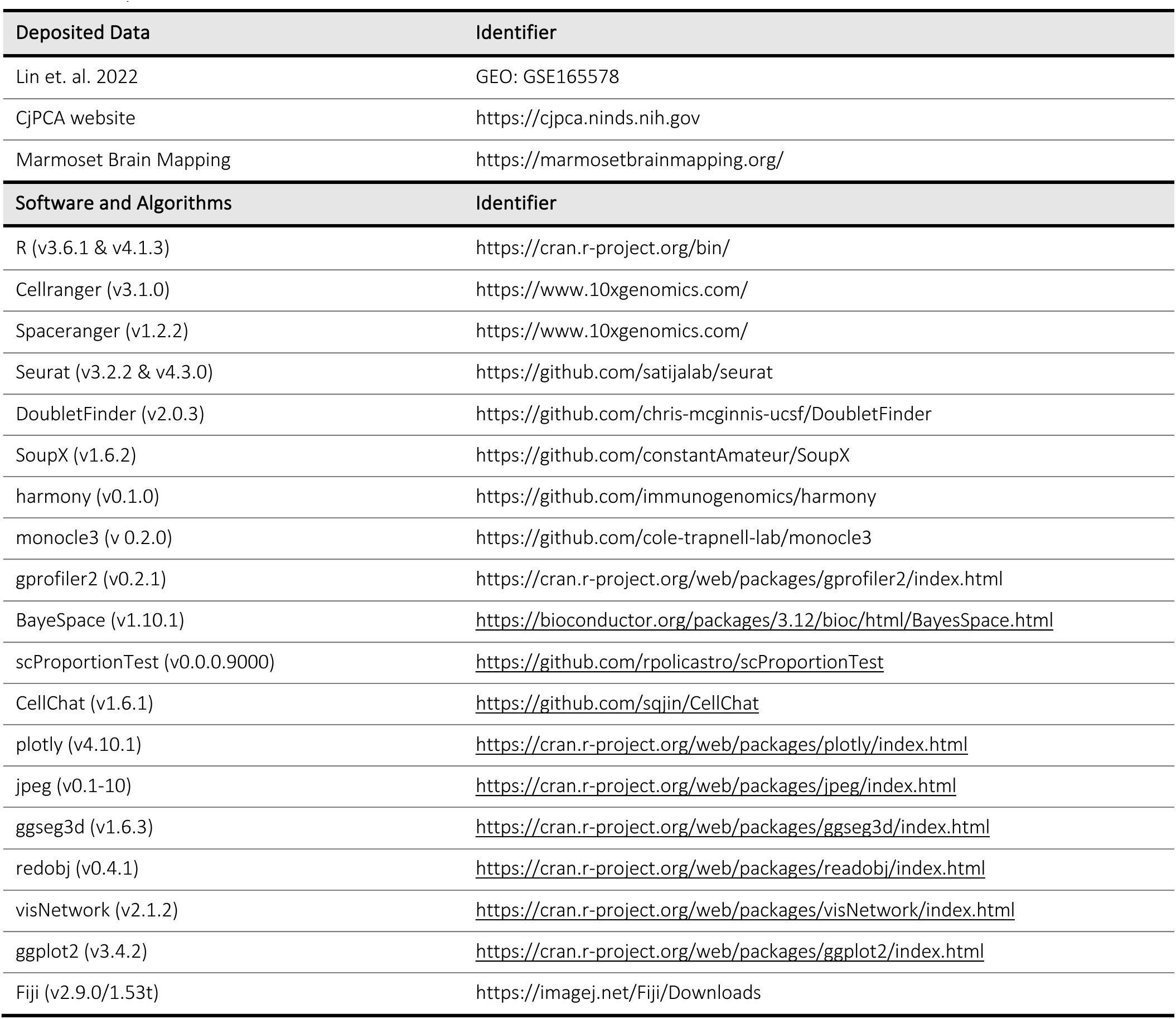
Key resources.

## Methods

### Animal EAE induction and MRI acquisition

All marmosets were housed and handled with the approval of the NINDS/NIDCD/NCCIH Animal Care and Use Committee. Before each experiment, marmosets will be neurologically examined with an expanded disability status scale (mEDSS) developed for marmoset EAE (30) to track the progress of clinical symptoms. To perform in vivo MRI scans, marmosets were given atropine sulfate (0.04 mg/kg, NDC 0641-6006-01, West-Ward Pharmaceuticals) and ketamine (10mg/kg) by intramuscular injection before being intubated (2.0 mm Endotracheal Tube Sheridan® Uncuffed™, 5-10404, McKesson Corporation) and ventilated with a mixture of isoflurane (2-4%) and oxygen. Gadobutrol (0.2 mmol/kg, Gadavist, NDC 50419-325-02, Bayer HealthCare Pharmaceuticals) in 3mL Lactated Ringer’s solution (NDC 0990-7953-02, ICU Medical) was injected intravenously through a catheter (24G x 3/4", Terumo Medical Surflash Polyurethane IV Catheter, SR-FF2419) and extension set (29" Male LL Adapter, 2C5645, Baxter) slowly over 2 min to identify newly inflamed demyelinated lesions (denoted as Gd.Lesion). A marker (LiquiMark MRI Markers, LM-1, suremark) was placed at the right hemisphere of the brain to identify image orientation. MRI was performed on a 7 Tesla scanner (Bruker, Biospin). Acquisition protocol included a proton density-weighted sequence (PDw; TE = 16 ms; TR = 2300 ms; number of acquisition = 3; spatial resolution = 0.15 x 0.15 x 1 mm^3^; matrix = 213 x 160 x 36; acquisition time = 23 min total) as a qualitative structural image highly sensitive to brain abnormalities and a T**_1_**-Magnetization-Prepared 2 Rapid Acquisition Gradient Echoes sequence (T**_1_**-MP2RAGE; TE = 3.5 ms; TR = 6000 ms; inversion time (TI) *1* = 1200 ms; TI*2* = 2000 ms; flip angles = 7°/8°; number of acquisition = 4; spatial resolution = 0.15 x 0.15 x 1 mm^3^; matrix = 213 x 160 x 36; acquisition time = 28 min total) (138). Quantitative values related to tissue properties, called T**_1_** values (in milliseconds) were extracted from this sequence for each voxel using a pipeline in Matlab (138). Resulting images are called T**_1_** maps and were generated for each animal and each time points. After each MRI scan, marmosets were weaned from isoflurane, recovered with a warmed lactated ringer injection subcutaneously, and returned to their original housing.

The transcriptomic map was generated from 11 (3 control and 8 EAE) 4–11 years old common marmosets (*Callithrix jacchus*), 4 males (CJR02, CJR05, CJH11, CJJ12) and 7 females (CJH01, CJP03, CJG04, CJM07, CJP08, CJH09, CJM10). EAE marmosets received 200mg human white matter (hWM, prepared from patient donors) or 200μg recombinant human myelin oligodendrocyte glycoprotein (hMOG 1–125, AS-55158-1000, AnaSpec) emulsified in complete (CFA; Difco Adjuvant, 231141, BD) or incomplete Freund’s adjuvant (IFA; Difco Adjuvant, 263910, BD) in a 1:1 volumetric ratio (See **FigS1-2** and **TableS1** for details). To generate homogenate with desired consistency, components of immunogen were triturated ∼50 times in an enclosed device. The device consisted of two 5-mL syringes (Luer-lok syring, 309646, BD) and one 3-way stopcock (Discofix® 3-way Stopcock, B.Braun Medical Inc). A total of 200 μL homogenate was injected into 4 dorsal spots around the lymph nodes.

### Tissue dissection for cryosection and nuclei isolation

Before the day of tissue harvest, a custom-made brain holder was generated for each marmoset brain by 3D printing (Ultimaker 2^+^) to guide tissue sampling (139). Tissue dissection was carried out as described (42). Briefly, marmosets were deeply anesthetized with 5% isoflurane until no signs of breathing, then were transcardially perfused with ice-cold artificial cerebrospinal fluid (aCSF) for 5 min. After skull and meninges removal, each brain was quickly positioned into the designated brain holder submerged in ice-cold aCSF solution within 10 min post-perfusion. The brain was sectioned into 12–13 slabs at 3 mm with a homemade blade-separator set submerged in aCSF solution. Each brain slab was transferred into a tissue cassette (70078-15, Electron Microscopy Sciences) with the anterior side of the brain slab facing the biopsy foam pad (62325-06, Electron Microscopy Sciences). The cassettes were then submerged into a jar full of RNAlater (RNAlater™ Stabilization Solution, AM7021, Invitrogen) and stored at 4°C overnight. The following day, brain slabs were transferred to 25 x 20 x 5 mm^3^ molds (Tissue-Tek® Cryomold®, 4557, Sakura Finetek) on ice to facilitate targeted sampling. Slabs were matched to terminal MRI and informed by marmoset 3D MRI atlases (140,141) for region annotation. For nuclei dissociation, a cylinder of tissue 2 mm in diameter and 3 mm in height for each area was collected with a tissue punch (EMS-core sampling tool, 69039-20, EMS), and ejected into PCR tubes filled with 100 µL of RNAlater and kept at −80°C for long-term storage. For cryosection, a 6.5 x 6.5 x 3 mm^3^ block of tissue per region of interest (ROI) was prepared by a customized 3D-printed brain cutter and stored in cryomold filled with RNAlater at −80°C or proceeded immediately to OCT (Tissue-Tek® O.C.T. Compound, 4583, Sakura) embedding. The quality of RNAlater-preserved tissue was assessed by measuring RNA Integrity Number (RIN) on the Agilent 2100 Bioanalyzer (G2939BA, Agilent). Bulk RNA was isolated with TRIzol™ Reagent (15596026, Invitrogen) and measured with Agilent RNA 6000 Pico Kit (5067-1513, Agilent); samples with RIN >8.5 were used in the study.

### Tissue block and library preparation for spatial transcriptomic (ST) analysis

RNAlater preserved tissue blocks were thawed at room temperature (RT), retrieved from the cryomold with a pair of RNase AWAY (surface decontaminant, 7000TS1, ThermoFisher) treated forceps, dabbed with clean Kimwipes to remove excess liquid, and incubated in a new cryomold filled with OCT at RT for 10 min to further remove RNAlater residual. At the end of incubation, tissue blocks were transferred to a new cryomold filled with OCT, positioned to desired orientation without creating bubbles, and left frozen at −20°C before trimming. Frozen OCT blocks were trimmed to desired size (∼8 mm on each side) to reduce the chances of tissue folding during serial sectioning. To increase the precision of tissue capture, a cryo-resistant plastic grid with matching capture areas was attached to the back of each Visium Spatial Gene Expression Slide (2000233, 10x Genomics) and 12 microslides (Superfrost^+^ and ColorFrost^+^, EF15978Z, Daigger Scientific). Gird-attached slides and tissue blocks were left equilibrated inside the cryostat chamber for 30 min prior to a serial tissue scan, which was performed to ensure the capture of the desired ROI with matching morphology to the terminal MRI. Specifically, tissue blocks were sectioned at 10 µm, captured with microslides, and stained every 120 µm with 3% Sudan black (199664-25G, Sigma-Aldrich) in ethylene glycol (BDH1125-1LP, VWR) solution at 56°C for 30 sec in a water bath to color myelin. After Sudan black (SB) stain, microslide was rinsed with running tap water for 1 min, transferred to hematoxylin (100% Surgipath SelecTech Hematoxylin 560MX, 3801575, Leica) at 56°C for 30 sec to color gray matter. After hematoxylin stain, microslide was rinsed with running tap water again for 1 min, coverslipped (Premium Cover Glasses, EF15972L, Daigger Scientific) with UltraPure^TM^ glycerol (15514-011, Invitrogen), and visualized with a Brightfield microscope (Leitz Laborlux S Wild GMBH). When the morphology of the tissue section meets the desired target, the adjacent section will be captured and mounted onto the Visium Spatial Gene Expression Slide without covering the fiducial frame to facilitate image alignment in the downstream data analysis. In parallel, pre- and post-Visium captured tissue sections were made and kept at −20°C for long-term storage.

At the end of tissue scans, Visium Spatial Transcriptomics slides containing the tissue sections of interest were transported on dry ice and kept at −80°C for long-term storage or immediately proceeded to staining and library preparation. The slide was thawed on a pre-warmed Thermocycler Adapter (3000380, 10x Genomics) at 37°C for 1 min to minimize tissue damage caused by condensation. To stain myelin, 1 mL of 1% SB/ethylene glycol was pipetted directly onto the leveled tissue slide and incubated for 5 minutes at RT. After removing excess SB solution, stained slide was dipped in 50 mL of ddH**_2_**O (351-029-131CS, Quality Biological) in a falcon tube for 5 times, dipped in 800 mL of ddH**_2_**O 15 times, and another 15 times in a separate glass beaker containing 800 mL of ddH**_2_**O. To get an optimal contrast between myelin and gray matter, tissue was dried for 1 min (the time should be extended if the tissue section is not dry) before being stained with 1 mL Nuclear Fast Red (NFR, ab246831, Abcam) for 10 minutes at RT at a leveled surface and rinsed again as described for SB staining. To avoid bubbles, slides were first saturated with 5 mL of ddH**_2_**O on a leveled surface and gradually replaced with 80% glycerol solution containing 5% RNase inhibitor (AM2684, Thermo Fisher Scientific) through steady vacuuming off ddH**_2_**O from one end of the slide and adding glycerol with a pipette at the opposite end simultaneously. At the end of solution swapping, the homogenous mounting media was coverslipped and proceeded immediately to 4X tiling imaging with a Nikon Eclipse Ci microscope. After imaging, the coverslip was rinsed off by submerging the slide in 3X SSC buffer (46-020-CM, Corning) diluted in ddH**_2_**O, then rinsed briefly in 1X SSC buffer to remove excess mounting media.

After imaging and coverslip removal, the slide was enzymatically permeabilized with Visium Spatial Gene Expression Reagent Kit (PN-1000184, Spatial 3’ v1, 10x Genomics) at 37 °C for 20 minutes, which was determined by Tissue Optimization protocol (PN-1000193, 10x Genomics). We then prepared cDNA library by following the manufacturer’s protocol (Visium Spatial Gene Expression Reagents Kits User Guide, Rev D). The cycle number for cDNA amplification for each library (**TableS2**) was determined using the Cq value obtained from the qPCR steps detailed in the manufacturer’s protocol. All libraries were sequenced using the Illumina Novaseq S2 platform. Library quantity and quality were assessed using Qubit™ dsDNA HS Assay Kit (Q32854, Invitrogen™) with a Qubit™ 4 Fluorometer (Q33226, Thermo Fisher Scientific) and High Sensitivity DNA Kit (5067-4626, Agilent Technologies) with a 2100 Bioanalyzer instrument (G2939BA, Agilent Technologies), respectively.

### Single-nucleus dissociation and library preparation for RNA sequencing

Nuclei preparation was carried out as described (42). Briefly, RNAlater preserved samples were thawed on ice, retrieved from the storage tube with a pair of clean forceps, dabbed with Kimwipes to remove residual RNAlater, and placed in a 1 mL douncer tube (Dounce Tissue Grinder, 357538, Wheaton). Each tissue was homogenized in 500 μL of lysis buffer containing 400 units of RNase inhibitor (RNaseOUT Recombinant Ribonuclease Inhibitor, 10777-019, Invitrogen) and 0.1% Triton-X100 in low sucrose buffer (0.32 M sucrose, 10 mM HEPES, 5 mM CaCl**_2_**, 3 mM MgAc, 0.1 mM EDTA, and 1 mM DTT in ddH**_2_**O, pH8) with loose pestle 25 times and tight pestle 10 times. Additional 5mL of low sucrose buffer was used to rinse the douncer, and the homogenate was filtered through a 40-μm mesh (Falcon® 40 µm Cell Strainer, 352340, Corning) to a 50-mL Falcon tube on ice and homogenized at a speed of ∼1000 rpm for 5 sec to brake nuclei clumps with a handheld homogenizer (VWR® 200 Homogenizer). After homogenization, 12 mL of high sucrose buffer (1 M sucrose, 10 mM HEPES, 3 mM MgAc, and 1 mM DTT in ddH**_2_**O, pH8) was placed underneath the lysate with a serological pipet by gravity and set on ice. Without disturbing the low-high sucrose interface, the Falcon tube was capped and placed in a swing bucket to be centrifuged at 3,200 rcf for 30 min at 4°C. At the end of spin, the supernatant was decanted quickly without tabbing, and 1 mL of resuspension buffer (0.02% BSA in 1X PBS, pH7.4) containing 200 units of RNase inhibitor was added to rinse off the nuclei. Nuclei were rinsed off the wall in courses of 2 sec per trituration for 20 times total per tube, the Falcon tube was then capped and spun at 3,200 rcf for 10 min at 4°C. At the end of the spin, tubes were gently tabbed to remove any visible supernatant and collected nuclei with 200 μL resuspension buffer. The nuclei suspension was filtered through a 35-μm mesh (Cell Strainer Snap Cap, 352235, Corning) twice and counted on a hemocytometer by trypan blue staining. During counting, the size and quantity of myelin and other debris were visually inspected under the scope, and the suspension was filtered 1–3 more times through the 35-μm mesh if necessary. Only round and dark-blue stained nuclei were considered of good quality and included in the final count.

The snRNA-seq dataset of the brain analyzed in this study consists of 43 libraries (26 of them were newly prepared from EAE animals, and 17 of them were from naïve animals and have been reported in Lin et. al, 2022, GSE165578). However, it is important to note that EAE samples were processed in the same batch with matching controls in tissue location or diseased conditions whenever possible (**FigS1-2**, **TableS3**). All libraries were prepared using 10x Genomics Chromium Single Cell 3’ Library & Gel Bead Kit v3 following the manufacturer’s protocol. Briefly, nuclei suspensions were diluted with resuspension buffer as described above at desired concentration and loaded into Chromium Controller to generate droplet emulsion. Twelve cycles were used for both cDNA amplification and library sample index PCR, and sequenced on Illumina Novaseq S2 (39 libraries) and Hiseq 4000 (4 libraries), according to the manufacturer’s protocol (**TableS3**).

### Single nucleus transcriptomic data analysis pipeline

*Alignment.* The raw snRNA-seq reads were aligned to a marmoset genome assembly, ASM275486v1 (GCA_002754865.1), with a reference package built as described in (42). CellRanger (version 3.1.0, 10x Genomics) software was used to align reads for snRNA-seq samples to generate cell barcode-to-gene feature matrix for downstream analysis, and automatic estimation of cell number was applied for most of the snRNA-seq samples unless otherwise specified (**TableS3**).

*Preprocessing and quality control*. Preprocessing and quality control parameters were applied as described in (42). Briefly, Seurat v3 object was created for individual samples and DoubletFinder (142) was applied to estimate and remove putative doublets to mitigate technical confounding artifacts. In addition, cells with gene numbers 200– 5000 and less than 5% mitochondrial genes were kept, and genes observed in more than 5 cells were kept. In parallel, SoupX (143) was applied to correct ambient RNA background, through which ambient RNA from empty droplets that contained <10 unique molecular identifiers (UMI) were analyzed, and the “soup” contamination fraction was calculated and removed for each cluster. Next, the cell barcodes that passed the DoubletFinder and additional QC were used as index to subset the SoupX-corrected matrix to generate a new matrix subset as downstream analysis input. For individual samples, post-QC Seurat object was created, and the index labels: IL01_uniqueID, IL02_species, IL03_source, IL04_sex, IL05_ageDays, IL06_tissue.1 (coarse category), IL06_tissue.2 (developmental category), IL06_tissue.3 (fine category), IL07_location, IL08_condition (diseased condition), IL09_illumina, IL10_chemistry, IL11_batch, IL12_lMinDays, IL13_lMaxDays, IL14_dataset, IL15_annotation were added to the metadata as cell attributes.

*Clustering and visualization.* As described in (42), hierarchical level 1 (L1) and level 2 (L2) analyses were employed for snRNA-seq dataset to facilitate cluster tracking and result interpretation. Briefly, a merged Seurat object was created from 43 snRNA-seq samples, log-normalized and scaled, and the top 3000 variable genes were calculated and used in Principal Component Analysis (PCA). Harmony (144) was applied to integrate different samples over IL01_uniqueID attribute, and the top 5 Harmony-corrected embeddings (H5) were used for Seurat to learn UMAP and annotate L1 cell classes. Canonical cell-type markers (*PTPRC* for immune cells, *PDGFRA* for OPC, *MAG* for oligodendrocytes, *GFAP* and *SLC1A2* for astrocytes, *LEPR* and *CEMIP* for vasculature and meningeal cells, and *CNTN5* and *NRG1* for neurons) annotated 6 of the classes unambiguously. Low-quality cells that got high percentage of reads mapped to the mitochondrial genome, low RNA counts and features, and/or expressed genes that mapped to multiple cell class canonical markers were removed. Nuclei that passed L1 QC were divided into 5 partitions (IMM, OPC, OLI, VAS/AST, NEU) for L2 analysis. Additional rounds of QC were applied to IMM and VAS/AST partitions prior to L2 analysis to facilitate artifact identifications among similar cell classes. IMM partition was further split into *FLT1*^high^ microglia (MIC) and *FLT1*^low^ peripheral immune cells (P.IMM) cell classes, and VAS/AST mixed partition was separated into *ALDH1L1*^high^ astrocytes (AST) and *ALDH1L1*^low^ vascular/meningeal/ventricular cells (VAS). As a result, a total of 7 partitions (AST, OPC, OLI, MIC, P.IMM, VAS, NEU) were parsed for L2 analysis. As detailed in (42), rounds of supervised QC, differentially expressed gene (DEG) search, and unsupervised clustering were performed to yield a total of 133 subclusters in this study. The following compound naming conversion was created to label subclusters: general cell class category in numeric order, major tissue type or diseased condition contributor for each subcluster. If a subcluster found in the current report is similar to the transcription profile of a subcluster reported by (42), the same numerical name is followed.

*Preparation of objects for cross-cluster and analysis.* To facilitate downstream analysis and comparison, several annotated object subsets were created after the subclustering and UMAP embedding were finalized for each cell class. An object (EAE200) containing up to 200 nuclei per cluster for all 133 subclusters was prepared by random sampling to facilitate global and local unique gene selection and comparison. An object (EAEwm) containing nuclei sampled from “WM” (fWM, tWM, pWM, pCC, and OpT) was prepared to analyze the relative prevalence of each subcluster across WM lesional states to create the centered and z-scored heatmap (**Fig3B**). The same EAEwm object was used to analyze the dominant subclusters during lesional states transition using scProportionTest (v0.0.0.9000) package. To infer and compare cellular interactions at the WM between control and marmoset EAE with CellChat (145), a nuclei number-balanced object (EAEwm200) between naïve control and EAE samples was prepared from the same “WM” sampling sites stated above to mitigate outlier biases. More specifically, subclusters within each condition with lower than 50 nuclei were disregarded from further analysis, clusters with many nuclei were down-sampled to 200, and an equal number (50–200) of nuclei were sampled from the matching subcluster found in control and EAE samples.

### Spatial transcriptomic (ST) data analysis pipeline

*Alignment, data pre-processing, and image correction*. The raw ST reads were aligned to the same reference package as for the snRNA-seq dataset using SpaceRanger (version 1.2.2, 10x Genomics) software to generate spot barcode-to-gene feature matrix for downstream analysis. Individual ST object was created by Load10X_Spatial() function with Seurat (v3.2.2) package (146) and normalized with SCTransform() function (147). The merge() function was used to aggregate 16 slices into one ST object (Visi) containing images, and the Visi@assays$SCT@counts was pulled to create a separate ST object (VisiDot) by CreateSeuratObject() function to get aggregated spots without images. A processing pipeline similar to that of snRNA-seq was employed for the VisiDot object; specifically, Harmony() was applied over “IL01_uniqueID” variable to integrate 16 samples. To match the terminal MRI images, the orientation of the Visium images was corrected by transforming the image array stored at Visi@images$slice1@image, and the spot information stored at Visi@images$slice1$@coordinates was swapped accordingly (See GitHub post for detail). The microenvironment (ME) cluster annotated in the VisiDot object was transferred back to the Visi object for spatial visualization.

*ST Gene module analysis and gene ontology (GO) analysis.* As described in (42), Monocle3 (148) was used to group genes in the Visi object (Visi@assays$Spatial@counts[rownames(Visi),]) into modules by their similarity along the learned neighbor (Knn) or principal (PG) graph. The score of the gene list was then calculated and added to VisiDot object by AddModuleScore() function and visualized in Seurat (**FigS6**). For each gene module, GO analysis was performed by the gost() function with gprofiler2 (v0.2.1) package using “cjacchus” database, including electronic GO annotations (IEA), and applying g:SCS for multiple testing correction. Terms that passed a significance cutoff (*p*=0.05) after correction were kept, and the fold enrichment was calculated as follows: (intersection_size/query_size)/(term_size/effective_domain_size).

*Lesion rim assignment and subregion analysis with ST image processing pipeline.* To annotate Visium spots by their histological and lesional features, several masks were created (**FigS4B**). Specifically, the myelinated WM area (SB^+^) was extracted (Colour_3 channel) from the SB/NFR image of Visium slide by “Colour Deconvolution” function in Fiji software (v2.9.0/1.53t, ImageJ2) with the default "H DAB" setting. The Colour_3 channel was then thresholded to create SB^+^ WM binary mask. After reorientating the SB^+^ WM mask to match the corrected Seurat object as stated above, the coordinates of the mask were exported and transferred and transferred to the 10x Visium hexagon coordinate system (Visi@images$slice1$@coordinates) by dilatating each SB^+^ pixel using 8-neighbor model 5 times (resulting ∼10 µm expansion in diameter). To further distinguish SB^-^-gray matter (GM) from SB^-^-demyelinated WM area, GM and lesion gene module scores were calculated and filtered to create GM and lesion masks accordingly. Specifically, EAE200 object was used to find DEG over the coarse tissue category (IL06_tissue.1 label, “WM,” “other,” and “GM”) with Seurat using the Wilcoxon Rank Sum test. Significantly (adjusted p-value < 0.05) enriched (average log fold change > 0.25) DEGs detected in lower than 10% (pct.2 < 0.1) of the other population were kept for enrichment score calculation. Filtered DEG gene lists for “GM” and “other” were combined and added to the Visi object with AddModuleScore() function and termed as “ModuleScore_EAE.otherGM;” similarly, filtered “WM” DEG module scores were added and termed “ModuleScore_EAE.WM” (See **TableS8** for gene list). The ModuleScore_EAE.otherGM and ModuleScore_EAE.WM scores >0.01 and >0.1 were filtered to create the GM and Lesion masks correspondingly (**FigS4B**). Next, spots that exhibited both SB^-^ and Lesion^+^ signals were identified as the lesion core, and 10 concentric rims (SB^+^WM_rims) extending outward from the lesion core (coordinates [x±1, y±1] and [x, y±2]) were assigned to mark the adjacent lesional neighborhoods. The normal-appearing (NA) WM area was annotated by subtracting the lesional neighborhoods from the SB^+^ WM mask in animals with experimental autoimmune encephalomyelitis (EAE), and this region was labeled as “SB^+^WM_NA.Ctrl." Additionally, lesional neighborhoods that overlapped with the GM mask were labeled as "SB^-^notWM_rims," while the supplemental area was labeled as "SB^-^notWM_EAE" in animals with EAE. Furthermore, the subregions within the lesion core were further divided based on centripetal rim assignments (SB^-^WM_-rims) with the same strategy as stated above. For healthy animals, "SB^+^WM_He.Ctrl" and “SB^-^notWM_He" labels were used to annotate tissue with or without SB staining, respectively. Regional differentially expressed genes (rDEG) across assigned subregions are calculated with Seurat.

*ST resolution enhancement.* To facilitate ST data exploration, several strategies were employed to increase the spatial resolution aiding the recognition of patterns and the deconvolution of cell types. First, BayesSpace (v 1.10.1) package was utilized to infer an enhanced transcription expression map to near single-cell level per subspot. Specifically, readVisium() function was applied to create SingleCellExperiment (Sce) object for each sample from the output of SpaceRanger. Sce object was pre-processed with sptialPreprocess() function that log-normalized the counts, analyzed PCA on 2000 variable genes, and kept the top 7 PCs. After applying qTune() function, parameters (q=9, d=7, model="t", gamma=2, nrep=1000, burn.in=100, jitter_prior=0.3, jitter_scale=3.5) were used for spatialEnhance(), and enhanceFeatures() function was used to predict expression for all genes to create enhanced Sce object (Bayes). The inferred expression matrix was then retrieved by “logcounts(Bayes)” function to create enhanced Seurat Spatial object (Bayes2Seurat) for visualization and rescaled to 1 for downstream processing. Specifically, “Bayes2Seurat@images” was swapped with a reconstituted object containing the original tissue image (“Seurat@images$slice1@image”) and enhanced indexes pulled from “colData(Bayes).” The image and indexes of Bayes2Seurat object were further reoriented as described above to match the terminal MRI. To annotate the subspot, scores of rescaled values were averaged and compared for selected genes among lists and assigned to each subspot by similarity (**FigS4C**).

### CellChat intercellular communication analysis

As described above, the nuclei number-balanced EAEwm200 object was used to analyze intercellular communication with CellChat (145) package (v1.6.1). The CellChat database (CellChatDB.human) contains 3 categories of interactions: secrete autocrine/paracrine signaling interactions (secreted–cell), cell-cell contact interactions (cell–cell), and extracellular matrix (ECM)-receptor interactions (ECM–cell). Cell-cell communications among nuclei subclusters residing in control and EAE WM were calculated separately using CellChat pipeline and then compared between conditions.

The incoming/outgoing signal strength and interaction category among subclusters of each condition were compared (**FigS12**), Cleveland Dot Plots (**Fig3E**) were created from the output of netAnalysis_signalingRole_scatter() function (**FigS10**). Lists of cellchat@net$prob, cellchat@net$pval, and cellchat@netP$pathways were filtered for significance (interaction probability > 0 and p-value < 0.05), and the similarity of ligand-receptor (LR) pairs and inferred signaling pathways were compared. Specifically, LR pairs utilized by the same pairs of subclusters in both conditions were placed in the “share” category; otherwise, unique LR pairs or among unique pairs of subclusters to each condition were placed in other bins accordingly (**Fig4A**).

The output of rankNet() measuring “weight” and “count” was utilized to create the Dot Plot summarizing the prevalence and strength of signaling pathways between conditions (**Fig4B**). For selected pathways, Chord Diagrams were created by netVisual_chord_cell() function to visualize the subcluster identities for each interaction (**FigS13**), and signal networks were created with visNetwork package (v2.1.2) to better-summarizing signaling roles (**Fig.4C– I**, **FigS13**). Specifically, the output of netAnalysis_signalingRole_network() and netVisual_heatmap() were utilized to create signal networks to identify dominant senders (out-degree), receivers (in-degree), mediators (flow betweenness), and influencers (information centrality). The output of netAnalysis_signalingRole_network() was acquired by adding “return(ht1)” in the code to modify the function. The output of netAnalysis_contribution() was used to generate Dotplot comparing the contribution of each LR pair within a selected signaling pathway (**FigS14**), the relative contribution per pathway over each condition is visualized as dot size.

The LR pair encounter probability in subspot resolution was quantified by counting the number of overlapping subspots with scaled expression > 0.15 (Bayes2Seurat objects) of targeted genes in the LR pair over total subspots across ME labeling and visualized as Pie Charts (**Fig4J**). To plot the expression of LR pairs, LR pair index was updated for cases that protein or complex names do not match with gene names, such that VEGFR1 was swapped to *FLT1*, VEGFR2 to *KDR*, NPNT1/NPNT2 to *NPNT*, *FRAS1*, *FREM1*, *FREM2*.

### MRI characterization of MS-like lesions in the Marmoset Quantitatively (M3Q) image processing pipeline

*WM lesion age estimation* 3D-PDw images taken from longitudinal MRI follow-ups of animals (CJM07, CJP08, CJH09, CJM10, CJH11) were post-processed to retrospectively date lesions (**Fig5F**). Briefly, a whole-brain mask was extracted from baseline image for each animal using the Algorithms > Brain tools > Anonymize > “Remove skull” function in MIPAV software (v11.0.3). Images from all time points were subjected to N4 bias field correction (149). Then, skull-removed baseline and N4-corrected images were imported to Fiji software for further processing. Skull residual areas remaining from the skull-removal function were manually removed (“Paintbrush Tool”) if necessary. Decks of binary masks (Image > Adjust > “Threshold”) and ROI (Edit > Selection > “Create Selection”) were created (Add to Manager) for image alignment. N4-corrected images of different time points were manually aligned for in-plane (image > Transform > “Translate”) and Z (image > Stacks > Tools > “Stack Sorter”) positions and to the standardized masks created from baseline, and then the inversed masks (Make Inverse) was subtracted from each image and saved as a 3D image deck. Next, the skull-removed and aligned 3D images of each time point were imported into MIPAV to create 4D images (Utilities > 4D volume tools > “Concat Multiple 3D to 4D”) in chronological order to facilitate lesion dating. The gaps in days between MRI scans were calculated, which gap is the maximum age of a lesion if noted only in the later observation point. In parallel, image decks for each brain slice across time points were then created (Utilities > 4D volume tools > “Swap dims 3<–>4”) for downstream processing.

#### WM lesion load quantification

To quantify lesion load, skull-removed and aligned images from baseline and terminal time points from all animals were individually registered to the matching slice acquired from the marmoset MRI atlas (140,141) using Fiji software (Plugins > Registration > “bUnwarpJ”), and the matrix of intensity was extracted (Analyze > Tools > save XY Coordinates) from the registered images. The XY coordinates of brain labels grouped as cortical GM, subcortical GM, and WM tracts were pulled by the same method and indexed onto the matrix of registered images with R. Whenever possible, Macro.ijm codes created by text rendering with R from a template created by Fiji (Plugins > Macros > “Recorder”) were applied to automate the pipeline stated above. The intensity of each PDw image was normalized to the median intensity of voxels labeled as cortical GM referenced in the atlas, and the lesion mask was created by subtracting the normalized PDw image at the terminal from the baseline and binary filtered (FigS4A). Voxels in the lesion mask referenced as WM in the atlas were pooled across animals to calculate the prevalence of WM lesions across brain regions. The probability of a voxel being identified as a lesioned hit was quantified by summarizing the results stated above across 5 EAE animals, and the percentage of such probability was visualized for each WM area and categorized by tracts with R (**Fig1B**). To visualize the distribution of WM lesions across the whole brain, WM lesion masks were overlaid onto a 3D brain shell created from the atlas. Specifically, decks of brain and lesion masks were scaled to 1 x 1 x 6.67, OBJ 3D geometry files were created with Fiji (Plugins > “3D viewer”), rendered in R with readobj (0.4.1), ggseg3d (v1.6.3), and plotly (v4.10.1) packages, and saved as an interactive HTML file format.

#### Imaging biomarker exploration for lesional subregions

To correlate T**_1_** value of each voxel at the terminal time point for each animal to its normalized PD intensity and quantified across selected ROI, the matching slice from the MRI atlas was registered to the PDw image at the terminal time point for each animal with the bUnwarpJ plugin with “Accurate” and “Save Transformations” mode checked. Whenever possible, the “Coarse” (Initial Deformation) to “Coase” (Final Deformation) setting was applied to register images. The registration results are visually inspected, in cases where optimal registration results cannot be achieved, the final and or initial deformation will be adjusted to “Very Coarse” and or “Fine.” The transformation matrix acquired was then applied to the labels (cortical GM, subcortical GM, and WM tracts) with “bunwarpj.bUnwarpJ_.loadElasticTransform” function to index images for each animal. The XY coordinates from PDw images, T**_1_** values from the T**_1_** mapping, and registered labels were extracted and quantified as stated above. The registered labels (numeric format) were rounded to the nearest integer, and voxels with matching index with the original label collection were kept for further analysis.

To contextualize the analyzed ratio of PDw image intensity to T**_1_** value, voxels were grouped into three colors in scatterplots and mapped onto the corresponding baseline image with ggplot2 (v3.4.2) and jpeg (v0.1-10) packages in R (**Fig5A**). The lesion mask was created the same way as stated above, and the normal-appearing WM (NA.WM) mask was created by subtracting lesion mask from WM mask registered to the terminal image. Voxels within the lesion mask were further divided into subregions, voxels with > 1250 ms T**_1_** values were colored in red (supplement set of voxels were colored in blue), mapped onto corresponding PDw images, and compared with ST results (**Fig5B**). The perilesional microenvironments were further analyzed by assigning 5 concentric rims outward from the lesion core (voxels with > 1250 ms T**_1_** values within the lesion mask) within the WM mask.

### Immunohistochemistry

Sections used for histology were formalin-fixed, paraffin-embedded (FFPE) sections cut at 5 µm from brain using a Leica RM2235 Manual Rotary Microtome. Superfrost^+^/Colorfrost^+^ microslides (75 x 25 mm, #EF15978Z, Daigger) were used to mounted sections and stored at room temperature. Before staining, sectioned slides were deparaffinized with xylene 3 times for 5 min each, rehydrated with EtOH (100%, 70%, 50% for 5 min each), and rinsed in DI H**_2_**O for 5 min at RT. Deparaffinized and rehydrated slides were submerged in 1X antigen retrieval solution (100X Tris-EDTA Buffer, pH 9.0, ab93684, Abcam) and placed in a tissue steamer (IHC-Tek Epitope Retrieval Steamer Set, NC0070392, IHC world) for 20 min to perform heat-induced epitope retrieval (HIER). At the end of HIER, sections were let cool for 10 min inside the steamer and transferred to pre-cooled (4°C) 1X TBS for 5 min, submerged in 3% H**_2_**O**_2_** for 10 min to block for endogenous peroxidase, and rinsed in 1X TBST (0.05% tween-20 in 1X TBS) for 1 min at RT. A parafilm pan (Super HT PAP Pens, 22006, Biotium) was used to create a solution barrier by demarcating each section after removing excessive liquid around it with Kimwipes, and 200 μL of blocking solution containing 50% serum-free protein block (X090930-2, Dako) and 50% normal horse serum (2.5% blocking solution, S-2012-50, Vector) was applied per section for 30 min at RT. Primary antibodies diluted in antibody diluent (S080983-2, Dako) were applied on sections for overnight at 4°C. Sections were rinsed in 1X TBST once for 1 min, then twice for 5 min each, and appropriate secondary antibodies were applied for 30 min at RT. Sections were rinsed in 1X TBST once for 1 min, then twice for 5 min each, and 200 μL of immunoperoxidase development solution (DAB Substrate Kit, ab64238, Abcam; Vector® VIP Substrate Kit, SK-4600, Vector) was applied per section for 1-10 min at RT. Chromogenic reactions were stopped by switching to DI water, and sections were rinsed with tap water for 5 min at RT. For double staining, 200 μL of alkaline phosphatase substrate solution (Vector® Blue Substrate Kit, SK-5300, Vector) was applied to each section for 10-30 min at RT. Chromogenic reactions were stopped by switching to DI water, and sections were rinsed with tap water for 5 min at RT. The following antibodies were used: mouse anti-IGFBP3 (R&D systems, MAB305-100, 1:200), rabbit anti-PAI1/SERPINE1 (Thermo Fisher, 13801-1-AP, 1:200), PV Poly-HRP Anti-Rabbit IgG (Leica, PV6119), PV Poly-HRP Anti-Mouse IgG (Leica, PV6119, 1:1), ImmPRESS®-AP Horse Anti-Rabbit IgG Polymer (Vector, MP-5401-50, 1:1), ImmPRESS®-AP Horse Anti-Mouse IgG Polymer (Vector, MP-5402-50, 1:1).

### Data availability

Raw and processed datasets are submitted to Gene Expression Omnibus (GEO). Source data are provided with this paper.

## Supporting information

SourceData & SupTable

## Acknowledgments

We thank Dr. Heather L. Narver, Dr. Stacey Piotrowski, and the staff of the NINDS animal program for the help and advice in animal care. We thank Dr. Qing Wang and the Dr. Miriam and Sheldon G. Adelson Medical Research Foundation (AMRF) Functional Genomics Resource at UCLA for valuable support in sequencing. We thank Dr. Chang-Ting Lin for his insightful guidance in quantitative data analysis. We thank the AMRF’s Program in Neurodegenerative Diseases–Multiple Sclerosis (APND-MD, DS, DSR) and the Intramural Research Program of NINDS (ZIA NS003119, DSR) for providing funding. We thank the NIH HPC Biowulf cluster for the computational resources utilized in this work. We sincerely appreciate their contributions, which were essential to the success of this research.

## Author contributions

J.-P.L. and D.S.R. designed the study, interpreted the results, and prepared manuscripts. J.-P.L., A.B., M.D., P.S., D.S.R., and S.J. developed protocols. J.-P.L., A.B., M.D., A.L., and P.S. handled animals and acquired MRI data. J.-P.L. and M.D. processed and analyzed MRI data. J.-P.L. acquired, processed, and analyzed snRNA-seq data. J.-P.L. and A.B. acquired, processed, and analyzed spatial transcriptomic and histology data. J.-P.L. cleaned and processed published datasets. D.H.G. and R.K. analyzed the data. D.P.S. provided critical intellectual content. D.S.R. supervised the study.

## Corresponding author

Correspondence to Jing-Ping Lin and Daniel S. Reich.

## Declaration of interests

D.S.R. has received research funding from Abata and Sanofi, unrelated to the current study.

---

**FigS1. related to Fig1.**
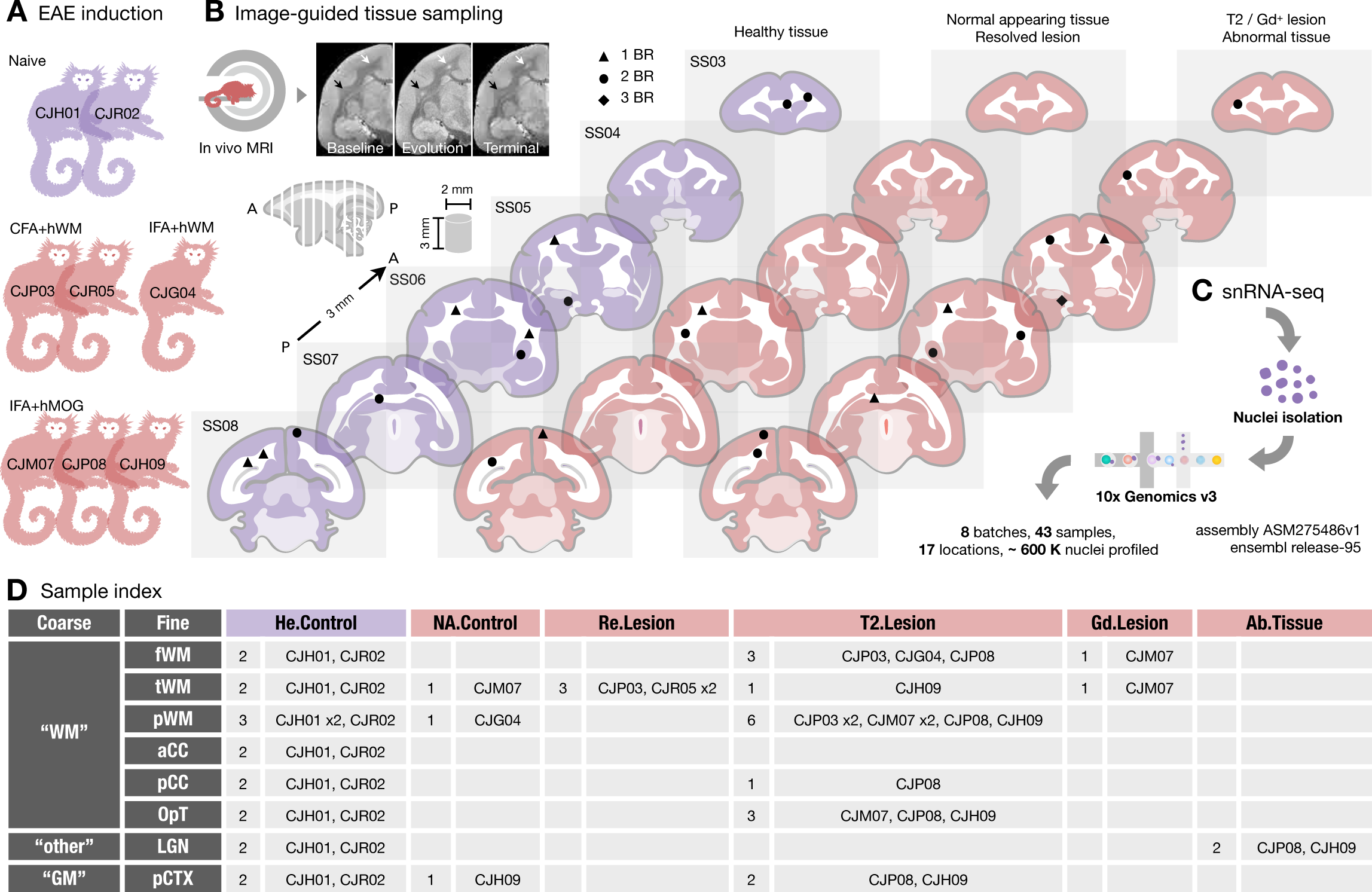
Experimental design used to create a transcriptome map for the evolution of white matter (WM) lesions with single nucleus resolution. (A) The dataset includes marmosets that were inoculated with human white matter (hWM) or recombinant human myelin oligodendrocyte glycoprotein (hMOG) emulsified in complete (CFA) or incomplete Freund’s adjuvant (IFA) to induce experimental autoimmune encephalomyelitis (EAE) and healthy, naïve controls. (B) The experimental workflow involved scanning and categorizing brain tissue using MRI. Postmortem brains were sliced into 3-mm slabs from anterior (A) to posterior (P). Specific areas of interest were sampled as cylinders with a diameter of 2 mm and height of 3 mm. These sampled areas were labeled on the standard slab (SS) index and grouped based on disease condition. The number of biological repeats (BR) is indicated by black annotations on the SS. (C) Nuclei were isolated from the sampled areas to prepare cDNA libraries, which were then sequenced. (D) Sampled areas were categorized into 3 types: coarse brain region, fine tissue location, and disease condition. Abbreviations: f (frontal), t (temporal), p (parietal), WM (white matter), a (anterior), p (posterior), CC (corpus callosum), OpT (optic tract), CTX (cortex), LGN (lateral geniculate nucleus), EAE (experimental autoimmune encephalomyelitis), He (healthy), NA (normal-appearing), Re (resolved), T2 (T**_2_**-hyperintense MRI detected), Gd (gadolinium), and Ab (abnormal).

**FigS2. related to Fig1.**
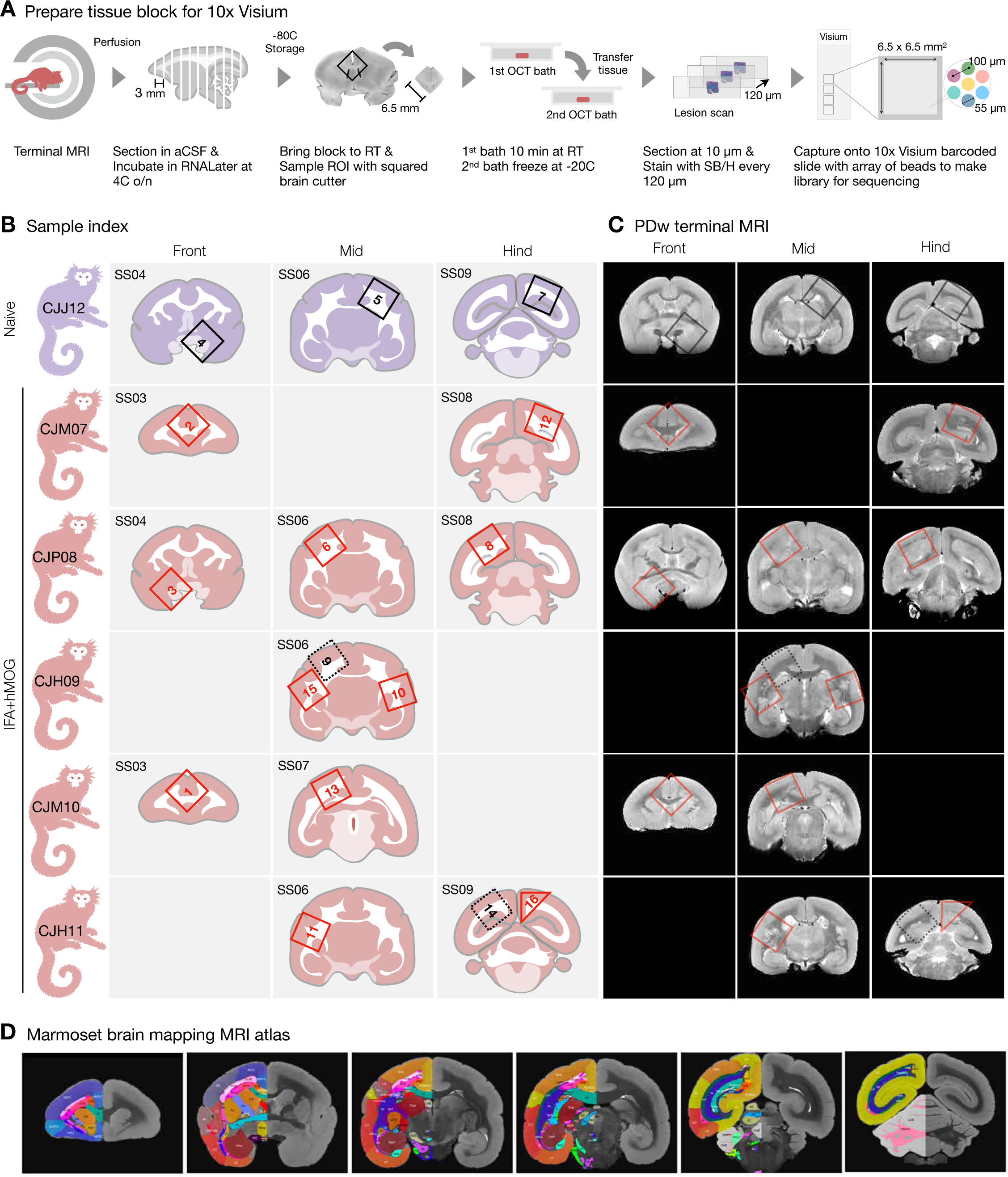
Experimental design for creating a transcriptome map of white matter (WM) lesions with spatial resolution. (A) The experimental workflow involves identifying WM lesions using magnetic resonance imaging (MRI) and preparing postmortem tissue for spatial transcriptome analysis using the 10x Visium platform. Abbreviations: aCSF (artificial cerebrospinal fluid), RT (room temperature), ROI (region of interest), OCT (optimal cutting temperature), SB (Sudan black), H (hematoxylin). (B) The dataset includes marmosets inoculated with recombinant human myelin oligodendrocyte glycoprotein (hMOG) emulsified in incomplete Freund’s adjuvant (IFA). Sampled areas were blocked (6.5x6.5x3 mm³), labeled onto the standard slab (SS) index, and grouped by matching brain area across diseased conditions. (C) Proton density-weighted (PDw) MRI was performed at the terminal time point and matched with the sampled tissue areas. (D) A postmortem MRI atlas overlaid with brain region labels was used to match with the selected terminal PDw MRI.

**FigS3. related to Fig1.**
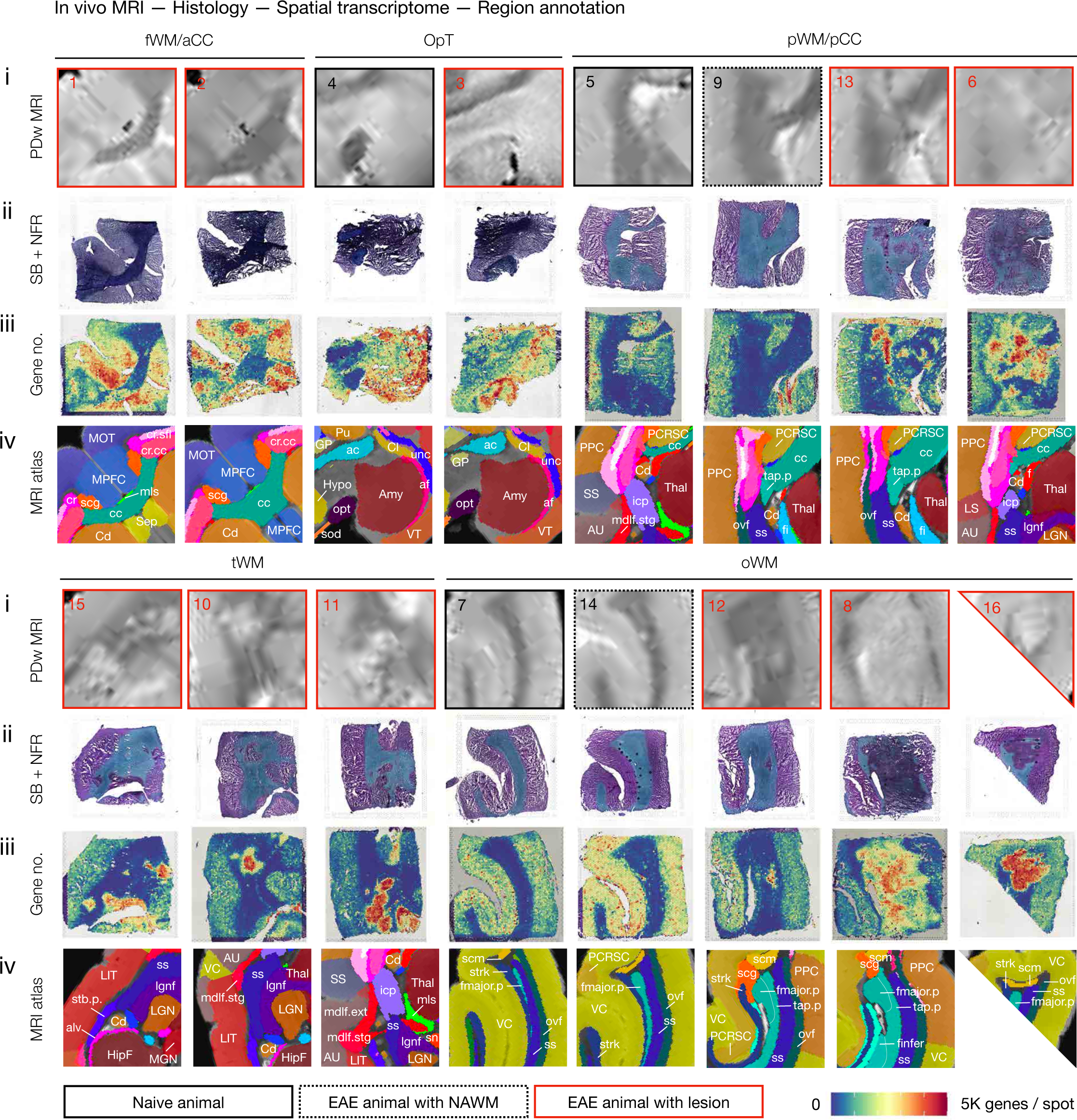
Matched brain regions imaged by different modalities. i) In vivo brain PDw MRI acquired on a 7 Tesla scanner at the disease terminal. ii) Histological examination of myelin content using Sudan black (SB) and nuclear fast red (NFR) staining of postmortem tissue. iii) RNA landscape visualized through the 10x Visium platform. iv) Marmoset MRI atlas with region annotations. See source data for the full list of abbreviations for brain regions.

**FigS4. related to Fig1.**
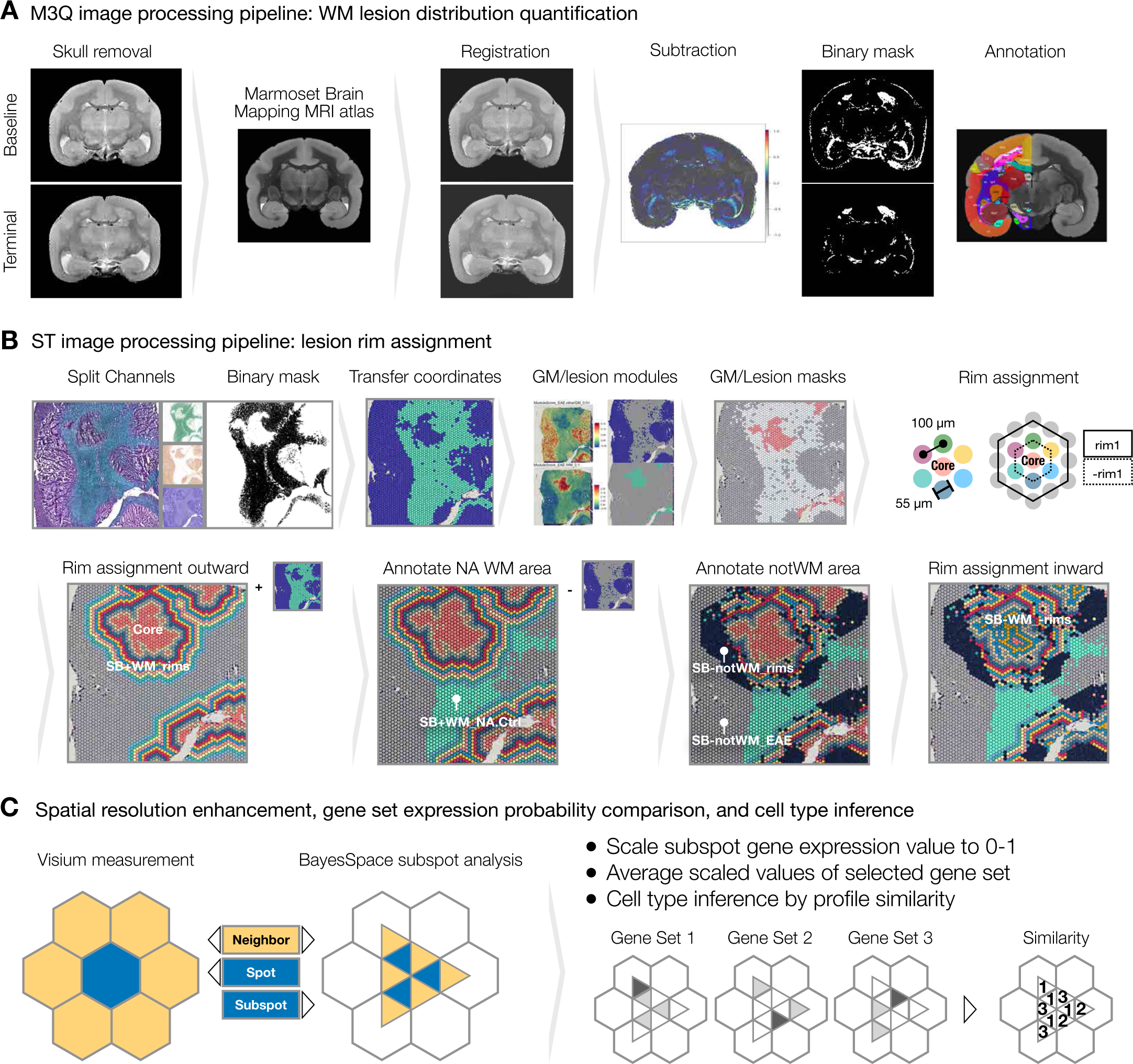
Image processing and resolution enhancement pipelines used to quantify white matter (WM) lesion load, generate lesion subregion masks, and deconvolute mixed transcript signals. (A) To gain insight into lesion development and progression, an <u>M</u>RI characterization of <u>M</u>S-like lesions in the <u>M</u>armoset <u>Q</u>uantitatively (M3Q) pipeline was developed, including the following steps (**Methods**): Proton density-weighted (PDw) MRI images at baseline (before EAE induction) and terminal (before tissue collection) time points were subjected to the N4 bias field correction algorithm. The brain portion of the images, corresponding to the region of interest, was extracted using a skull-removal algorithm to improve image alignment. Each image was individually registered to the marmoset MRI atlas using bUnwarpJ, achieving spatial alignment across time points and animals. MRI intensity changes were calculated by subtracting the normalized terminal image from the baseline image, providing information about lesion location. Binary lesion masks were created by applying intensity thresholding to the subtracted images, segmenting the region of interest. The WM portion of the lesion mask was parsed and analyzed using atlas annotation indexing, enabling further characterization of the lesions based on location and distribution. (B) To gain insight into the regionally enriched signal distribution within and near the WM lesion, a spatial transcriptome (ST) image processing pipeline was employed with the following steps: The myelinated WM area (Sudan black-positive) was extracted using the "Color Deconvolution" function in Fiji with the default "H DAB" setting, resulting in an SB^+^ WM binary mask. The coordinates of the SB^+^ WM mask were transferred to the 10x Visium spot hexagon coordinate system using Seurat, facilitating the spatial mapping of gene expression data. To distinguish SB^-^ gray matter (GM) from SB^-^ demyelinated WM, GM and lesion gene module scores were calculated and filtered to create GM and lesion masks accordingly. Spots that exhibited both SB^-^ and IMM^+^ signals were identified as the lesion core. Next, 10 concentric rims (SB^+^WM_rims) extending outward from the lesion core were assigned to mark the adjacent lesion neighborhoods. The normal-appearing (NA) WM area was annotated by subtracting the lesion neighborhoods from the SB^+^ WM mask in animals with experimental autoimmune encephalomyelitis (EAE), and this region was labeled as “SB^+^WM_NA.Ctrl." Additionally, lesion neighborhoods that overlapped with the GM mask were labeled as "SB^-^notWM_rims," while the supplemental area was labeled as "SB^-^notWM_EAE" in animals with EAE. Subregions within the lesion core were further divided based on centripetal rim assignments (SB^-^WM_- rims). For healthy animals, "SB^+^WM_He.Ctrl" and “SB^-^notWM_He" labels were used to annotate tissue with or without SB staining, respectively. (C) The spatial transcriptome resolution, initially measured at the spot level using the 10x Visium platform, was further enhanced to the subspot level using the BayesSpace algorithm. The enhanced signals at the subspot level were then rescaled to a ratio of 1 across genes. To gain insights into cell-type distributions, the averaged expression of gene sets enriched in specific cell types, acquired from single-nucleus RNA sequencing (snRNA-seq) references, was calculated. Cell-type locations were inferred by assessing the relative profile similarity score, allowing identification and mapping of different cell types within the spatial transcriptome data. By employing these techniques, the spatial transcriptome analysis achieved higher resolution, enabling a more detailed understanding of the cellular composition and gene expression patterns within the tissue of interest.

**FigS5. related to Fig2.**
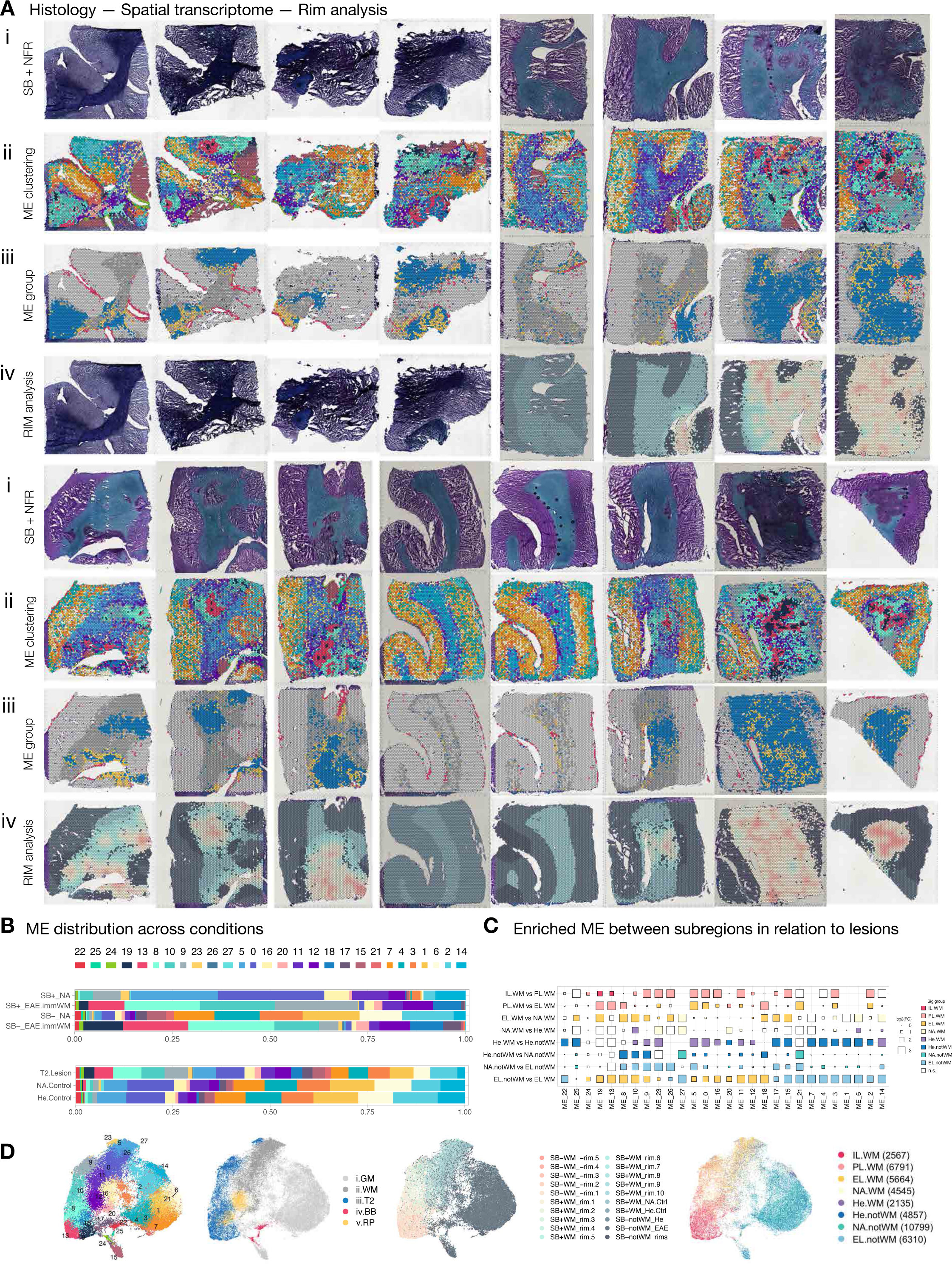
Matched brain regions of interest across pathological states are color-coded based on different microenvironment (ME) phenotypes. (A) To demonstrate the phenotypic characterization of the ME within matched brain regions, contrasts between: i) myelin content visualized through Sudan black (SB) and nuclear fast red (NFR) staining, ii) unbiased ME clustering achieved through transcriptome similarity analysis, iii) 5 ME groups assigned by ME profile similarity, and iv) lesion subregions assigned using rim analysis and overlaid onto SB/NFR stained images, were indexed. The "Color Deconvolution" (**FigS4B**) for Samples 1–4 was unsuccessful due to suboptimal contrast between SB and NFR staining, resulting in their exclusion from the lesion subregion assignment in the rim analysis; however, they are included for ME clustering analysis. (B) Stacked bar plots show the relative proportions of transcriptomic ME at the spot resolution across different pathological states (top), as well as the levels of myelin and inflammation (bottom). (C) Dot plots depict the change in spot proportion across subregional white matter (WM) of Samples 5–16. Significantly (FDR < 0.05 & abs(Log2FC) > 0.5) enriched ME between pairs of subregional WM area are colored accordingly. “IL.WM” contains SB-WM_-rim5 to SB-WM_-rim2, “PL.WM” contains SB-WM_-rim1 to SB+WM_rim1, “EL.WM” contains SB+WM_rim2 to SB+WM_rim10), “NA.WM” contains SB+WM_NA.Ctrl, “He.WM” contains SB+WM_He.Ctrl, “He.notWM” contains SB-notWM_He, “NA.notWM” contains SB-notWM_EAE, and “EL.noWM” contains SB-notWM_rims. (D) UMAP plots colored by ME cluster (**left**), subregions assigned by rim analysis (**middle**), and subregional WM (**right**).

**FigS6. related to Fig2.**
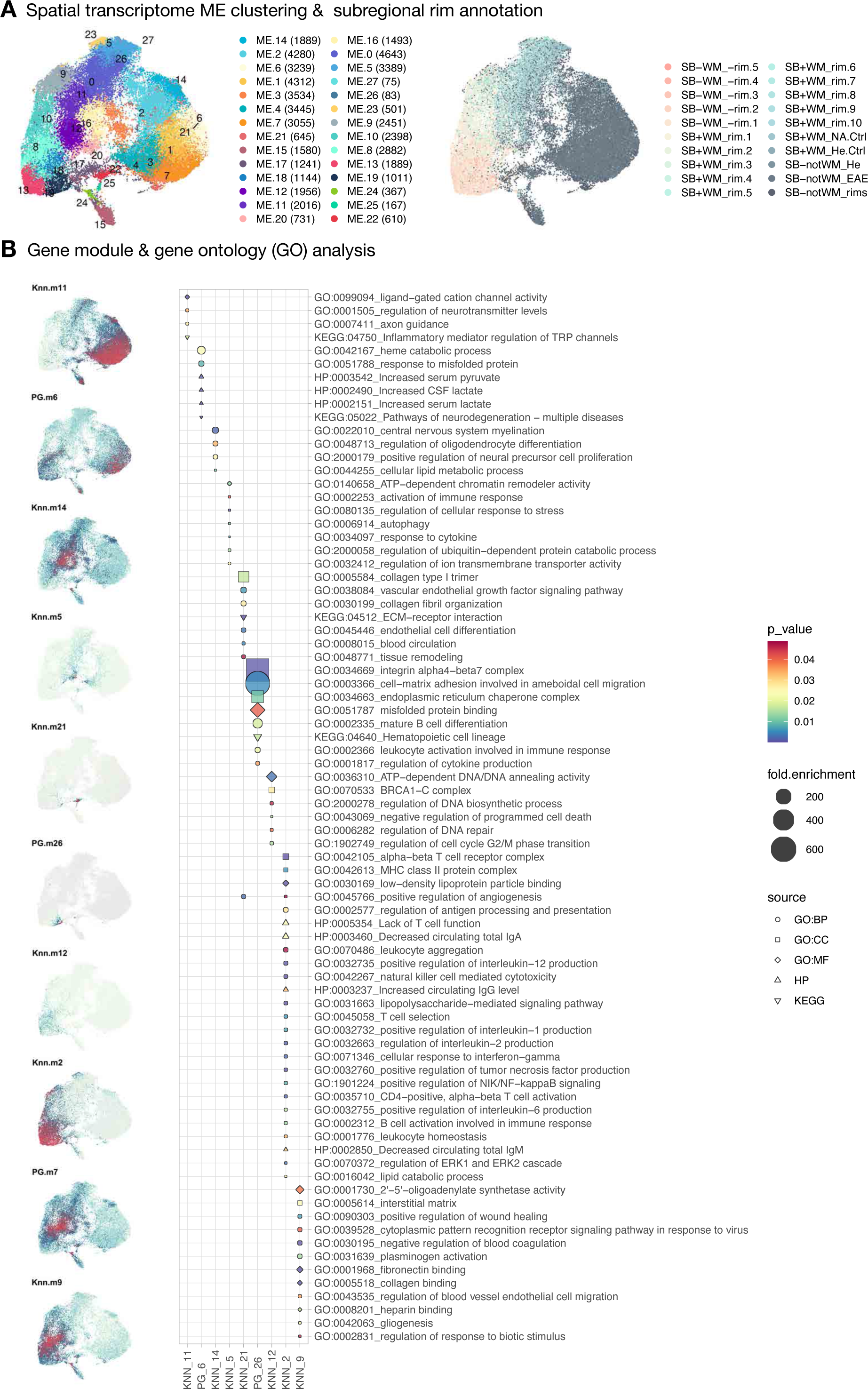
Spatial organization of transcriptomes within different microenvironments and functional enrichment of gene modules within subregions. (A) UMAP scatter plots color-coded by microenvironment (ME) clustering and subregional labeling. The number of spots per microenvironment is indicated in parentheses. Abbreviations: SB (Sudan black), WM (white matter), NA (normal appearing), He (healthy), Ctrl (control). (B) UMAP scatter plots color-coded by gene expression. Gene modules are annotated with enriched Gene Ontology (GO) terms, including three major subontologies: Molecular Functions (MF), Biological Process (BP), and Cellular Component (CC). When available, additional annotations from the KEGG and HP databases are included. Specifically, the Knn.m11 gene module is enriched in ME14, 2, 6, 1, 3, 4, 7, 21, 15, 17, which dominate the SB-notWM (gray matter) area and are involved in the regulation of neurotransmitter levels, as expected. Perilesional (SB+WM_rims) and lesional (SB-WM_rims) ME are enriched with modules involved in various processes, including myelination and lipid metabolism (Knn.m14), glycometabolism and neurodegeneration (PG.m6), immune and stress response (Knn.m5 and Knn.m2), extracellular matrix (ECM) and vascular function (Knn.m21 and Knn.m9), hematopoietic and leukocytic cell development (PG.m26 and Ken.m2), and programmed cell death and cell cycle (Knn.m12).

**FigS7. related to Fig2.**
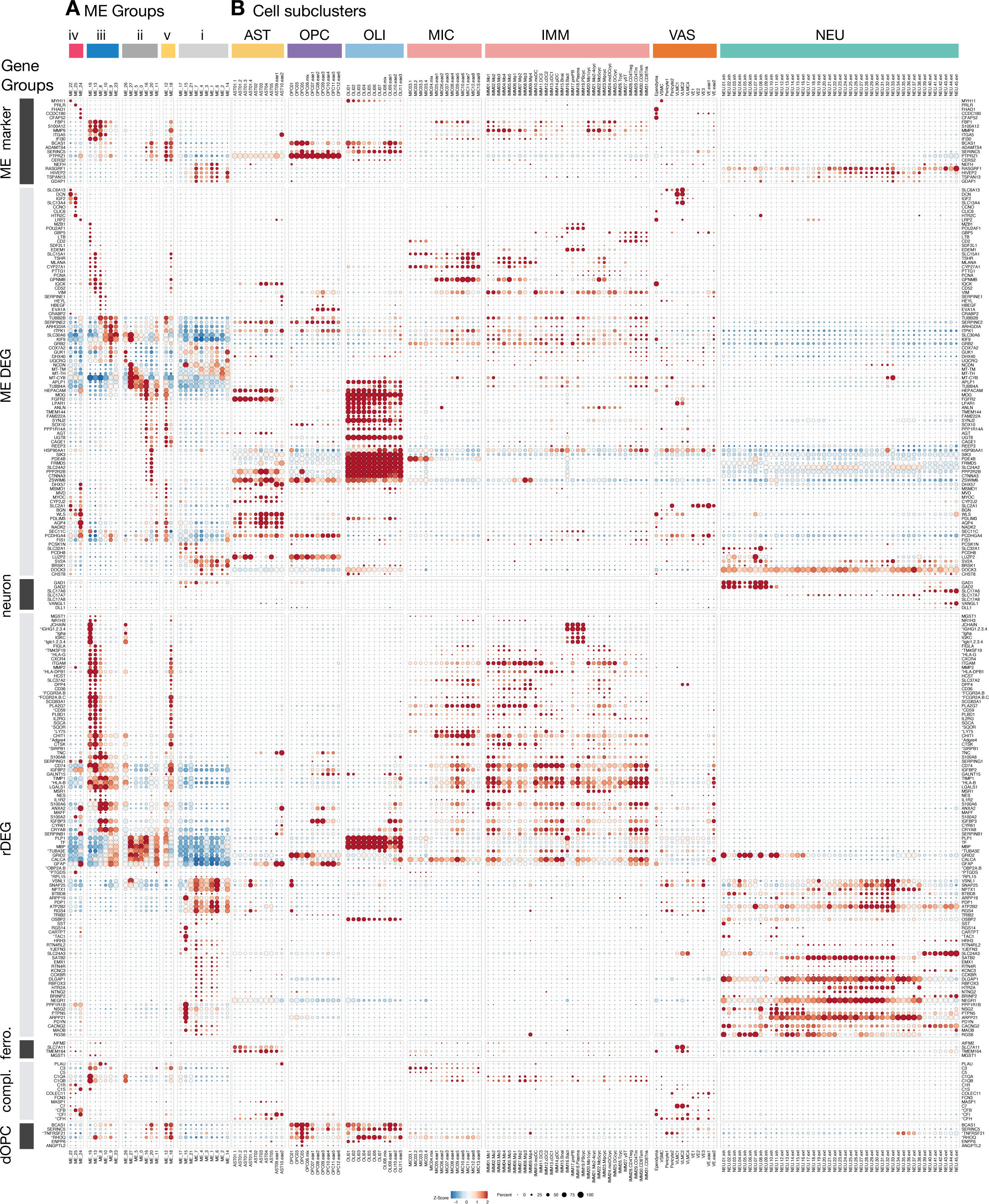
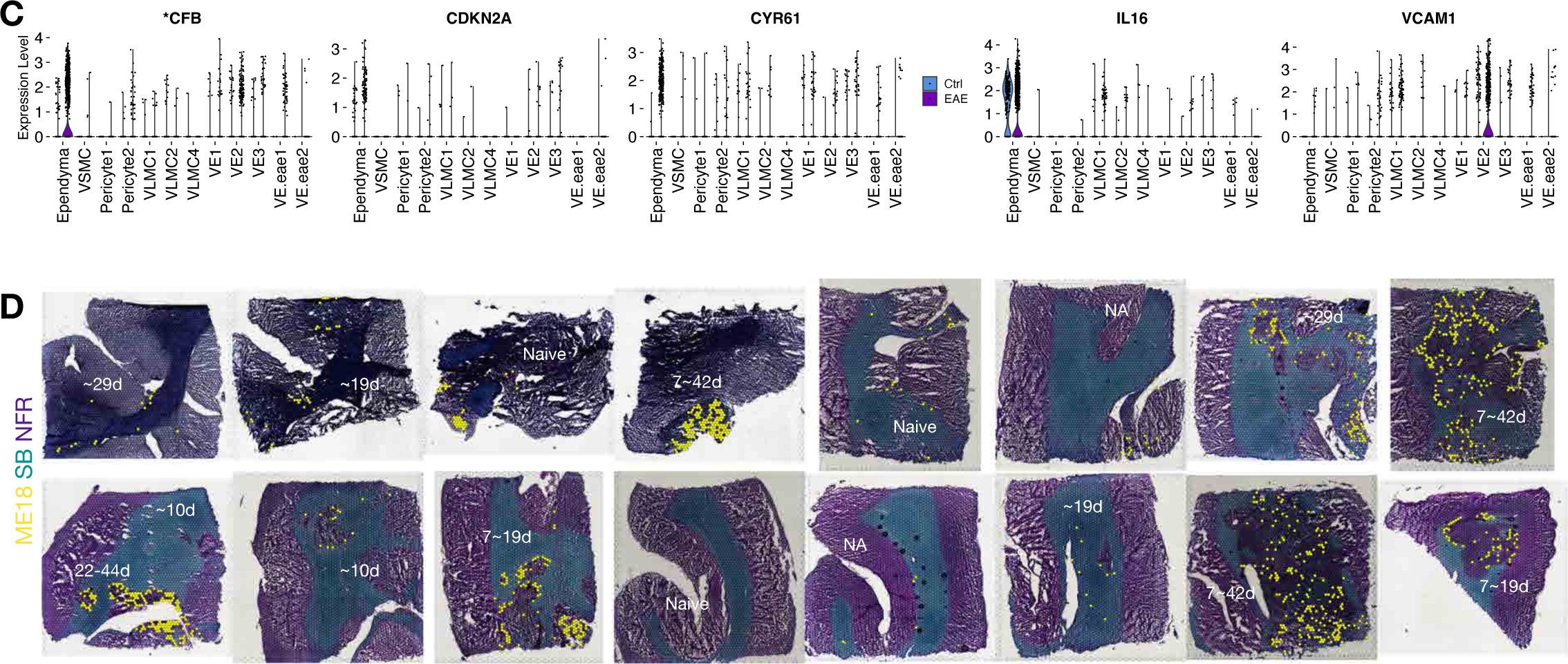
Complement factor B (CFB) expression is elevated in EAE ependyma compared to control, and ME18 is enriched in older lesions. (A) Dot plot showing the averaged and scaled expression of selected genes across microenvironments (ME). Gene names starting with “*” indicate human (hs) or mouse (mm) orthologs of marmoset gene identification numbers (See **Table S9** for the full list). (B) Dot plot showing the averaged and scaled expression of selected genes across L2 subclusters. Genes are split into groups to aid label tracking. Abbreviations: ME marker (genes used to annotate ME groups in **Fig2A**), ME DEG (differentially expressed genes across 28 ME), rDEG (regional differentially expressed genes), ferro. (ferroptosis genes), compl. (complement genes), dOPC (differentiating OPC enriched genes). (C) Violin plot showing the expression of \**CFB* (human homolog of marmoset ENSCJAG00000048204), *CDKN2A*, *CYR61*, *IL16*, and *VCAM1* across vascular cells in control and EAE. (D) SB/NFR-stained tissue across 16 ROI labeled by the distribution of ME18 (yellow dots) and annotated by lesion age, dated by longitudinal MRI (**Methods**).

**FigS8. related to Fig2.**
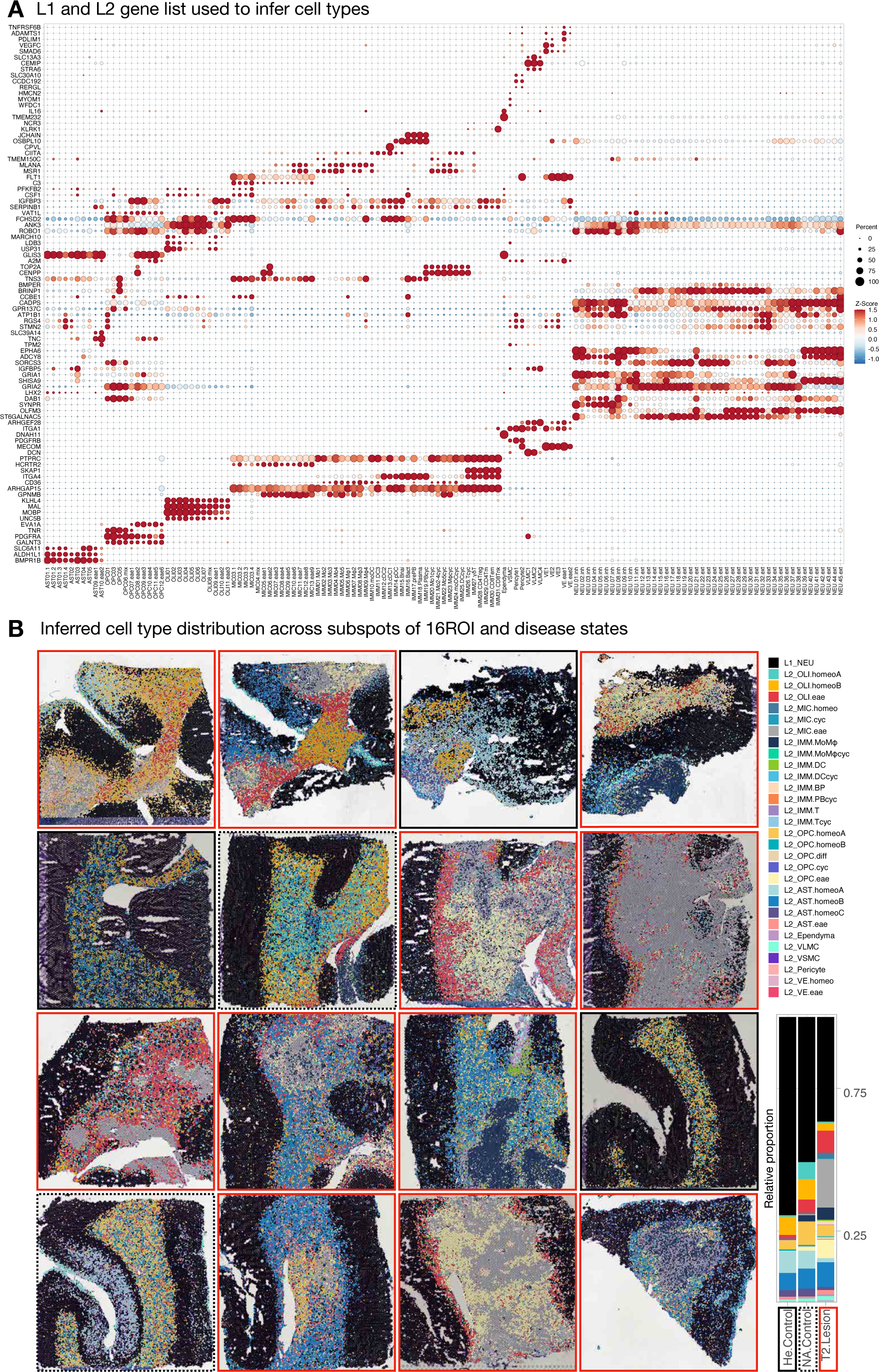
Selected L1 and L2 genes for each cell class and subcluster. (A) Dot plot showing the averaged and scaled expression of selected genes used to infer cell types for BayesSpace-enhanced subspots across L2 subclusters. (B) BayesSpace-enhanced subspots colored by inferred cell types across 16 ROI labeled by crude disease category.

**FigS9. related to Fig3.**
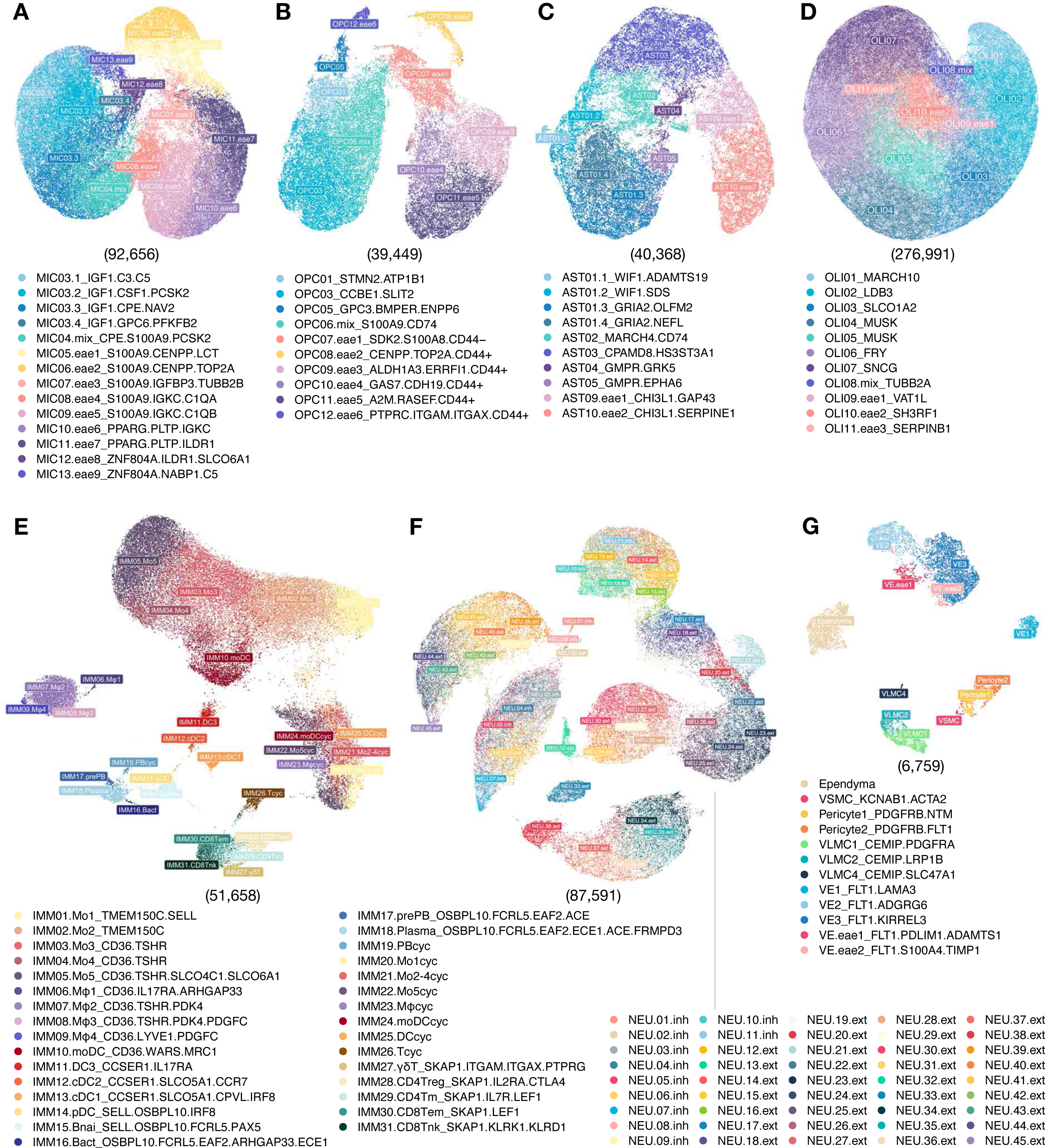
UMAP scatter plot of the level 2 analysis conducted on major cell classes. (A) – (G) The level 1 (L1) analysis identified 6 major cell classes, which were further divided into 7 partitions in the level 2 (L2) analysis. UMAP scatter plots show subclustering of the following cell classes: microglia (MIC), oligodendrocyte progenitor cells (OPC), astrocytes (AST), oligodendrocytes (OLI), peripherally derived immune cells (P.IMM), neurons (NEU), and vascular/meningeal/ventricular cells (VAS). The number of nuclei analyzed in each L2 UMAP plot is listed in parentheses.

**FigS10. related to Fig3.**
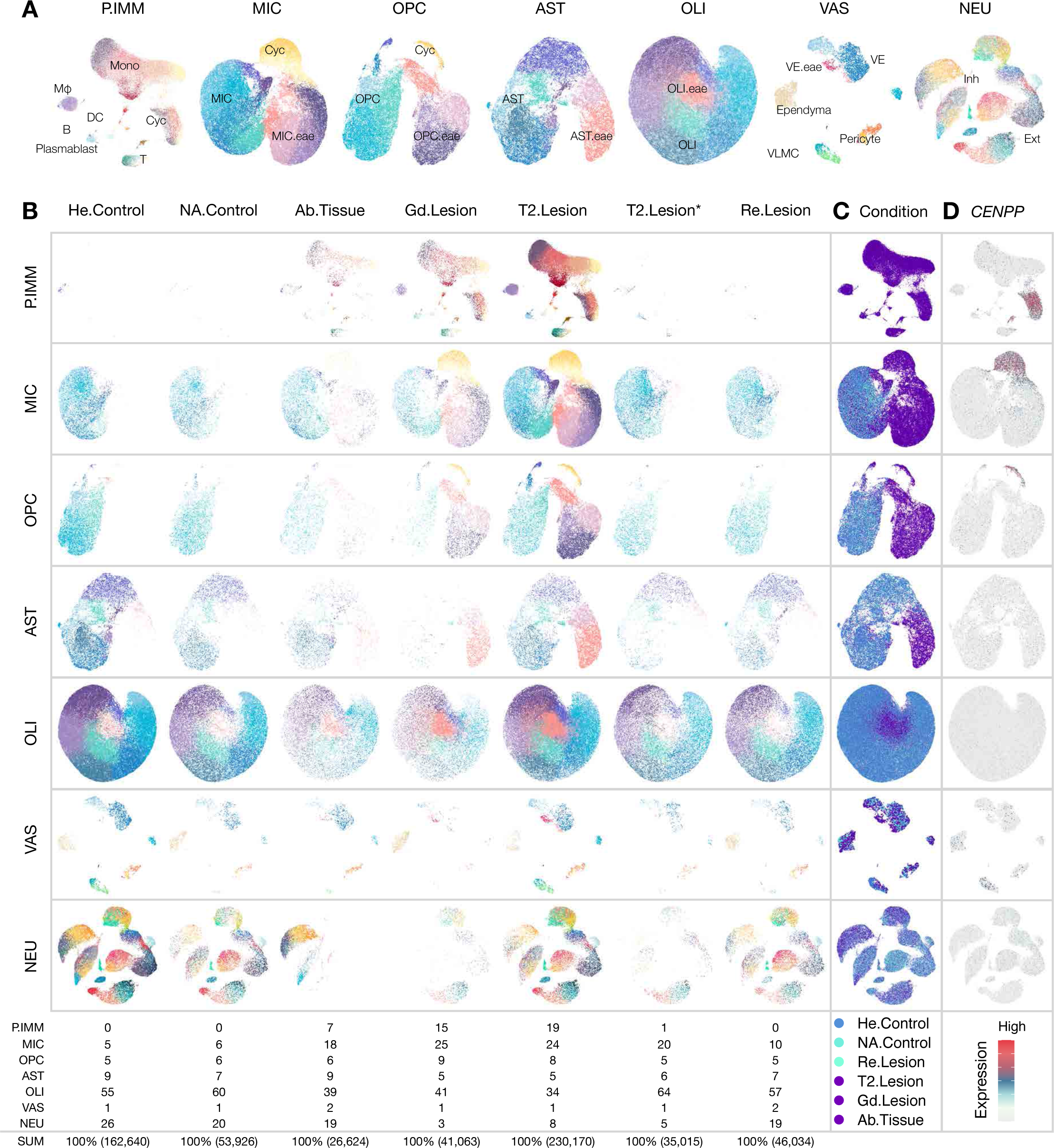
Glial and immune cells diversity across pathological states. (A) UMAP plots of the level 2 (L2) partitions labeled with major subtypes enriched in animals with experimental autoimmune encephalomyelitis (EAE). The abbreviations: mono (monocytes), Mφ (macrophages), DC (dendritic cells), B (B cells), T (T cells), Cyc (cycling cells), VE (vascular endothelial cells), VLMC (vascular leptomeningeal cells), Inh (inhibitory neurons), and Ext (excitatory neurons). (B) UMAP plots split by disease conditions and colored by level 2 subclusters. This visual representation allows for the comparison and observation of cell distribution patterns specific to each disease condition. The abbreviations: He (healthy), NA (normal-appearing), Re (resolved), T2 (transverse relaxation time), Gd (gadolinium), and Ab (abnormal). (C) UMAP plots across the L2 cell classes, colored by disease conditions. This representation helps in visualizing the dominant subclusters across different disease conditions. (D) UMAP plots across the L2 cell classes, colored by the expression level of the CENPP (Centromere Protein P) gene. This annotation allows for the identification of cycling cells within the cell classes.

**FigS11. related to Fig3.**
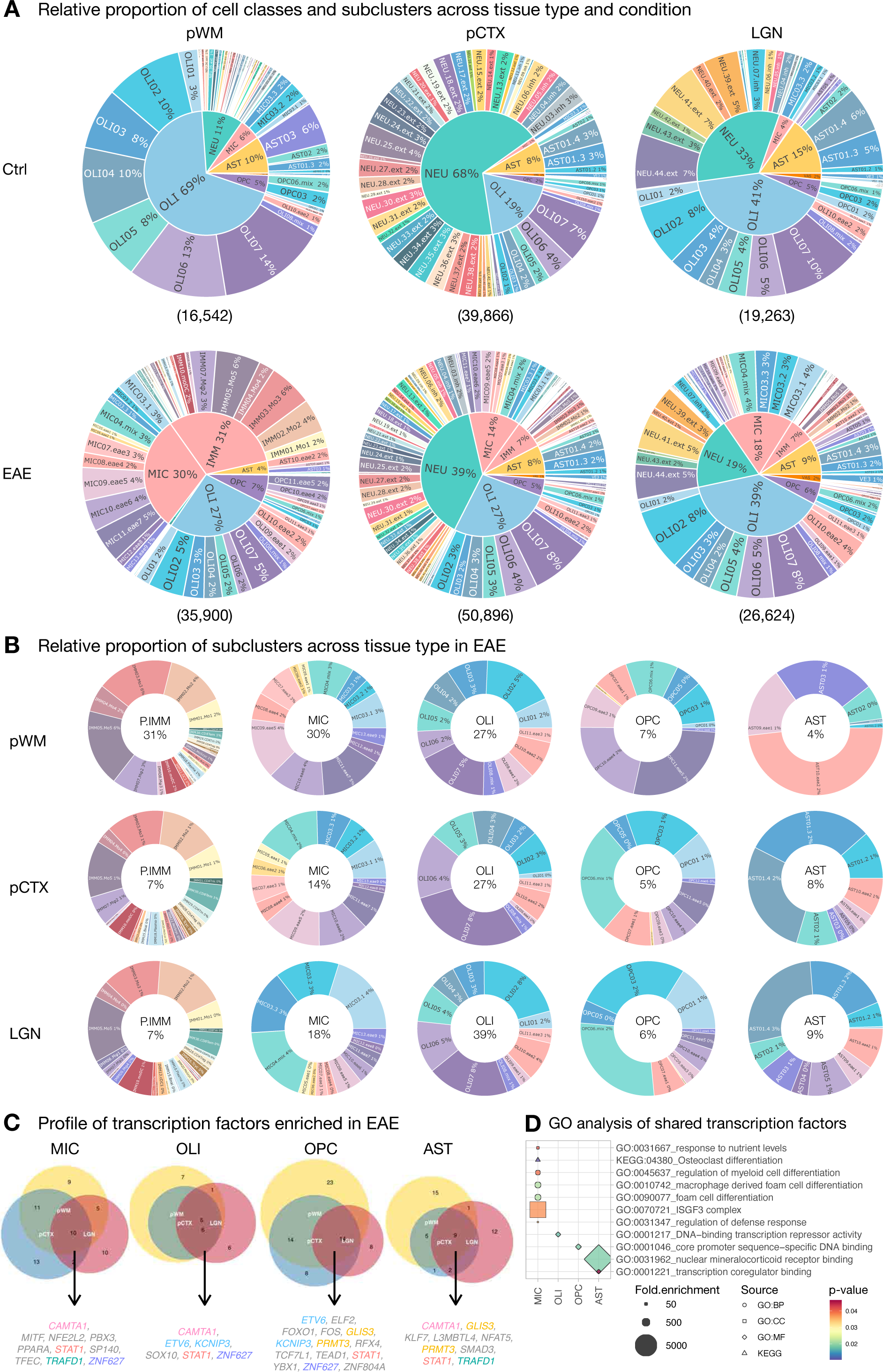
Comparative analysis of cell types and changes in transcription factors across different tissue types in response to EAE. (A) The inner pie charts show the relative nuclei proportion of L1 cell classes, and the outer donut charts display L2 sub-clusters. Both control and EAE samples for each tissue type are included, with the total number of nuclei listed in parentheses. In control animals, the composition of L1 cell classes varied across tissue types, as expected. Specifically, a higher number of glial cells were found in parietal white matter (pWM), while neurons were predominant in parietal cortex (pCTX) and lateral geniculate nucleus (LGN) region. In EAE animals, there was a significant expansion of microglia (MIC) and peripheral immune cells (P.IMM) partitions in all tissue types. (B) Donut charts show the relative nuclei proportion of glial and immune clusters in EAE animals across different tissue types. The compositions of the P.IMM and oligodendrocytes (OLI) partitions were largely similar across tissue types. However, the compositions of MIC, oligodendrocyte precursor cell (OPC), and astrocyte (AST) partitions were unique to certain tissue types. Specifically, the OPC and AST compositions were more similar in pCTX and LGN compared to pWM. On the other hand, the MIC composition was more similar in pWM and pCTX compared to LGN. (C) Venn diagrams illustrate the similarity and diversity of transcription factors that are significantly enriched in EAE compared to control animals across different tissue types for each glial cell class. The elevated transcription factors shared by all tissue types in EAE animals are listed for each cell class, and the shared transcription factors across different cell classes are color-coded accordingly. (D) Dot plot shows selective GO terms enriched for each list of shared transcription factors per glial cell class listed in (C).

**FigS12. related to Fig3.**
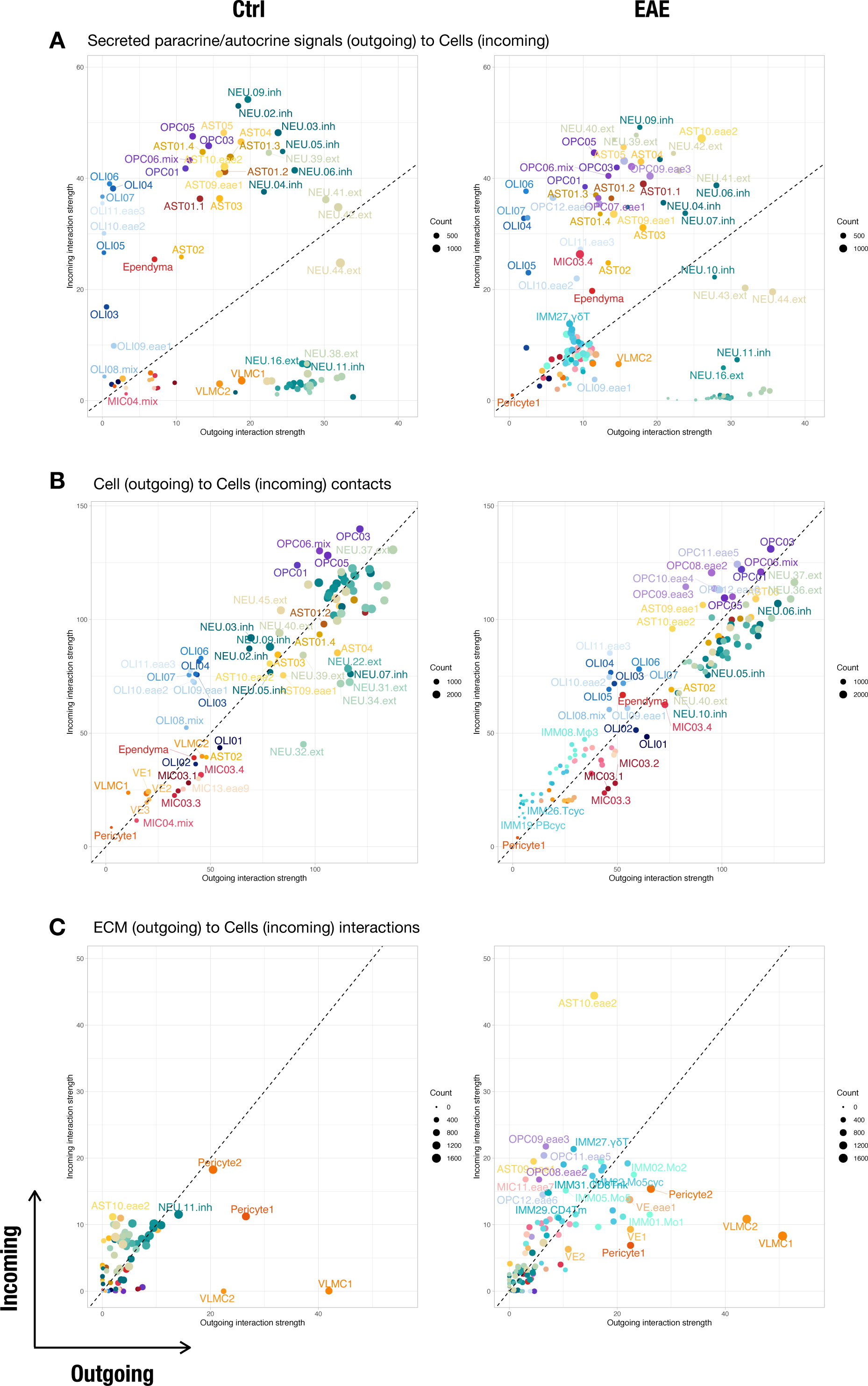
Communication preferences of cells within the white matter (WM) inferred by CellChat. (A) Incoming and outgoing secreted signal-to-cell strength across different conditions. (B) Incoming and outgoing cell-to-cell signaling strength across different conditions. (C) Incoming and outgoing extracellular matrix (ECM) signal-to-cell strength across different conditions.

**FigS13. related to Fig4.**
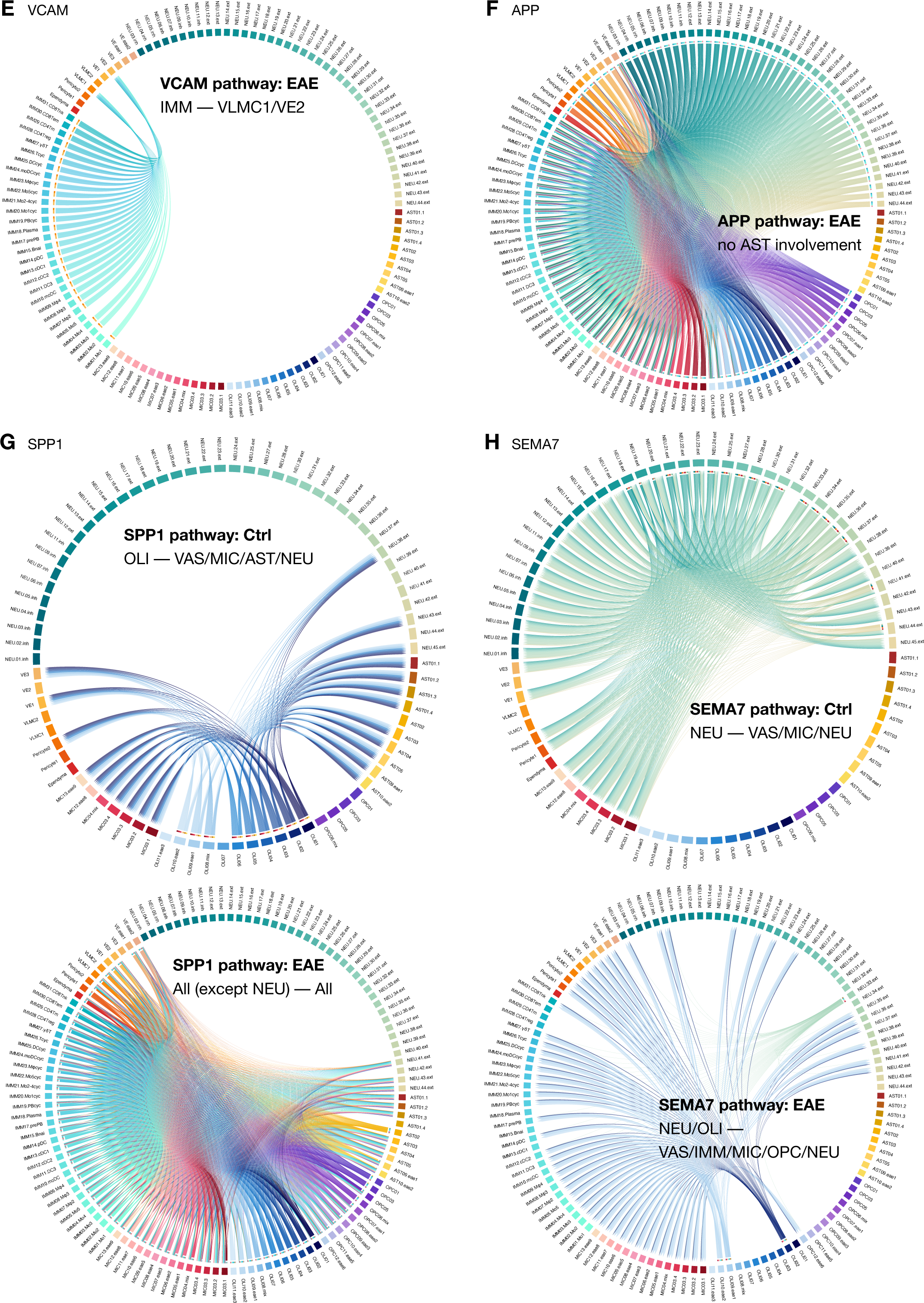
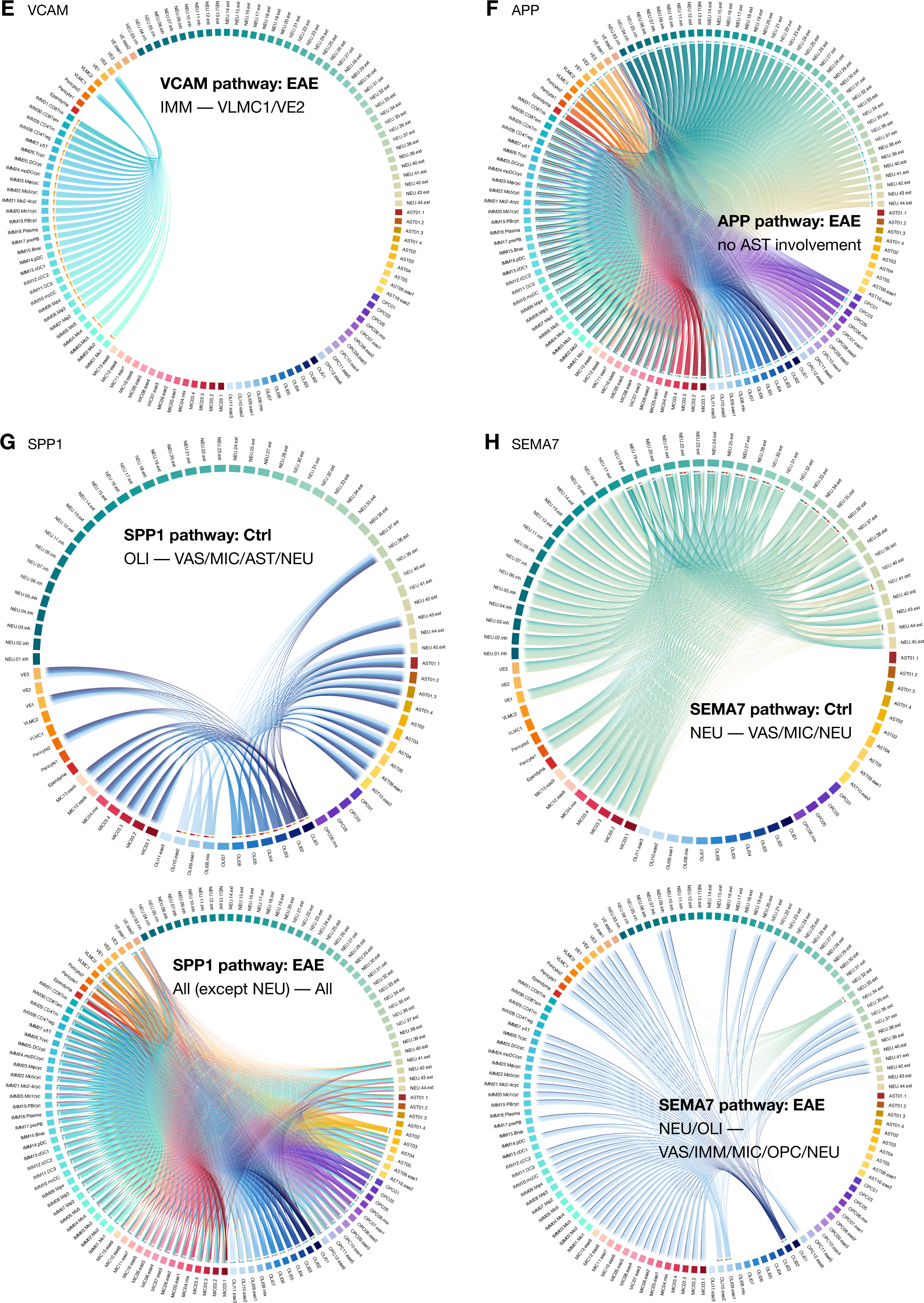
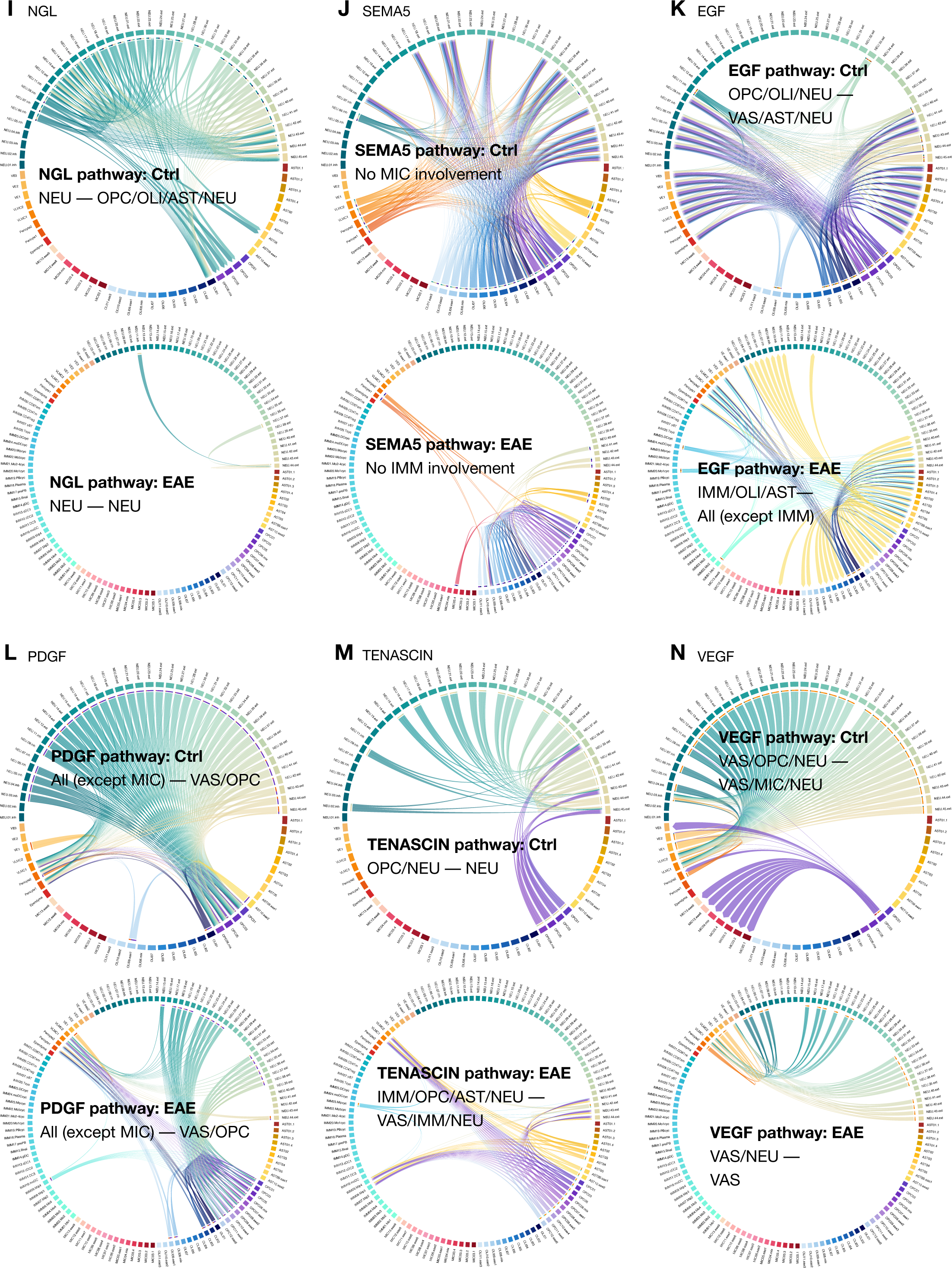
Chord diagrams and network plots summarize the intercellular communication profile of selected pathways. (A) Inferred sender (ligand) and receiver (receptor) pairs of the MHC-I pathway between L2 subclusters of white matter (WM) in EAE animals. (B) Same as (A) for MHC-II pathway in EAE animals. (C) Same as (A) for CD45 pathway in EAE animals. (D) Same as (A) for CD86 pathway in EAE animals. (E) Same as (A) for VCAM pathway in EAE animals. (F) Same as (A) for APP pathway in EAE animals. (G) Same as (A) for SPP1 pathway in control and EAE animals. (H) Same as (A) for SEMA7 pathway in control and EAE animals. (I) Same as (A) for NGL pathway in control and EAE animals. (J) Same as (A) for SEMA5 pathway in control and EAE animals. (K) Same as (A) for EGF pathway in control and EAE animals. (L) Same as (A) for PDGF pathway in control and EAE animals. (M) Same as (A) for TENASCIN pathway in control and EAE animals. (N) Same as (A) for VEGF pathway in control and EAE animals.

**FigS14. related to Fig4.**
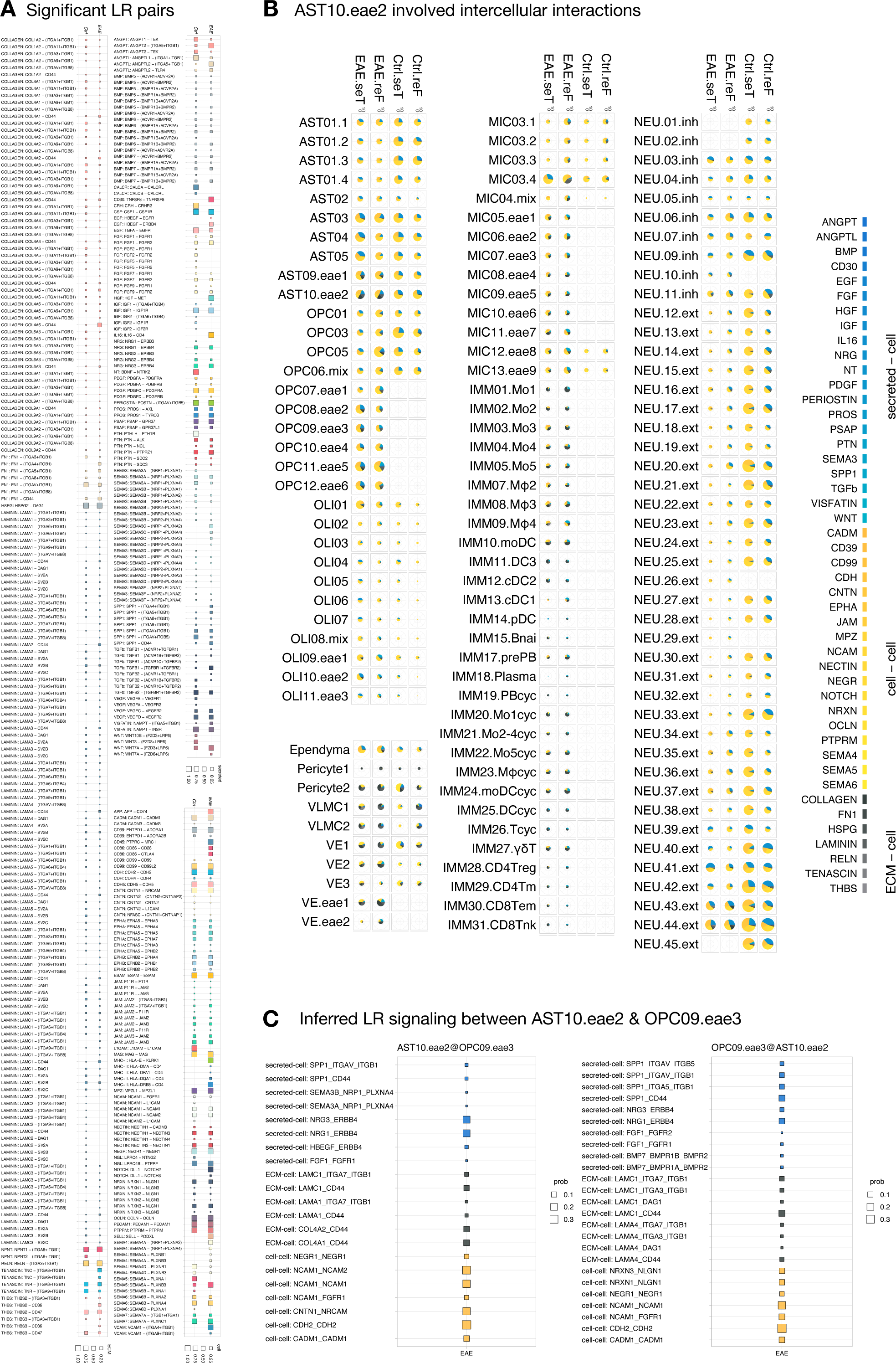
Significant ligand-receptor pairs inferred by CellChat across different pathological states. (A) Dot plots showcase the relative contribution of each ligand-receptor (LR) pair to each signaling pathway; signaling strength is indicated by dot color. Interactions are categorized into 3 types: secreted autocrine/paracrine signaling interactions (secreted-cell), cell-cell contact interactions (cell-cell), and extracellular matrix (ECM)-receptor interactions (ECM-cell). Each label follows the format of "pathway name: ligand name – receptor name.” (B) Pie charts depict the relative contribution of each pathway to overall interactions involving the AST10.eae2 subcluster per pathological state. (C) Dot plots show the relative contribution of each LR pair to each signaling category between the AST10.eae2 and OPC09.eae3 subclusters. Each label follows the format of "pathway category: ligand name_receptor name.”

## Notes

### Competing Interest Statement

The authors have declared no competing interest.

## References

1. Kuhlmann T, Moccia M, Coetzee T, Cohen JA, Correale J, Graves J, et al. Multiple sclerosis progression: time for a new mechanism-driven framework. Lancet Neurol. 2023 Jan;22(1):78–88.

2. McGinley MP, Goldschmidt CH, Rae-Grant AD. Diagnosis and Treatment of Multiple Sclerosis: A Review. JAMA. 2021 Feb 23;325(8):765–79.

3. Kametani Y, Shiina T, Suzuki R, Sasaki E, Habu S. Comparative immunity of antigen recognition, differentiation, and other functional molecules: similarities and differences among common marmosets, humans, and mice. Exp Anim. 2018 Jul 30;67(3):301–12.

4. ’t Hart BA. Experimental autoimmune encephalomyelitis in the common marmoset: a translationally relevant model for the cause and course of multiple sclerosis. Primate Biol. 2019 May 10;6(1):17–58.

5. Wattjes MP, Ciccarelli O, Reich DS, Banwell B, de Stefano N, Enzinger C, et al. 2021 MAGNIMS-CMSC-NAIMS consensus recommendations on the use of MRI in patients with multiple sclerosis. Lancet Neurol. 2021 Aug;20(8):653–70.

6. Filippi M, Preziosa P, Banwell BL, Barkhof F, Ciccarelli O, De Stefano N, et al. Assessment of lesions on magnetic resonance imaging in multiple sclerosis: practical guidelines. Brain. 2019 Jul;142(7):1858–75.

7. Fisher E, Reich DS. Imaging new lesions: enhancing our understanding of multiple sclerosis pathogenesis. Neurology. 2013 Jul 16;81(3):202–3.

8. Absinta M, Vuolo L, Rao A, Nair G, Sati P, Cortese ICM, et al. Gadolinium-based MRI characterization of leptomeningeal inflammation in multiple sclerosis. Neurology. 2015 Jul;85(1):18–28.

9. Al-Louzi O, Letchuman V, Manukyan S, Beck ES, Roy S, Ohayon J, et al. Central Vein Sign Profile of Newly Developing Lesions in Multiple Sclerosis: A 3-Year Longitudinal Study. Neurology - Neuroimmunology Neuroinflammation [Internet]. 2022 Mar 1 [cited 2023 Jul 29];9(2). Available from: https://nn.neurology.org/content/9/2/e1120

10. Maggi P, Absinta M, Grammatico M, Vuolo L, Emmi G, Carlucci G, et al. Central vein sign differentiates Multiple Sclerosis from central nervous system inflammatory vasculopathies. Annals of Neurology. 2018 Feb;83(2):283–94.

11. Absinta M, Sati P, Masuzzo F, Nair G, Sethi V, Kolb H, et al. Association of Chronic Active Multiple Sclerosis Lesions With Disability In Vivo. JAMA Neurol. 2019 Dec 1;76(12):1474–83.

12. Kolb H, Al-Louzi O, Beck ES, Sati P, Absinta M, Reich DS. From pathology to MRI and back: Clinically relevant biomarkers of multiple sclerosis lesions. Neuroimage Clin. 2022;36:103194.

13. Procaccini C, De Rosa V, Pucino V, Formisano L, Matarese G. Animal models of Multiple Sclerosis. Eur J Pharmacol. 2015 Jul 15;759:182–91.

14. Boretius S, Schmelting B, Watanabe T, Merkler D, Tammer R, Czéh B, et al. Monitoring of EAE onset and progression in the common marmoset monkey by sequential high-resolution 3D MRI. NMR in Biomedicine. 2006;19(1):41–9.

15. Gaitán MI, Maggi P, Wohler J, Leibovitch E, Sati P, Calandri IL, et al. Perivenular brain lesions in a primate multiple sclerosis model at 7-tesla magnetic resonance imaging. Mult Scler. 2014 Jan 1;20(1):64–71.

16. Maggi P, Macri SMC, Gaitán MI, Leibovitch E, Wholer JE, Knight HL, et al. The formation of inflammatory demyelinated lesions in cerebral white matter. Annals of Neurology. 2014;76(4):594–608.

17. Maggi P, Sati P, Massacesi L. Magnetic resonance imaging of experimental autoimmune encephalomyelitis in the common marmoset. Journal of Neuroimmunology. 2017 Mar 15;304:86–92.

18. Donadieu M, Kelly H, Szczupak D, Lin JP, Song Y, Yen CCC, et al. Ultrahigh-resolution MRI Reveals Extensive Cortical Demyelination in a Nonhuman Primate Model of Multiple Sclerosis. Cereb Cortex. 2020 Sep 8;31(1):439–47.

19. Ventura-Antunes L, Mota B, Herculano-Houzel S. Different scaling of white matter volume, cortical connectivity, and gyrification across rodent and primate brains. Front Neuroanat. 2013 Apr 9;7:3.

20. Freund J, Stern ER, Pisani TM. Isoallergic encephalomyelitis and radiculitis in guinea pigs after one injection of brain and Mycobacteria in water-in-oil emulsion. J Immunol. 1947 Oct;57(2):179–94.

21. Billiau A, Matthys P. Modes of action of Freund’s adjuvants in experimental models of autoimmune diseases. Journal of Leukocyte Biology. 2001;70(6):849–60.

22. ‘t Hart BA, Hintzen RQ, Laman JD. Multiple sclerosis – a response-to-damage model. Trends in Molecular Medicine. 2009 Jun 1;15(6):235–44.

23. Jagessar SA, Kap YS, Heijmans N, van Driel N, van Straalen L, Bajramovic JJ, et al. Induction of progressive demyelinating autoimmune encephalomyelitis in common marmoset monkeys using MOG34-56 peptide in incomplete freund adjuvant. Journal of neuropathology and experimental neurology. 2010 Apr;69(4):372–85.

24. Charil A, Zijdenbos AP, Taylor J, Boelman C, Worsley KJ, Evans AC, et al. Statistical mapping analysis of lesion location and neurological disability in multiple sclerosis: application to 452 patient data sets. NeuroImage. 2003 Jul 1;19(3):532–44.

25. Pongratz V, Bussas M, Schmidt P, Grahl S, Gasperi C, El Husseini M, et al. Lesion location across diagnostic regions in multiple sclerosis. NeuroImage: Clinical. 2023 Jan 1;37:103311.

26. Lee NJ, Ha SK, Sati P, Absinta M, Luciano NJ, Lefeuvre JA, et al. Spatiotemporal distribution of fibrinogen in marmoset and human inflammatory demyelination. Brain. 2018 Jun 1;141(6):1637–49.

27. Hawkins CP, Mackenzie F, Tofts P, du Boulay EP, McDonald WI. Patterns of blood-brain barrier breakdown in inflammatory demyelination. Brain. 1991 Apr;114 (Pt 2):801–10.

28. Absinta M, Nair G, Monaco MCG, Maric D, Lee NJ, Ha SK, et al. THE “CENTRAL VEIN SIGN” IN INFLAMMATORY DEMYELINATION: THE ROLE OF FIBRILLAR COLLAGEN TYPE I. Ann Neurol. 2019 Jun;85(6):934–42.

29. Lee NJ, Ha SK, Sati P, Absinta M, Nair G, Luciano NJ, et al. Potential role of iron in repair of inflammatory demyelinating lesions. J Clin Invest. 2019 Oct 1;129(10):4365–76.

30. Villoslada P, Hauser SL, Bartke I, Unger J, Heald N, Rosenberg D, et al. Human nerve growth factor protects common marmosets against autoimmune encephalomyelitis by switching the balance of T helper cell type 1 and 2 cytokines within the central nervous system. J Exp Med. 2000 May 15;191(10):1799–806.

31. McDonald WI, Barnes D. The ocular manifestations of multiple sclerosis. 1. Abnormalities of the afferent visual system. Journal of Neurology, Neurosurgery & Psychiatry. 1992 Sep 1;55(9):747–52.

32. Frohman EM, Frohman TC, Zee DS, McColl R, Galetta S. The neuro-ophthalmology of multiple sclerosis. The Lancet Neurology. 2005 Feb 1;4(2):111–21.

33. Kolappan M, Henderson APD, Jenkins TM, Wheeler-Kingshott CAM, Plant GT, Thompson AJ, et al. Assessing structure and function of the afferent visual pathway in multiple sclerosis and associated optic neuritis. J Neurol. 2009 Mar;256(3):305–19.

34. Sotirchos ES, Gonzalez Caldito N, Filippatou A, Fitzgerald KC, Murphy OC, Lambe J, et al. Progressive Multiple Sclerosis Is Associated with Faster and Specific Retinal Layer Atrophy. Ann Neurol. 2020 Jun;87(6):885–96.

35. Lublin FD, Reingold SC, Sclerosis* NMSS (USA) AC on CT of NA in M. Defining the clinical course of multiple sclerosis: Results of an international survey. Neurology. 1996 Apr 1;46(4):907–11.

36. Lublin FD, Reingold SC, Cohen JA, Cutter GR, Sørensen PS, Thompson AJ, et al. Defining the clinical course of multiple sclerosis: The 2013 revisions. Neurology. 2014 Jul 15;83(3):278–86.

37. Brenner M, Messing A. Regulation of GFAP Expression. ASN Neuro. 2021 Jan 1;13:1759091420981206.

38. Khan A, Molitor A, Mayeur S, Zhang G, Rinaldi B, Lannes B, et al. A Homozygous Missense Variant in PPP1R1B/DARPP-32 Is Associated With Generalized Complex Dystonia. Mov Disord. 2022 Feb;37(2):365–74.

39. Guerra San Juan I, Nash LA, Smith KS, Leyton-Jaimes MF, Qian M, Klim JR, et al. Loss of mouse Stmn2 function causes motor neuropathy. Neuron. 2022 May 18;110(10):1671–1688.e6.

40. Jahn O, Tenzer S, Werner HB. Myelin Proteomics: Molecular Anatomy of an Insulating Sheath. Mol Neurobiol. 2009 Aug 1;40(1):55–72.

41. Cuzner ML, Hayes GM, Newcombe J, Woodroofe MN. The nature of inflammatory components during demyelination in multiple sclerosis. J Neuroimmunol. 1988 Dec;20(2–3):203–9.

42. Lin JP, Kelly HM, Song Y, Kawaguchi R, Geschwind DH, Jacobson S, et al. Transcriptomic architecture of nuclei in the marmoset CNS. Nat Commun. 2022 Sep 21;13(1):5531.

43. Jiang C, Qiu W, Yang Y, Huang H, Dai ZM, Yang A, et al. ADAMTS4 Enhances Oligodendrocyte Differentiation and Remyelination by Cleaving NG2 Proteoglycan and Attenuating PDGFRα Signaling. J Neurosci. 2023 Jun 14;43(24):4405–17.

44. Fard MK, van der Meer F, Sánchez P, Cantuti-Castelvetri L, Mandad S, Jäkel S, et al. BCAS1 expression defines a population of early myelinating oligodendrocytes in multiple sclerosis lesions. Sci Transl Med. 2017 Dec 6;9(419):eaam7816.

45. Scarisbrick IA, Blaber SI, Lucchinetti CF, Genain CP, Blaber M, Rodriguez M. Activity of a newly identified serine protease in CNS demyelination. Brain. 2002 Jun;125(Pt 6):1283–96.

46. Harroch S, Furtado GC, Brueck W, Rosenbluth J, Lafaille J, Chao M, et al. A critical role for the protein tyrosine phosphatase receptor type Z in functional recovery from demyelinating lesions. Nat Genet. 2002 Nov;32(3):411–4.

47. Becker I, Wang-Eckhardt L, Yaghootfam A, Gieselmann V, Eckhardt M. Differential expression of (dihydro)ceramide synthases in mouse brain: oligodendrocyte-specific expression of CerS2/Lass2. Histochem Cell Biol. 2008 Feb;129(2):233–41.

48. Wang X, Ge X, Qin Y, Liu D, Chen C. Ifi30 Is Required for Sprouting Angiogenesis During Caudal Vein Plexus Formation in Zebrafish. Front Physiol. 2022;13:919579.

49. Sun CK, Leu S, Sheu JJ, Tsai TH, Sung HC, Chen YL, et al. Paradoxical impairment of angiogenesis, endothelial function and circulating number of endothelial progenitor cells in DPP4-deficient rat after critical limb ischemia. Stem Cell Res Ther. 2013 Mar 21;4(2):31.

50. Gentile MT, Muto G, Lus G, Lövblad KO, Svenningsen ÅF, Colucci-D’Amato L. Angiogenesis and Multiple Sclerosis Pathogenesis: A Glance at New Pharmaceutical Approaches. J Clin Med. 2022 Aug 9;11(16):4643.

51. Liu X, Song C, Yang S, Ji Q, Chen F, Li W. IFI30 expression is an independent unfavourable prognostic factor in glioma. J Cell Mol Med. 2020 Nov;24(21):12433–43.

52. Ding L, Li LM, Hu B, Wang JL, Lu YB, Zhang RY, et al. TM4SF19 aggravates LPS-induced attenuation of vascular endothelial cell adherens junctions by suppressing VE-cadherin expression. Biochem Biophys Res Commun. 2020 Dec 17;533(4):1204–11.

53. Kasarello K, Mirowska-Guzel D. Anti-CD52 Therapy for Multiple Sclerosis: An Update in the COVID Era. Immunotargets Ther. 2021;10:237–46.

54. Chen X, Hou H, Qiao H, Fan H, Zhao T, Dong M. Identification of blood-derived candidate gene markers and a new 7-gene diagnostic model for multiple sclerosis. Biol Res. 2021 Apr 1;54(1):12.

55. Rang X, Liu Y, Wang J, Wang Y, Xu C, Fu J. Identification of multiple sclerosis-related genes regulated by EBV-encoded microRNAs in B cells. Multiple Sclerosis and Related Disorders. 2022 Mar 1;59:103563.

56. Bjornevik K, Cortese M, Healy BC, Kuhle J, Mina MJ, Leng Y, et al. Longitudinal analysis reveals high prevalence of Epstein-Barr virus associated with multiple sclerosis. Science. 2022 Jan 21;375(6578):296–301.

57. Hu W, Gauthier L, Baibakov B, Jimenez-Movilla M, Dean J. FIGLA, a basic helix-loop-helix transcription factor, balances sexually dimorphic gene expression in postnatal oocytes. Mol Cell Biol. 2010 Jul;30(14):3661–71.

58. Kunkle BW, Grenier-Boley B, Sims R, Bis JC, Damotte V, Naj AC, et al. Genetic meta-analysis of diagnosed Alzheimer’s disease identifies new risk loci and implicates Aβ, tau, immunity and lipid processing. Nat Genet. 2019 Mar;51(3):414–30.

59. Perrone F, Cacace R, van der Zee J, Van Broeckhoven C. Emerging genetic complexity and rare genetic variants in neurodegenerative brain diseases. Genome Med. 2021 Apr 14;13:59.

60. Smith AM, Davey K, Tsartsalis S, Khozoie C, Fancy N, Tang SS, et al. Diverse human astrocyte and microglial transcriptional responses to Alzheimer’s pathology. Acta Neuropathol. 2022 Jan;143(1):75–91.

61. Wang H, Devadoss D, Nair M, Chand HS, Lakshmana MK. Novel Alzheimer risk factor IQ motif containing protein K is abundantly expressed in the brain and is markedly increased in patients with Alzheimer’s disease. Front Cell Neurosci. 2022;16:954071.

62. Dai E, Zhang W, Cong D, Kang R, Wang J, Tang D. AIFM2 blocks ferroptosis independent of ubiquinol metabolism. Biochem Biophys Res Commun. 2020 Mar 19;523(4):966–71.

63. Dodson M, Anandhan A, Zhang DD. MGST1, a new soldier of NRF2 in the battle against ferroptotic death. Cell Chem Biol. 2021 Jun 17;28(6):741–2.

64. Lee J, Roh JL. SLC7A11 as a Gateway of Metabolic Perturbation and Ferroptosis Vulnerability in Cancer. Antioxidants. 2022 Dec;11(12):2444.

65. Liu J, Liu Y, Wang Y, Li C, Xie Y, Klionsky DJ, et al. TMEM164 is a new determinant of autophagy-dependent ferroptosis. Autophagy. 2023 Mar;19(3):945–56.

66. Hu J, Li G, Qu L, Li N, Liu W, Xia D, et al. TMEM166/EVA1A interacts with ATG16L1 and induces autophagosome formation and cell death. Cell Death Dis. 2016 Aug;7(8):e2323–e2323.

67. Zhao S, Wang H. EVA1A Plays an Important Role by Regulating Autophagy in Physiological and Pathological Processes. Int J Mol Sci. 2021 Jun 8;22(12):6181.

68. Shen S, Yang C, Liu X, Zheng J, Liu Y, Liu L, et al. RBFOX1 Regulates the Permeability of the Blood-Tumor Barrier via the LINC00673/MAFF Pathway. Mol Ther Oncolytics. 2020 Jun 26;17:138–52.

69. Opneja A, Kapoor S, Stavrou EX. Contribution of Platelets, the Coagulation and Fibrinolytic Systems to Cutaneous Wound Healing. Thromb Res. 2019 Jul;179:56–63.

70. Emre Y, Imhof BA. Matricellular protein CCN1/CYR61: a new player in inflammation and leukocyte trafficking. Semin Immunopathol. 2014 Mar;36(2):253–9.

71. Boraschi D, Tagliabue A. The interleukin-1 receptor family. Semin Immunol. 2013 Dec 15;25(6):394–407.

72. Stuard WL, Titone R, Robertson DM. The IGF/Insulin-IGFBP Axis in Corneal Development, Wound Healing, and Disease. Front Endocrinol (Lausanne). 2020;11:24.

73. Guilarte TR, Burton NC, Verina T, Prabhu VV, Becker KG, Syversen T, et al. Increased APLP1 expression and neurodegeneration in the frontal cortex of manganese-exposed non-human primates. J Neurochem. 2008 Jun;105(5):1948–59.

74. Galvan V, Chen S, Lu D, Logvinova A, Goldsmith P, Koo EH, et al. Caspase cleavage of members of the amyloid precursor family of proteins. J Neurochem. 2002 Jul;82(2):283–94.

75. Russell FA, King R, Smillie SJ, Kodji X, Brain SD. Calcitonin gene-related peptide: physiology and pathophysiology. Physiol Rev. 2014 Oct;94(4):1099–142.

76. Argunhan F, Brain SD. The Vascular-Dependent and -Independent Actions of Calcitonin Gene-Related Peptide in Cardiovascular Disease. Frontiers in Physiology [Internet]. 2022 [cited 2023 Aug 8];13. Available from: https://www.frontiersin.org/articles/10.3389/fphys.2022.833645

77. Schaefer L, Babelova A, Kiss E, Hausser HJ, Baliova M, Krzyzankova M, et al. The matrix component biglycan is proinflammatory and signals through Toll-like receptors 4 and 2 in macrophages. J Clin Invest. 2005 Aug;115(8):2223–33.

78. Choi DJ, An J, Jou I, Park SM, Joe EH. A Parkinson’s disease gene, DJ-1, regulates anti-inflammatory roles of astrocytes through prostaglandin D2 synthase expression. Neurobiol Dis. 2019 Jul;127:482–91.

79. Unno K, Konishi T. Preventive Effect of Soybean on Brain Aging and Amyloid-β Accumulation: Comprehensive Analysis of Brain Gene Expression. Recent Pat Food Nutr Agric. 2015;7(2):83–91.

80. Suzuki M, Tezuka K, Handa T, Sato R, Takeuchi H, Takao M, et al. Upregulation of ribosome complexes at the blood-brain barrier in Alzheimer’s disease patients. J Cereb Blood Flow Metab. 2022 Nov;42(11):2134–50.

81. Guo S, Chen Y, Xue X, Yang Y, Wang Y, Qiu S, et al. TRIB2 desensitizes ferroptosis via βTrCP-mediated TFRC ubiquitiantion in liver cancer cells. Cell Death Discov. 2021 Jul 27;7(1):196.

82. Wang C, JeBailey L, Ridgway ND. Oxysterol-binding-protein (OSBP)-related protein 4 binds 25-hydroxycholesterol and interacts with vimentin intermediate filaments. Biochem J. 2002 Feb 1;361(Pt 3):461– 72.

83. Muller WA. Getting Leukocytes to the Site of Inflammation. Vet Pathol. 2013 Jan;50(1):7–22.

84. Kivisäkk P, Mahad DJ, Callahan MK, Sikora K, Trebst C, Tucky B, et al. Expression of CCR7 in multiple sclerosis: Implications for CNS immunity. Annals of Neurology. 2004;55(5):627–38.

85. Piao JH, Wang Y, Duncan ID. CD44 is required for the migration of transplanted oligodendrocyte progenitor cells to focal inflammatory demyelinating lesions in the spinal cord. Glia. 2013 Mar;61(3):361–7.

86. Liu Y, Yang H, Liang C, Huang X, Deng X, Luo Z. Expression of functional thyroid-stimulating hormone receptor in microglia. Ann Endocrinol (Paris). 2022 Feb;83(1):40–5.

87. Qualai J, Li LX, Cantero J, Tarrats A, Fernández MA, Sumoy L, et al. Expression of CD11c Is Associated with Unconventional Activated T Cell Subsets with High Migratory Potential. PLoS One. 2016 Apr 27;11(4):e0154253.

88. Funk JL, Migliati E, Chen G, Wei H, Wilson J, Downey KJ, et al. Parathyroid hormone-related protein induction in focal stroke: a neuroprotective vascular peptide. Am J Physiol Regul Integr Comp Physiol. 2003 Apr;284(4):R1021–1030.

89. Kushnir MM, Peterson LK, Strathmann FG. Parathyroid hormone related protein concentration in human serum and CSF correlates with age. Clinical Biochemistry. 2018 Feb 1;52:56–60.

90. Yan H, Rivkees SA. Hepatocyte growth factor stimulates the proliferation and migration of oligodendrocyte precursor cells. J Neurosci Res. 2002 Sep 1;69(5):597–606.

91. Ohya W, Funakoshi H, Kurosawa T, Nakamura T. Hepatocyte growth factor (HGF) promotes oligodendrocyte progenitor cell proliferation and inhibits its differentiation during postnatal development in the rat. Brain Res. 2007 May 25;1147:51–65.

92. Tissir F, Goffinet AM. Expression of planar cell polarity genes during development of the mouse CNS. Eur J Neurosci. 2006 Feb;23(3):597–607.

93. Kawaguchi D, Furutachi S, Kawai H, Hozumi K, Gotoh Y. Dll1 maintains quiescence of adult neural stem cells and segregates asymmetrically during mitosis. Nat Commun. 2013 May 21;4(1):1880.

94. Grandbarbe L, Bouissac J, Rand M, Hrabé de Angelis M, Artavanis-Tsakonas S, Mohier E. Delta-Notch signaling controls the generation of neurons/glia from neural stem cells in a stepwise process. Development. 2003 Apr;130(7):1391–402.

95. Janghorban M, Xin L, Rosen JM, Zhang XHF. Notch Signaling as a Regulator of the Tumor Immune Response: To Target or Not To Target? Frontiers in Immunology [Internet]. 2018 [cited 2023 Aug 15];9. Available from: https://www.frontiersin.org/articles/10.3389/fimmu.2018.01649

96. Rivailler P, Cho YG, Wang F. Complete genomic sequence of an Epstein-Barr virus-related herpesvirus naturally infecting a new world primate: a defining point in the evolution of oncogenic lymphocryptoviruses. J Virol. 2002 Dec;76(23):12055–68.

97. Titus HE, Chen Y, Podojil JR, Robinson AP, Balabanov R, Popko B, et al. Pre-clinical and Clinical Implications of “Inside-Out” vs. “Outside-In” Paradigms in Multiple Sclerosis Etiopathogenesis. Front Cell Neurosci. 2020;14:599717.

98. Mapunda JA, Tibar H, Regragui W, Engelhardt B. How Does the Immune System Enter the Brain? Front Immunol. 2022;13:805657.

99. Tian L, Rauvala H, Gahmberg CG. Neuronal regulation of immune responses in the central nervous system. Trends in Immunology. 2009 Feb 1;30(2):91–9.

100. Molofsky AV, Slutsky SG, Joseph NM, He S, Pardal R, Krishnamurthy J, et al. Increasing p16INK4a expression decreases forebrain progenitors and neurogenesis during ageing. Nature. 2006 Sep 28;443(7110):448–52.

101. Zhang R, Zhang K. Mitochondrial NAD kinase in health and disease. Redox Biol. 2023 Apr;60:102613.

102. Port F, Basler K. Wnt Trafficking: New Insights into Wnt Maturation, Secretion and Spreading. Traffic. 2010;11(10):1265–71.

103. Harkins D, Cooper HM, Piper M. The role of lipids in ependymal development and the modulation of adult neural stem cell function during aging and disease. Seminars in Cell & Developmental Biology. 2021 Apr 1;112:61–8.

104. Field J, Browning SR, Johnson LJ, Danoy P, Varney MD, Tait BD, et al. A polymorphism in the HLA-DPB1 gene is associated with susceptibility to multiple sclerosis. PLoS One. 2010 Oct 26;5(10):e13454.

105. Wang Z, Sadovnick AD, Traboulsee AL, Ross JP, Bernales CQ, Encarnacion M, et al. Nuclear Receptor NR1H3 in Familial Multiple Sclerosis. Neuron. 2016 Jun 1;90(5):948–54.

106. Oldoni E, Smets I, Mallants K, Vandebergh M, Van Horebeek L, Poesen K, et al. CHIT1 at Diagnosis Reflects Long-Term Multiple Sclerosis Disease Activity. Ann Neurol. 2020 Apr;87(4):633–45.

107. Chen BJ, Mills JD, Takenaka K, Bliim N, Halliday GM, Janitz M. Characterization of circular RNAs landscape in multiple system atrophy brain. J Neurochem. 2016 Nov;139(3):485–96.

108. Li Y, Zheng Q, Bao C, Li S, Guo W, Zhao J, et al. Circular RNA is enriched and stable in exosomes: a promising biomarker for cancer diagnosis. Cell Res. 2015 Aug;25(8):981–4.

109. Coppé JP, Desprez PY, Krtolica A, Campisi J. The Senescence-Associated Secretory Phenotype: The Dark Side of Tumor Suppression. Annu Rev Pathol. 2010;5:99–118.

110. Chinta SJ, Woods G, Rane A, Demaria M, Campisi J, Andersen JK. Cellular senescence and the aging brain. Experimental Gerontology. 2015 Aug 1;68:3–7.

111. Sikora E, Bielak-Zmijewska A, Dudkowska M, Krzystyniak A, Mosieniak G, Wesierska M, et al. Cellular Senescence in Brain Aging. Frontiers in Aging Neuroscience [Internet]. 2021 [cited 2023 Aug 17];13. Available from: https://www.frontiersin.org/articles/10.3389/fnagi.2021.646924

112. Kang C, Elledge SJ. How autophagy both activates and inhibits cellular senescence. Autophagy. 2016 May 3;12(5):898–9.

113. Vidal R, Wagner JUG, Braeuning C, Fischer C, Patrick R, Tombor L, et al. Transcriptional heterogeneity of fibroblasts is a hallmark of the aging heart. JCI Insight. 2019 Nov 14;4(22):e131092, 131092.

114. Wu MX, Wang SH, Xie Y, Chen ZT, Guo Q, Yuan WL, et al. Interleukin-33 alleviates diabetic cardiomyopathy through regulation of endoplasmic reticulum stress and autophagy via insulin-like growth factor-binding protein 3. J Cell Physiol. 2021 Jun;236(6):4403–19.

115. Pandey R, Shukla P, Anjum B, Gupta HP, Pal S, Arjaria N, et al. Estrogen deficiency induces memory loss via altered hippocampal HB-EGF and autophagy. J Endocrinol. 2020 Jan 1;244(1):53–70.

116. Li ZL, Zhang HL, Huang Y, Huang JH, Sun P, Zhou NN, et al. Autophagy deficiency promotes triple-negative breast cancer resistance to T cell-mediated cytotoxicity by blocking tenascin-C degradation. Nat Commun. 2020 Jul 30;11(1):3806.

117. Su BC, Hsu PL, Mo FE. CCN1 triggers adaptive autophagy in cardiomyocytes to curb its apoptotic activities. J Cell Commun Signal. 2020 Mar;14(1):93–100.

118. Guo P, Ma Y, Deng G, Li L, Gong Y, Yang F, et al. CYR61, regulated by miR-22-3p and MALAT1, promotes autophagy in HK-2 cell inflammatory model. Transl Androl Urol. 2021 Aug;10(8):3486–500.

119. Jakovcevski I, Miljkovic D, Schachner M, Andjus PR. Tenascins and inflammation in disorders of the nervous system. Amino Acids. 2013 Apr;44(4):1115–27.

120. Fujita M, Sasada M, Eguchi M, Iyoda T, Okuyama S, Osawa T, et al. Induction of cellular senescence in fibroblasts through β1-integrin activation by tenascin-C-derived peptide and its protumor effect. Am J Cancer Res. 2021 Sep 15;11(9):4364–79.

121. Oyagi A, Hara H. Essential roles of heparin-binding epidermal growth factor-like growth factor in the brain. CNS Neurosci Ther. 2012 Oct;18(10):803–10.

122. Hoeflich A, Fitzner B, Walz C, Hecker M, Tuchscherer A, Brenmoehl J, et al. Reduced Fragmentation of IGFBP-2 and IGFBP-3 as a Potential Mechanism for Decreased Ratio of IGF-II to IGFBPs in Cerebrospinal Fluid in Response to Repeated Intrathecal Administration of Triamcinolone Acetonide in Patients With Multiple Sclerosis. Frontiers in Endocrinology [Internet]. 2021 [cited 2022 Jul 15];11. Available from: https://www.frontiersin.org/articles/10.3389/fendo.2020.565557

123. Elzi DJ, Lai Y, Song M, Hakala K, Weintraub ST, Shiio Y. Plasminogen activator inhibitor 1--insulin-like growth factor binding protein 3 cascade regulates stress-induced senescence. Proc Natl Acad Sci U S A. 2012 Jul 24;109(30):12052–7.

124. Chesik D, De Keyser J, Wilczak N. Insulin-like growth factor binding protein-2 as a regulator of IGF actions in CNS: Implications in multiple sclerosis. Cytokine & Growth Factor Reviews. 2007 Jun 1;18(3):267–78.

125. Lanzillo R, Di Somma C, Quarantelli M, Ventrella G, Gasperi M, Prinster A, et al. Insulin-like growth factor (IGF)-I and IGF-binding protein-3 serum levels in relapsing-remitting and secondary progressive multiple sclerosis patients. Eur J Neurol. 2011 Dec;18(12):1402–6.

126. Lu H, Wu PF, Ma DL, Zhang W, Sun M. Growth Factors and Their Roles in Multiple Sclerosis Risk. Front Immunol. 2021;12:768682.

127. Akcali A, Bal B, Erbagci B. Circulating IGF-1, IGFB-3, GH and TSH levels in multiple sclerosis and their relationship with treatment. Neurological Research. 2017 Jul 3;39(7):606–11.

128. Varma Shrivastav S, Bhardwaj A, Pathak KA, Shrivastav A. Insulin-Like Growth Factor Binding Protein-3 (IGFBP-3): Unraveling the Role in Mediating IGF-Independent Effects Within the Cell. Frontiers in Cell and Developmental Biology [Internet]. 2020 [cited 2023 Aug 18];8. Available from: https://www.frontiersin.org/articles/10.3389/fcell.2020.00286

129. Absinta M, Maric D, Gharagozloo M, Garton T, Smith MD, Jin J, et al. A lymphocyte–microglia–astrocyte axis in chronic active multiple sclerosis. Nature. 2021 Sep 30;597(7878):709–14.

130. Hong DE, Yu JE, Yoo SS, Yeo IJ, Son DJ, Yun J, et al. CHI3L1 induces autophagy through the JNK pathway in lung cancer cells. Sci Rep. 2023 Jun 20;13(1):9964.

131. Shen X, Kan S, Liu Z, Lu G, Zhang X, Chen Y, et al. EVA1A inhibits GBM cell proliferation by inducing autophagy and apoptosis. Exp Cell Res. 2017 Mar 1;352(1):130–8.

132. Zhang T, Bhambri A, Zhang Y, Barbosa D, Bae HG, Xue J, et al. Autophagy collaborates with apoptosis pathways to control oligodendrocyte number. Cell Reports. 2023 Aug 29;42(8):112943.

133. Thorin-Trescases N, Labbé P, Mury P, Lambert M, Thorin E. Angptl2 is a Marker of Cellular Senescence: The Physiological and Pathophysiological Impact of Angptl2-Related Senescence. Int J Mol Sci. 2021 Nov 12;22(22):12232.

134. Huang H, Ni H, Ma K, Zou J. ANGPTL2 regulates autophagy through the MEK/ERK/Nrf-1 pathway and affects the progression of renal fibrosis in diabetic nephropathy. Am J Transl Res. 2019;11(9):5472–86.

135. Bankston AN, Forston MD, Howard RM, Andres KR, Smith AE, Ohri SS, et al. Autophagy is essential for oligodendrocyte differentiation, survival, and proper myelination. Glia. 2019;67(9):1745–59.

136. Belgrad J, Pace RD, Fields RD. Autophagy in Myelinating Glia. J Neurosci. 2020 Jan 8;40(2):256–66.

137. Misrielal C, Mauthe M, Reggiori F, Eggen BJL. Autophagy in Multiple Sclerosis: Two Sides of the Same Coin. Frontiers in Cellular Neuroscience [Internet]. 2020 [cited 2022 Jun 28];14. Available from: https://www.frontiersin.org/article/10.3389/fncel.2020.603710

138. Marques JP, Kober T, Krueger G, van der Zwaag W, Van de Moortele PF, Gruetter R. MP2RAGE, a self bias-field corrected sequence for improved segmentation and T1-mapping at high field. Neuroimage. 2010 Jan 15;49(2):1271–81.

139. Luciano NJ, Sati P, Nair G, Guy JR, Ha SK, Absinta M, et al. Utilizing 3D printing technology to merge MRI with histology: A protocol for brain sectioning. Journal of visualized experiments : JoVE. 2016 Dec;(118):e54780–e54780.

140. Liu C, Ye FQ, Yen CCC, Newman JD, Glen D, Leopold DA, et al. A digital 3D atlas of the marmoset brain based on multi-modal MRI. Neuroimage. 2018 Apr 1;169:106–16.

141. Liu C, Ye FQ, Newman JD, Szczupak D, Tian X, Yen CCC, et al. A resource for the detailed 3D mapping of white matter pathways in the marmoset brain. Nat Neurosci. 2020 Feb;23(2):271–80.

142. McGinnis CS, Murrow LM, Gartner ZJ. DoubletFinder: Doublet detection in single-cell RNA sequencing data using artificial nearest neighbors. Cell systems. 2019 Apr;8(4):329–337.e4.

143. Young MD, Behjati S. SoupX removes ambient RNA contamination from droplet based single-cell RNA sequencing data. bioRxiv. 2020 Feb;12(1):303727.

144. Korsunsky I, Millard N, Fan J, Slowikowski K, Zhang F, Wei K, et al. Fast, sensitive and accurate integration of single-cell data with Harmony. Nature Methods. 2019 Dec;16(12):1289–96.

145. Jin S, Guerrero-Juarez CF, Zhang L, Chang I, Ramos R, Kuan CH, et al. Inference and analysis of cell-cell communication using CellChat. Nat Commun. 2021 Feb 17;12(1):1088.

146. Stuart T, Butler A, Hoffman P, Hafemeister C, Papalexi E, Mauck WM, et al. Comprehensive Integration of Single-Cell Data. Cell. 2019 Jun 13;177(7):1888–1902.e21.

147. Hafemeister C, Satija R. Normalization and variance stabilization of single-cell RNA-seq data using regularized negative binomial regression. Genome Biol. 2019 Dec 23;20(1):296.

148. Cao J, Spielmann M, Qiu X, Huang X, Ibrahim DM, Hill AJ, et al. The single-cell transcriptional landscape of mammalian organogenesis. Nature. 2019 Feb;566(7745):496–502.

149. Tustison NJ, Avants BB, Cook PA, Zheng Y, Egan A, Yushkevich PA, et al. N4ITK: improved N3 bias correction. IEEE Trans Med Imaging. 2010 Jun;29(6):1310–20.

